# Identification of biomarkers and pathways for focal segmental glomerulosclerosis using next generation sequencing data and bioinformatics analysis

**DOI:** 10.1101/2022.06.08.495272

**Authors:** Basavaraj Vastrad, Chanabasayya Vastrad

## Abstract

Focal segmental glomerulosclerosis (FSGS) is a renal disease leading threat to human health around the world. Here we aimed to explore novel molecular and potential therapeutic targets in FSGS through adopting integrated bioinformatics tools. Next generation sequencing (NGS) data of GSE197307 was available from Gene Expression Omnibus (GEO) database. Furthermore, differentially expressed genes (DEGs) were screened using the DESeq2 package in R software. Gene ontology (GO) and REACTOME pathway enrichment analyses of DEGs were performed via g:Profiler. Then, the protein-protein interaction (PPI) network, miRNA-hub gene regulatory network and TF-hub gene regulatory network were constructed using the HIPPIE, miRNet and NetworkAnalyst databases. Hub genes were validated using the receiver operating characteristic curve (ROC) analysis. By performing DEGs analysis, 488 up regulated genes and 488 down regulated genes were successfully identified from GSE197307, respectively. And they were mainly enriched in the terms of multicellular organismal process, cell periphery, protein binding, metabolism, immune system process, signaling receptor binding and immune system. Based on the data of protein–protein interaction (PPI), miRNA-hub gene regulatory network and TF-hub gene regulatory network, the top hub genes were ranked, including ILK, DDX5, MATR3, ALB, FOS, MKI67, PLK1, TNF, LCK and GTSE1. Bioinformatics analysis of NGS data identified signaling pathways and hub genes, potentially representing molecular mechanisms for the occurrence, progression, and risk prediction in FSGS.

## Introduction

Focal segmental glomerulosclerosis (FSGS) is a glomerular injury characterized by marked proteinuria and podocyte injury [1]. The number of cases of FSGS is rising worldwide and it has become a key health concern. The complications of FSGS include diabetes mellitus [2], sickle cell disease [3], hypertension [4], cardiovascular diseases [5], polycystic ovarian syndrome (PCOS) [6], obesity [7] and multiple sclerosis [8],which might reduce the life expectancy. Patients with FSGS always pose considerable psychological, financial, and physical burdens. Therefore, it is necessary to improve our understanding of FSGS pathogenesis and to develop better screening methods for FSGS.

Thus, because of such serious implications, researchers are focusing their attention on investigating the molecular pathogenesis and mechanisms underlying FSGS. Several investigations have focused on genes and signaling pathways in the pathogenesis of FSGS. Genes include APOL1 [9], INF2 [10], COL4A [11], MYO1E [12] and COL4A3 [13] are involved in the occurrence and development of FSGS. Signaling pathways include TGF-beta/Smad signaling pathway [14], JAK-STAT signaling pathway [15], Nrf2 signaling pathway [16], notch signaling pathway [17] and C3aR/C5aR-sphingosine kinase 1 pathway [18] plays a critical role in the pathophysiology of FSGS. Despite the increasing insights into several aspects of these mechanisms, a comprehensive overview of the integrated biological landscape underlying FSGS.

Recently, next generation sequencing (NGS) of human disease samples have generated massive bioinformatics data, which facilitated understanding the molecular mechanisms involved in the related biological process [19]. A high-quality NGS data could potentially link molecular biomarkers to the advancement, diagnosis, prognosis and treatment of FSGS.

In this investigation, we identified DEGs in NGS data GSE197307 from the National Centre of Biotechnology Information (NCBI) Gene Expression Omnibus database (GEO, https://www.ncbi.nlm.nih.gov/geo/) [20]. We performed Gene Ontology (GO) and REACTOME pathway enrichment analyses and constructed a protein–protein interaction (PPI) network, modules, miRNA-hub gene regulatory network and TF-hub gene regulatory network. Validations of hub genes were determined by receiver operating characteristic curve (ROC) analysis. Hub genes and the clinical prognosis of patients with FSGS were compared and analyzed with the objective of identifying novel prognostic and diagnostic biomarkers, and therapeutic targets.

## Materials and Methods

### Next generation sequencing data source

The gene expression profile data of GSE197307, downloaded from GEO database was used to screen DEGs in FSGS. Total 101 samples (93 FSGS samples and 8 normal control samples) were included in the data series of GSE197307. The NGS data from GSE197307 was based on GPL18573 Illumina NextSeq 500 (Homo sapiens).

### Identification of DEGs

DESeq2 package of R software [21] was used to identify DEGs between FSGS and normal control. The adjusted P-value and [logFC] were determined. The Benjamini & Hochberg false discovery rate method was used as a correction factor for the adjusted P-value [22]. The statistically significant DEGs were identified according to adjusted P<0.05, [log FC] > 0.97 for up regulated genes and [log FC] < −2.15 for down regulated genes. We also used an ggplot2 and gplot in R software to volcano plot and heat map of the DEGs of NGS dataset.

### GO and pathway enrichment analyses of DEGs

g:Profiler (http://biit.cs.ut.ee/gprofiler/) [23] is an open database that integrates biological data and analytical tools for functional annotation of genes and pathways21. GO (http://www.geneontology.org) [24] is a bioinformatics tool for annotating genes and analyzing the biological processes they are associated in. REACTOME (https://reactome.org/) [25] is a pathway database for analyzing relevant signaling pathways in large scale molecular datasets generated by high-throughput experimental techniques. g:Profiler was used for GO enrichment analysis of the DEGs in terms of the biological process (BP), cell composition (CC) and molecular function (MF) for each gene. REACTOME pathway enrichment analysis was performed to clarify the function of the DEGs and the cell signaling pathways. When P < 0.05, we believe that statistics are significantly different.

### Construction of the PPI network and module analysis

Human Integrated Protein-Protein Interaction rEference (HIPPIE) (http://cbdm-01.zdv.uni-mainz.de/~mschaefer/hippie/index.php) [26]. was used to construct a PPI network for the DEGs, which was visualized in 3.8.2 (http://www.cytoscape.org/) [27] (version 3.9.1). The Network Analyzer in Cytoscape was utilized to calculate node degree [28], betweenness [29], stress [30] and closeness [31]. PEWCC1 [32] was used to perform modular analysis, with the parameters set as follows: a Degree Cutoff=2, Node Score Cutoff=0.2, K-Core=2, and Max. Depth=100. Finally, pathway enrichment analyses of candidate genes in each module of PPI network were performed using g:Profiler.

### miRNA-hub gene regulatory network construction

miRNet database (https://www.mirnet.ca/) [33] is a bioinformatics platform for predicting miRNA-hub gene pairs. In the current investigation, the targets of the hub genes and miRNAs were predicted using fourteen databases: TarBase, miRTarBase, miRecords, miRanda (S mansoni only), miR2Disease, HMDD, PhenomiR, SM2miR, PharmacomiR, EpimiR, starBase, TransmiR, ADmiRE, and TAM 2. The screening criterion was that the miRNA target exists in the fourteen databases concurrently. The miRNA-hub gene regulatory network was depicted and visualized using Cytoscape software [27].

### TF-hub gene regulatory network construction

NetworkAnalyst database (https://www.networkanalyst.ca/) [34] is a bioinformatics platform for predicting TF-hub gene pairs. In the current investigation, the targets of the hub genes and TFs were predicted using Jasper database. The TF-hub gene regulatory network was depicted and visualized using Cytoscape software [27].

### Receiver operating characteristic curve (ROC) analysis

To evaluate the role of candidate genes in the diagnosis of FSGS, receiver operating characteristic (ROC) curve analysis was conducted by using pROC package [35]. The area under the ROC curve (AUC) value was utilized to determine the diagnostic effectiveness in discriminating FSGS from normal control samples in the GSE175759 dataset.

## Results

### Identification of DEGs

The NGS dataset was selected using the DESeq2 package (P-value <0.05, [log FC] > 0.97 for up regulated genes and [log FC] < −2.15 for down regulated genes) in R software. After a comprehensive analysis of the NGS dataset, we identified 976 DEGs, of which 488 genes were up regulated and 488 genes were down regulated and are presented in Table 1. The volcano plot presents the DEGs in the GSE197307 (Fig. 1). The DEGs are presented by a cluster heatmap in Fig. 2.

**Fig. 1.**
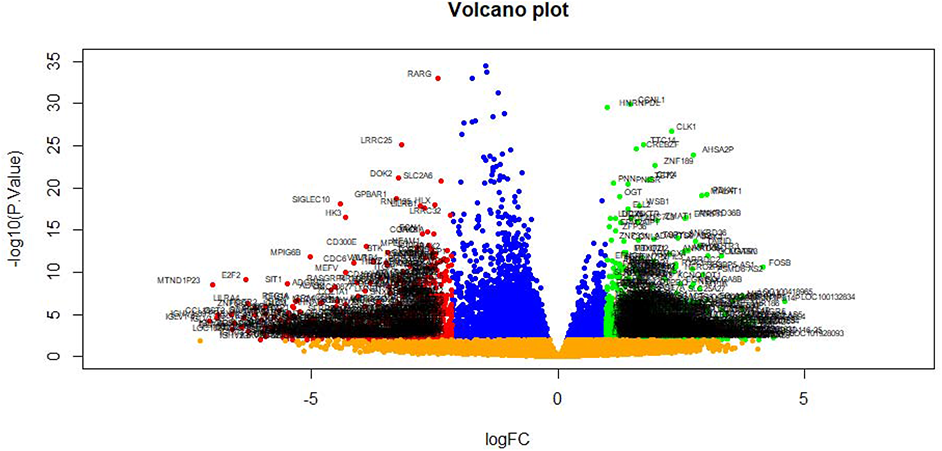
Volcano plot of differentially expressed genes. Genes with a significant change of more than two-fold were selected. Green dot represented up regulated significant genes and red dot represented down regulated significant genes.

**Fig. 2.**
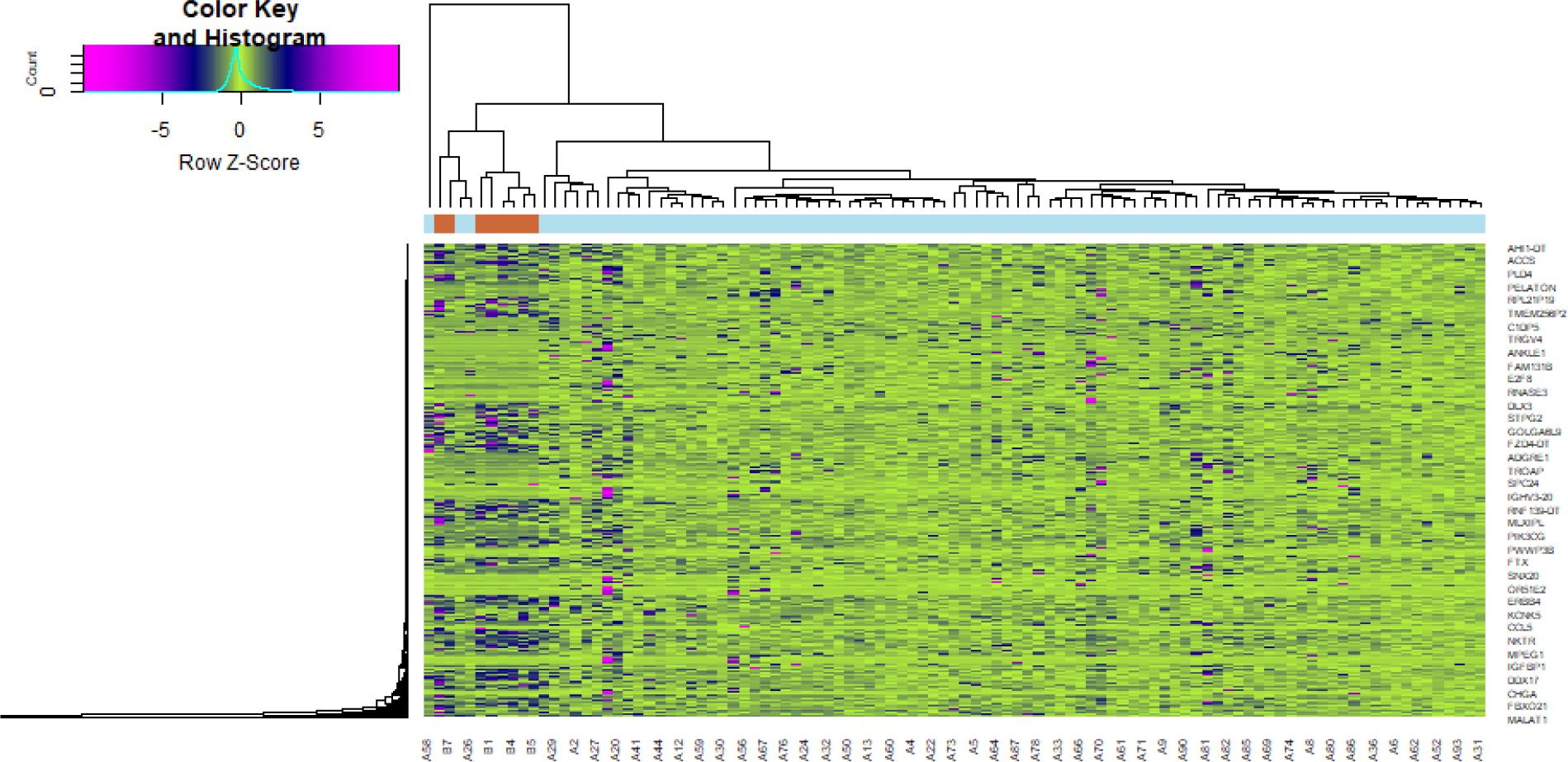
Heat map of differentially expressed genes. Legend on the top left indicate log fold change of genes. (A1 – A93 = FSGS samples; B1 – B8 = normal control samples)

**Table 1.**
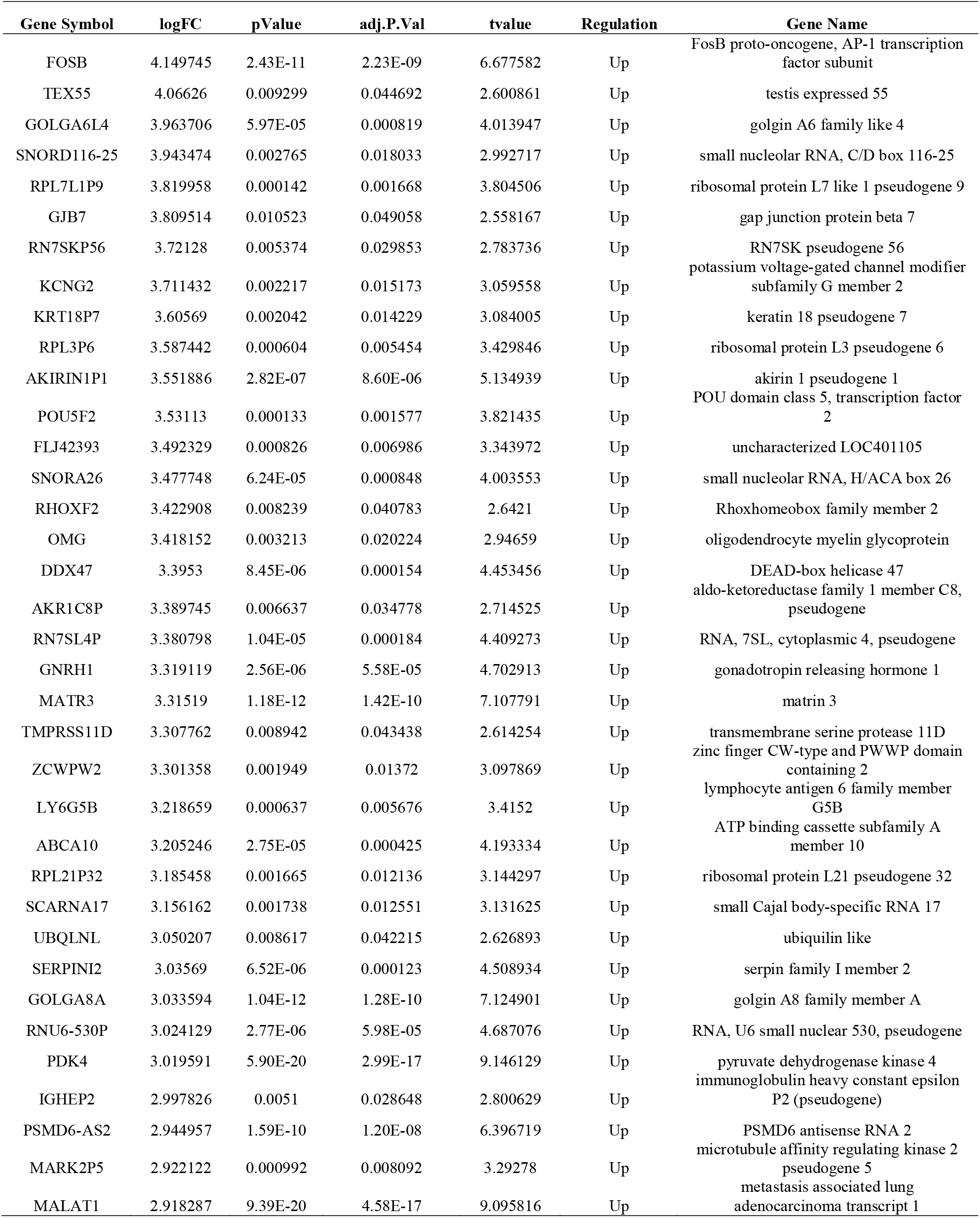
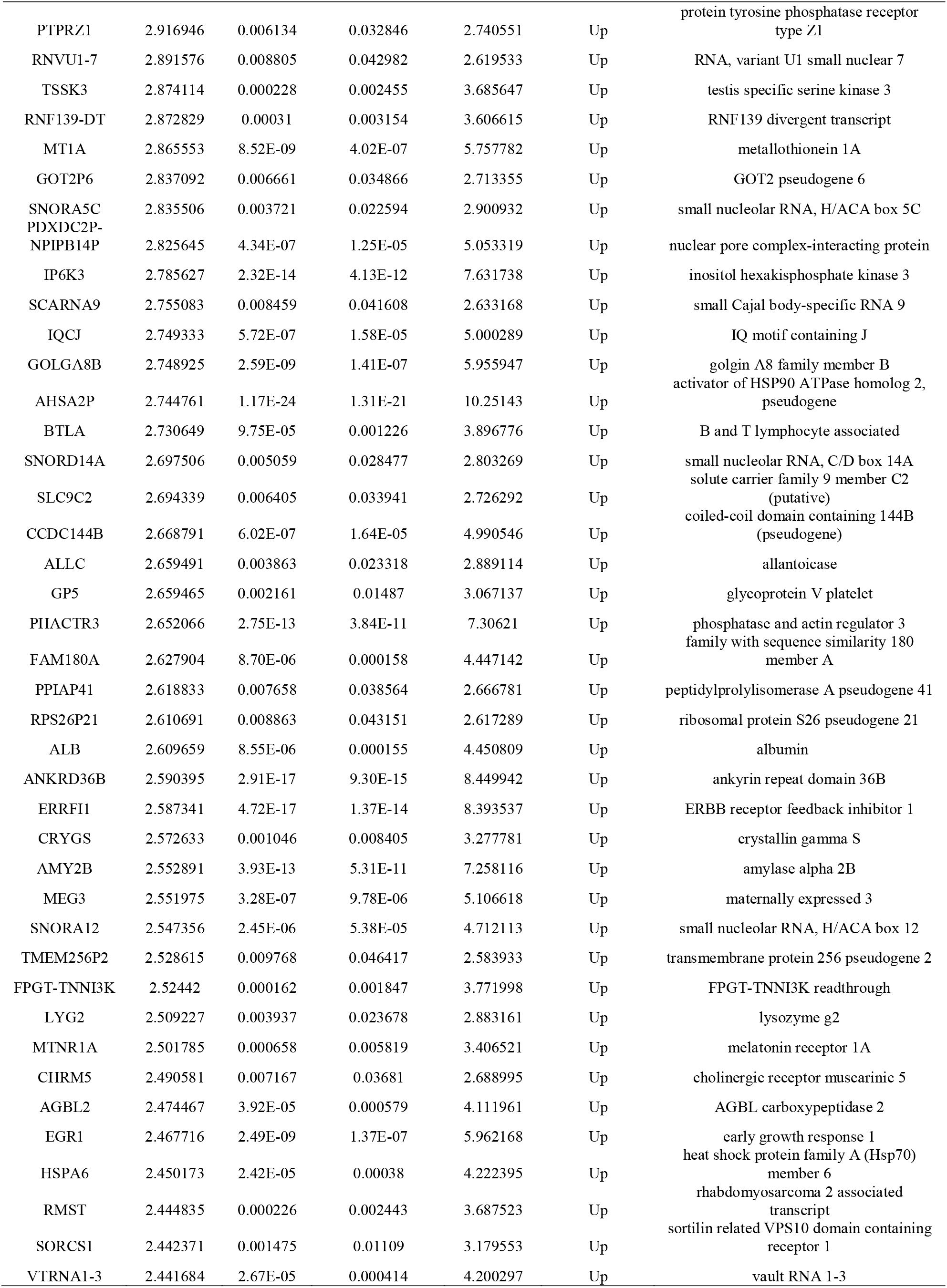
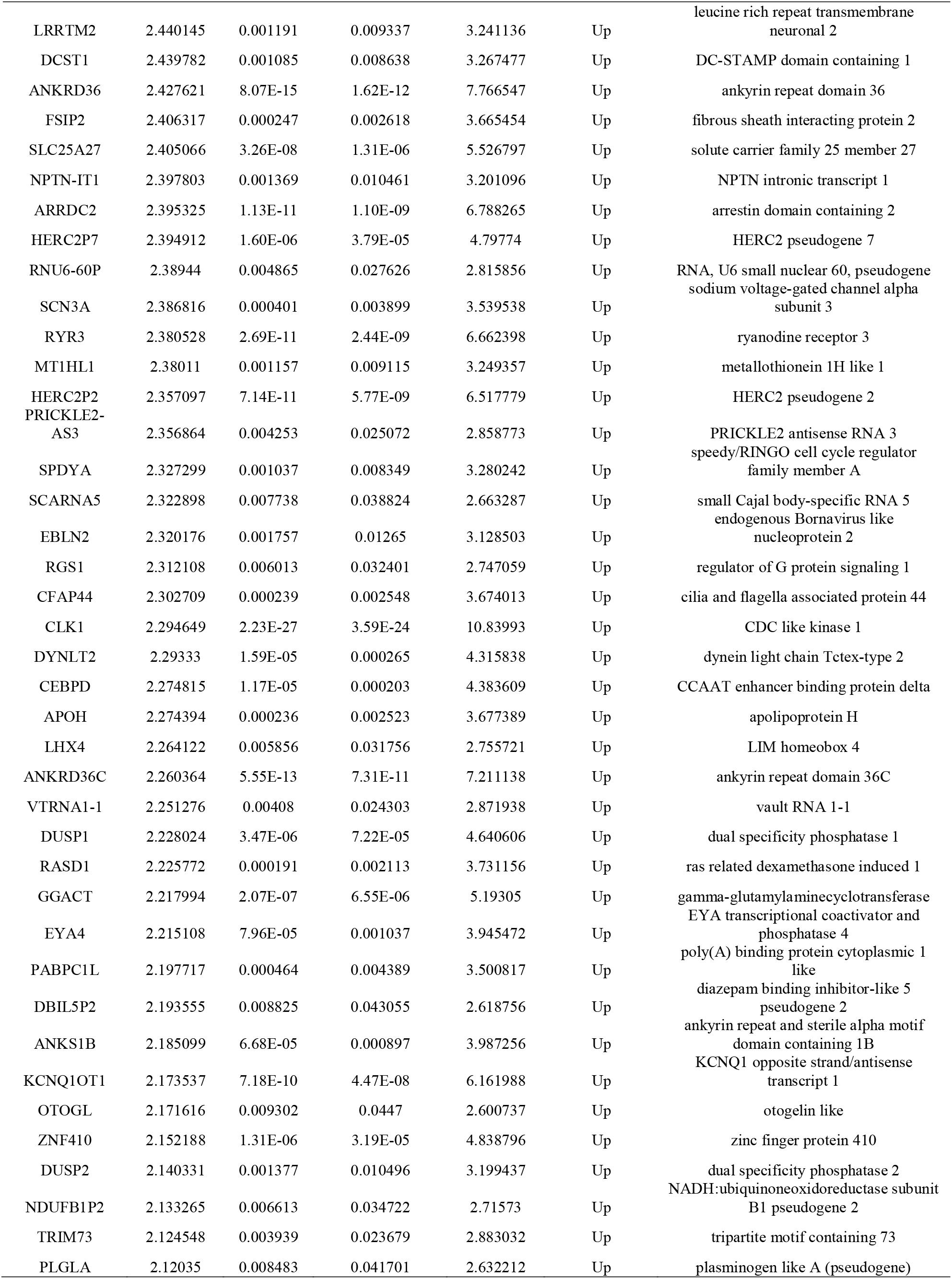
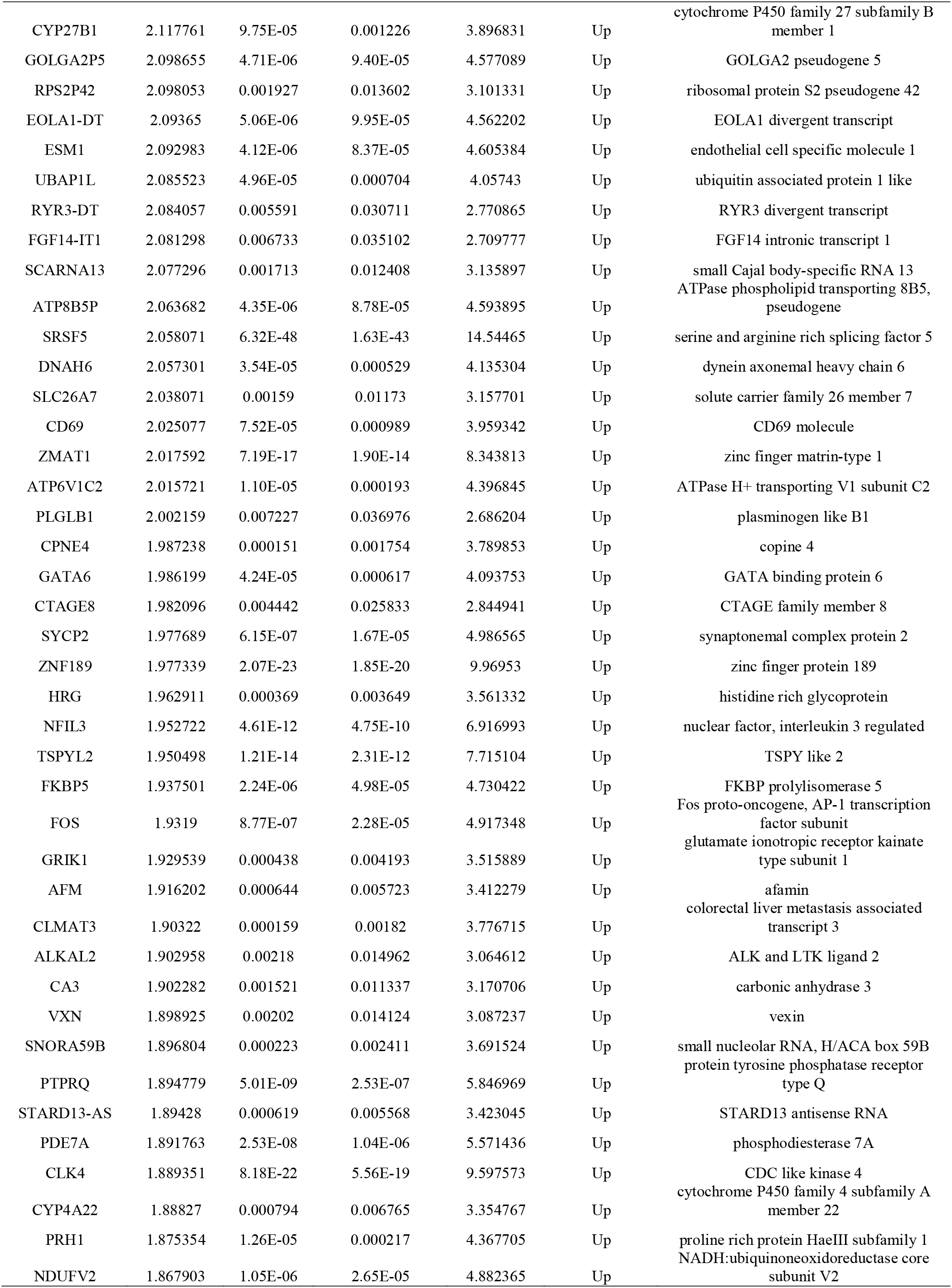
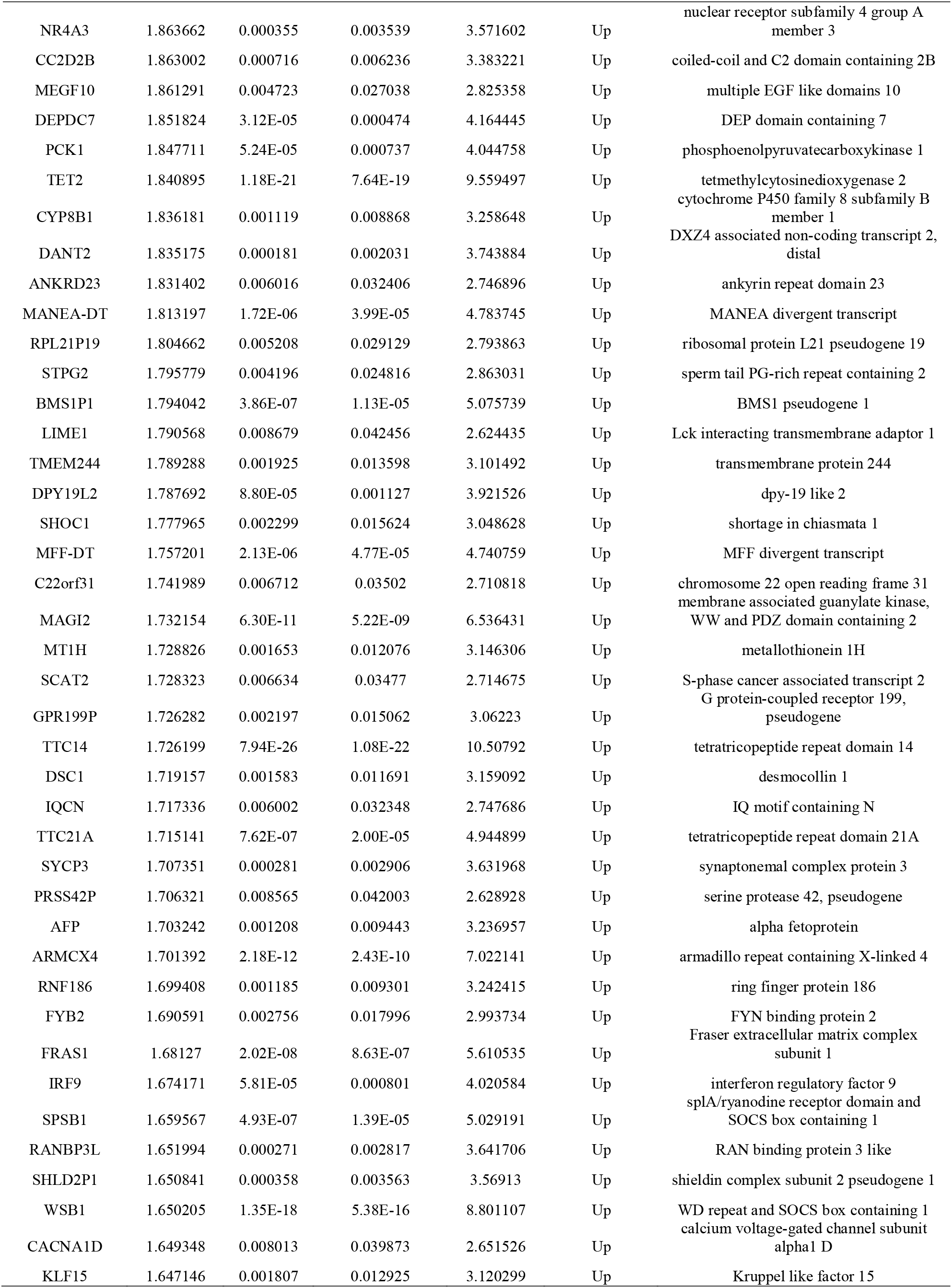
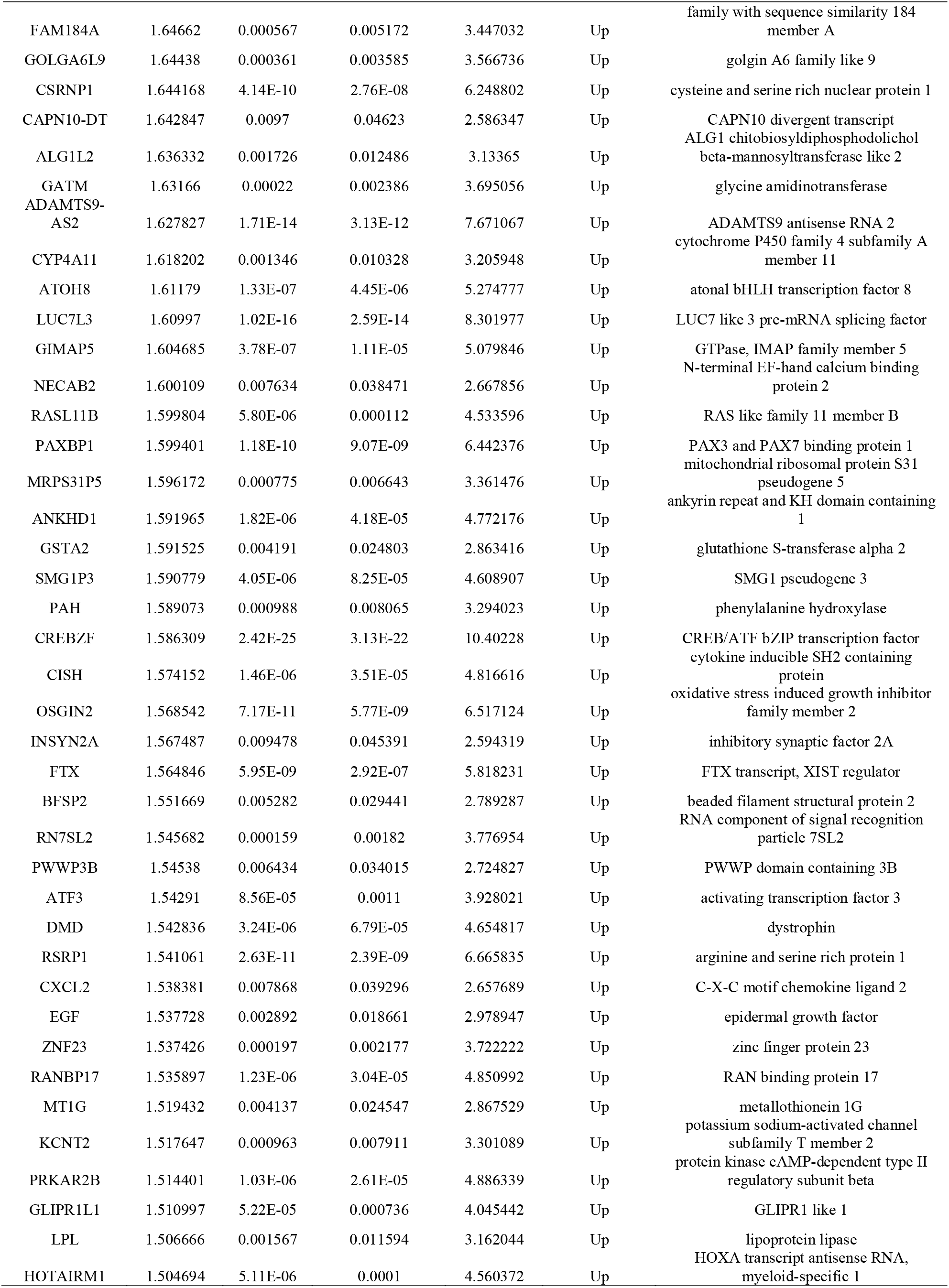
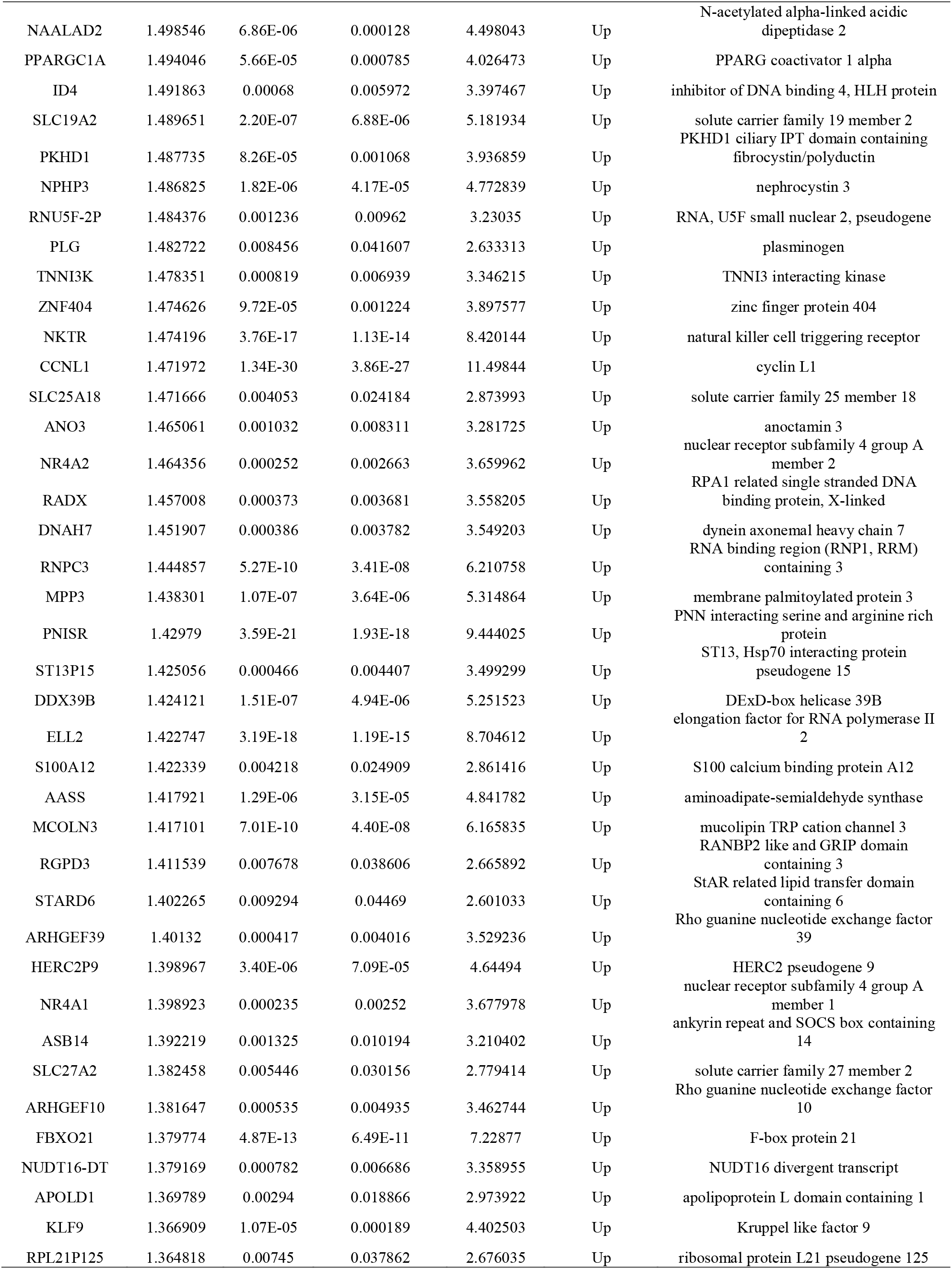
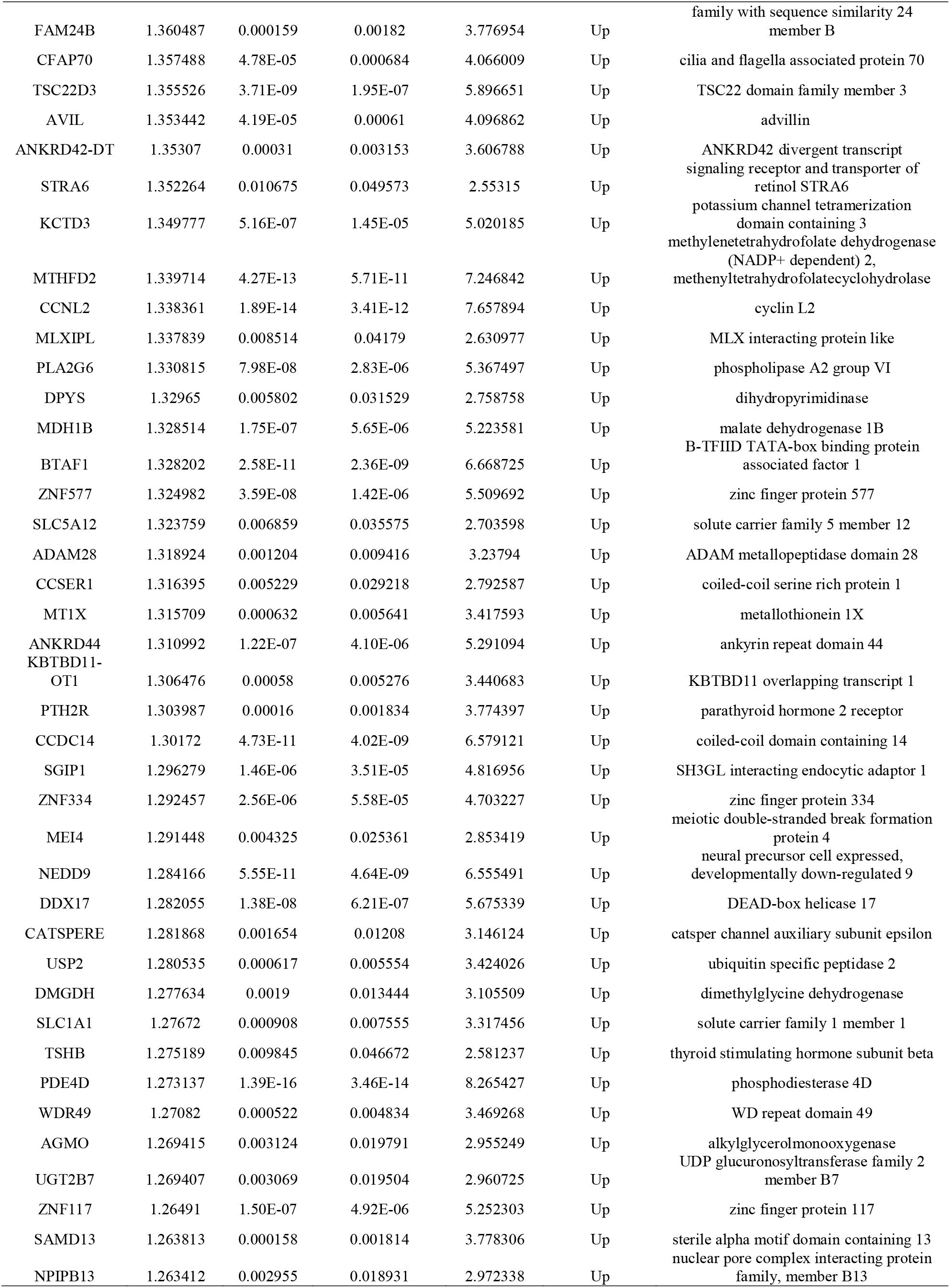
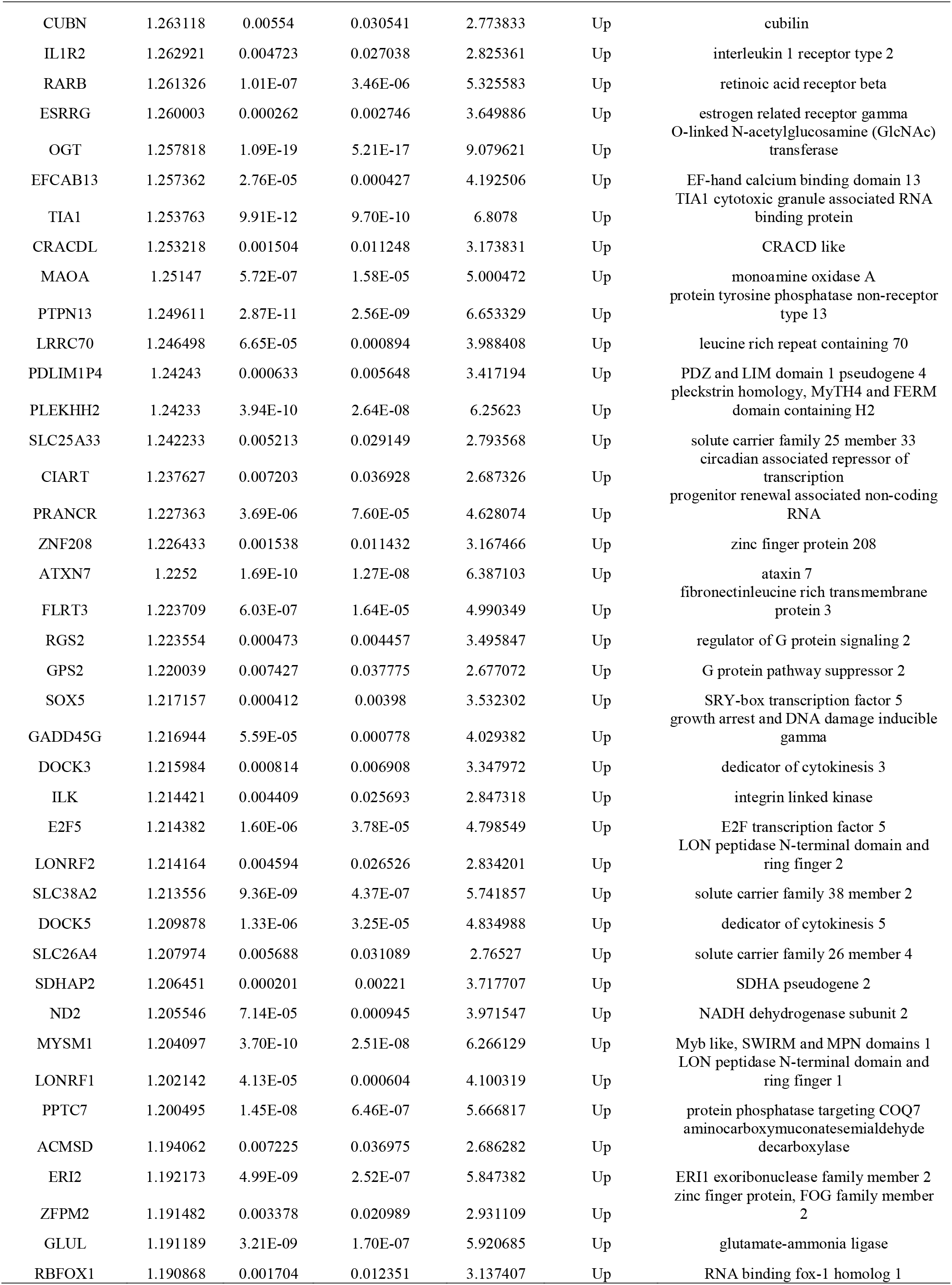
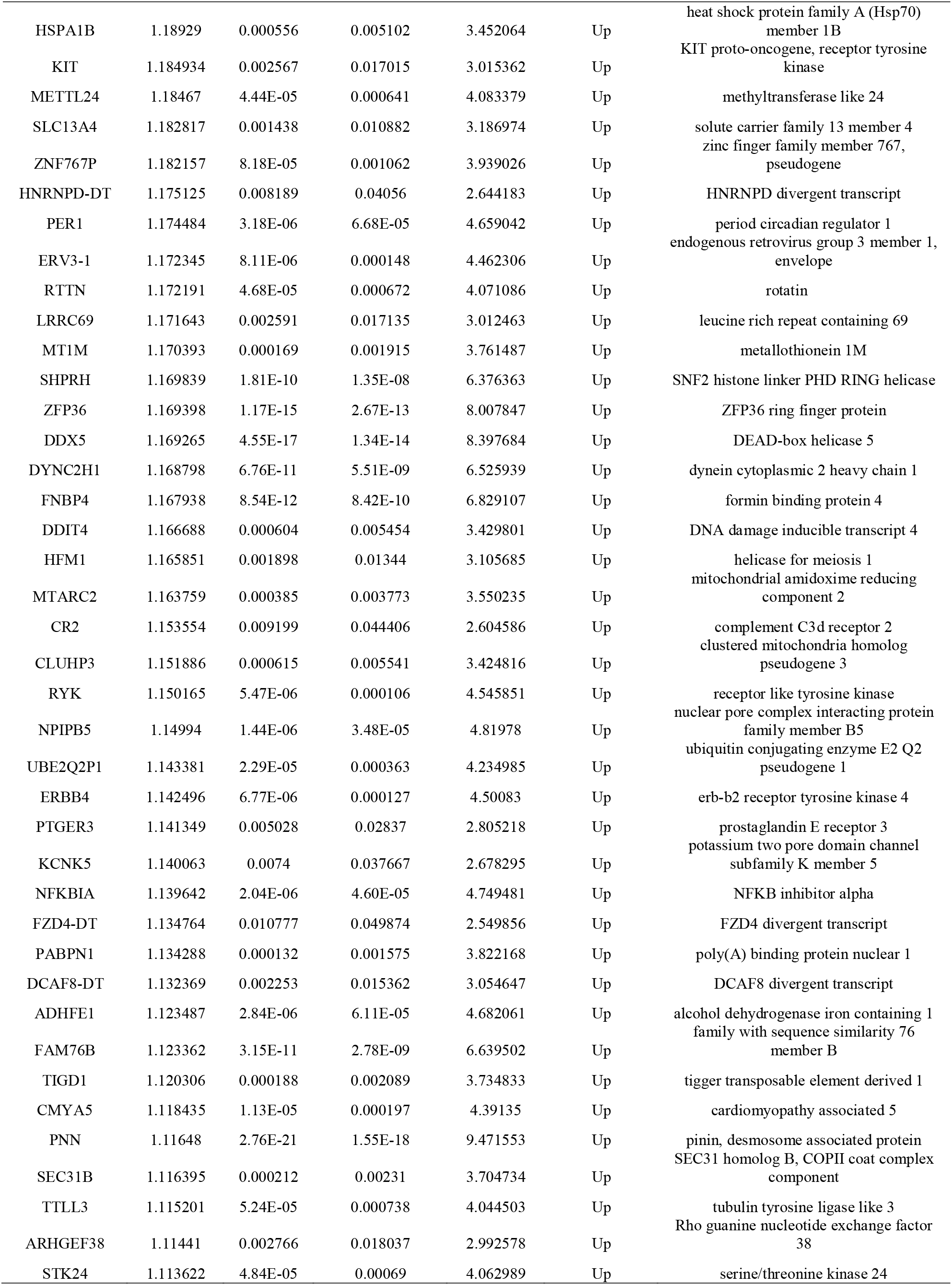
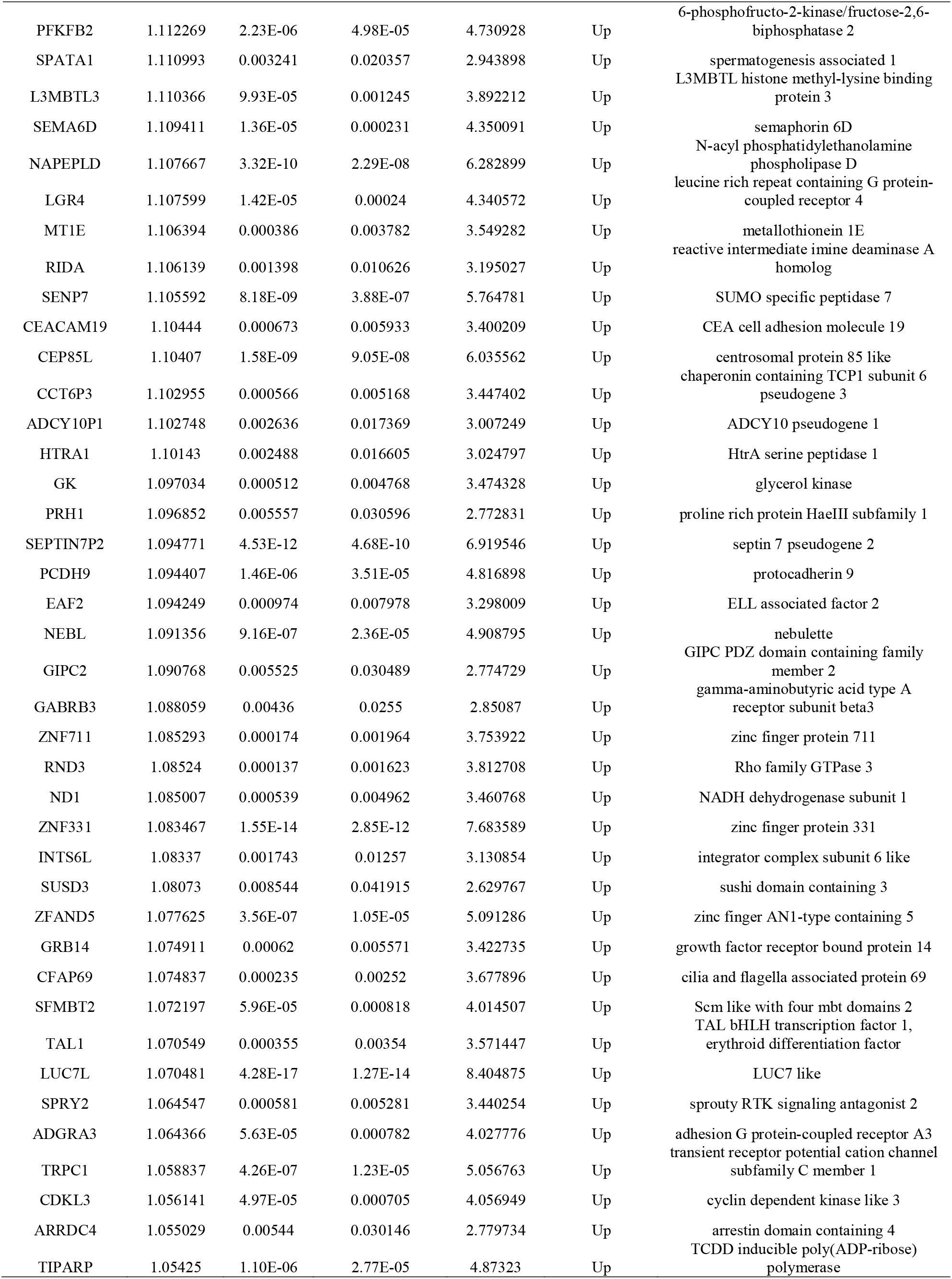
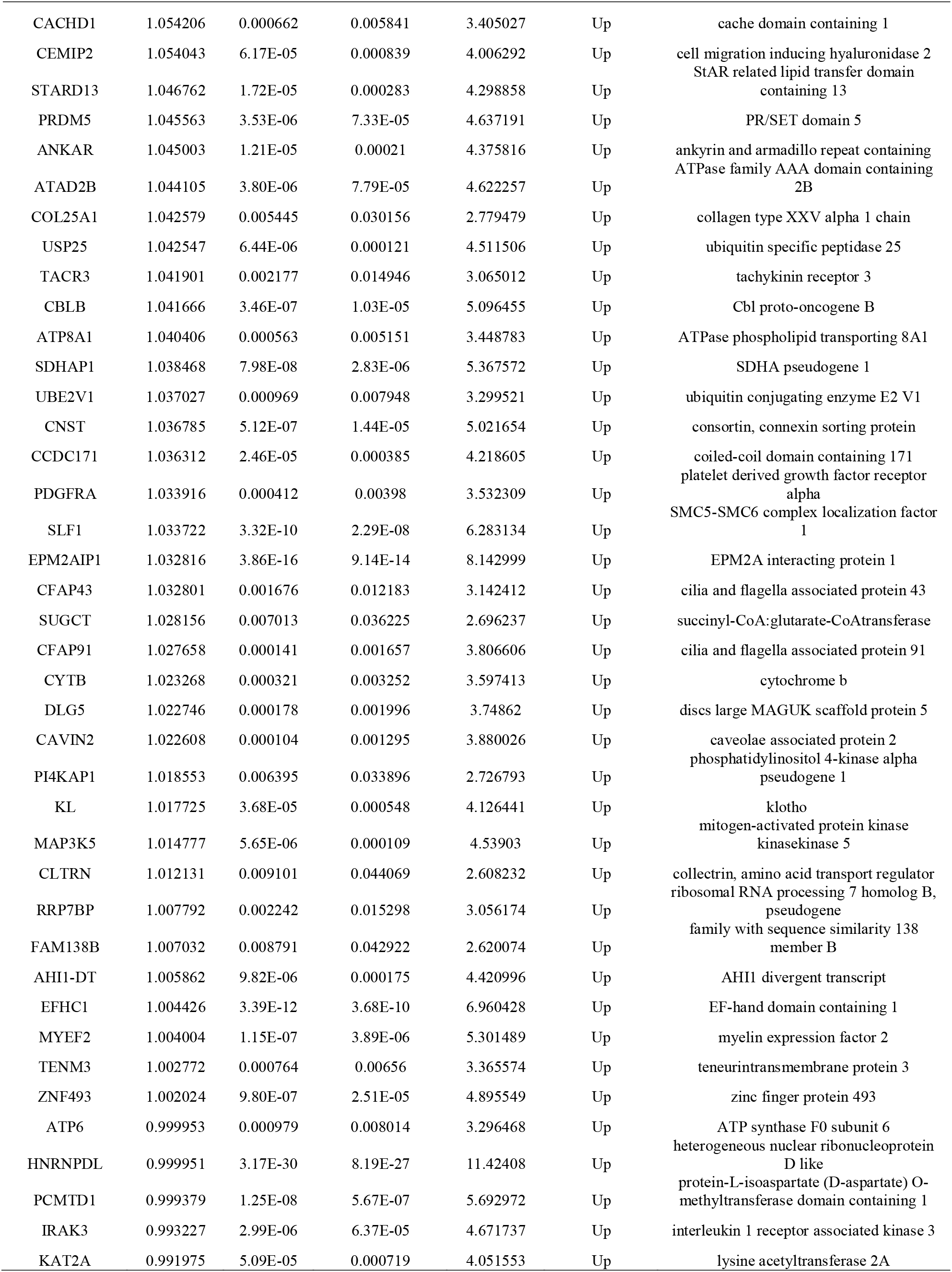
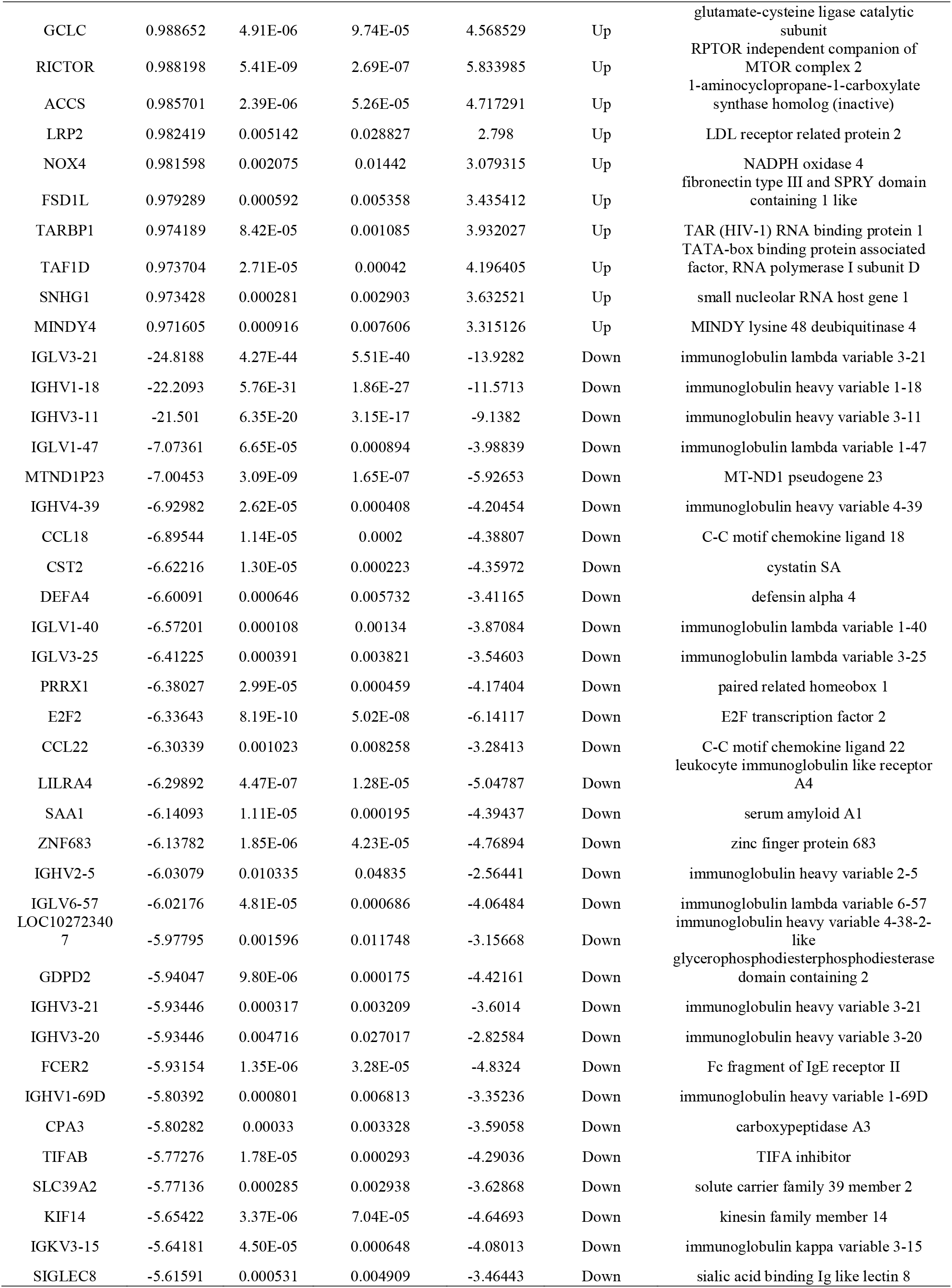
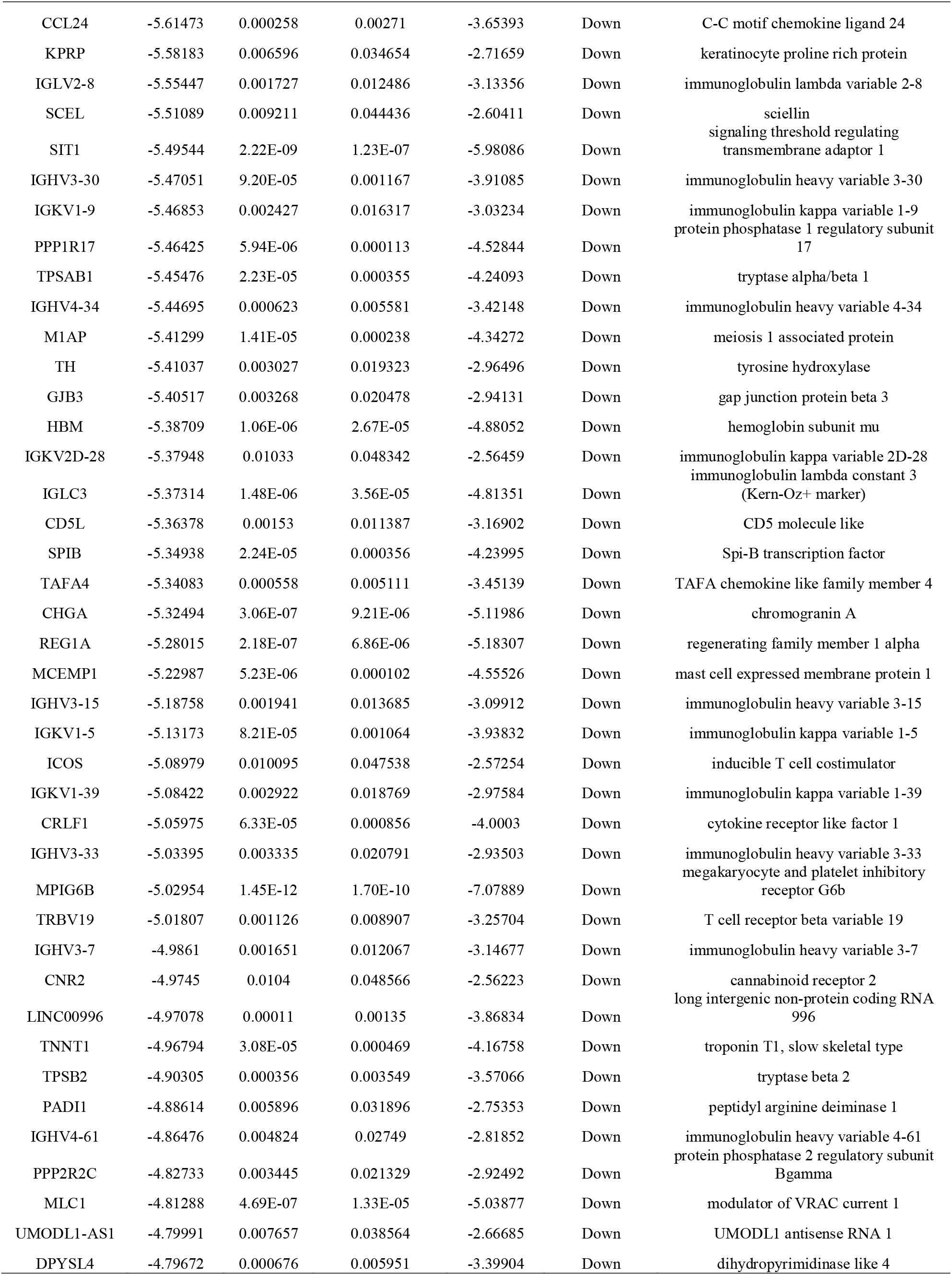
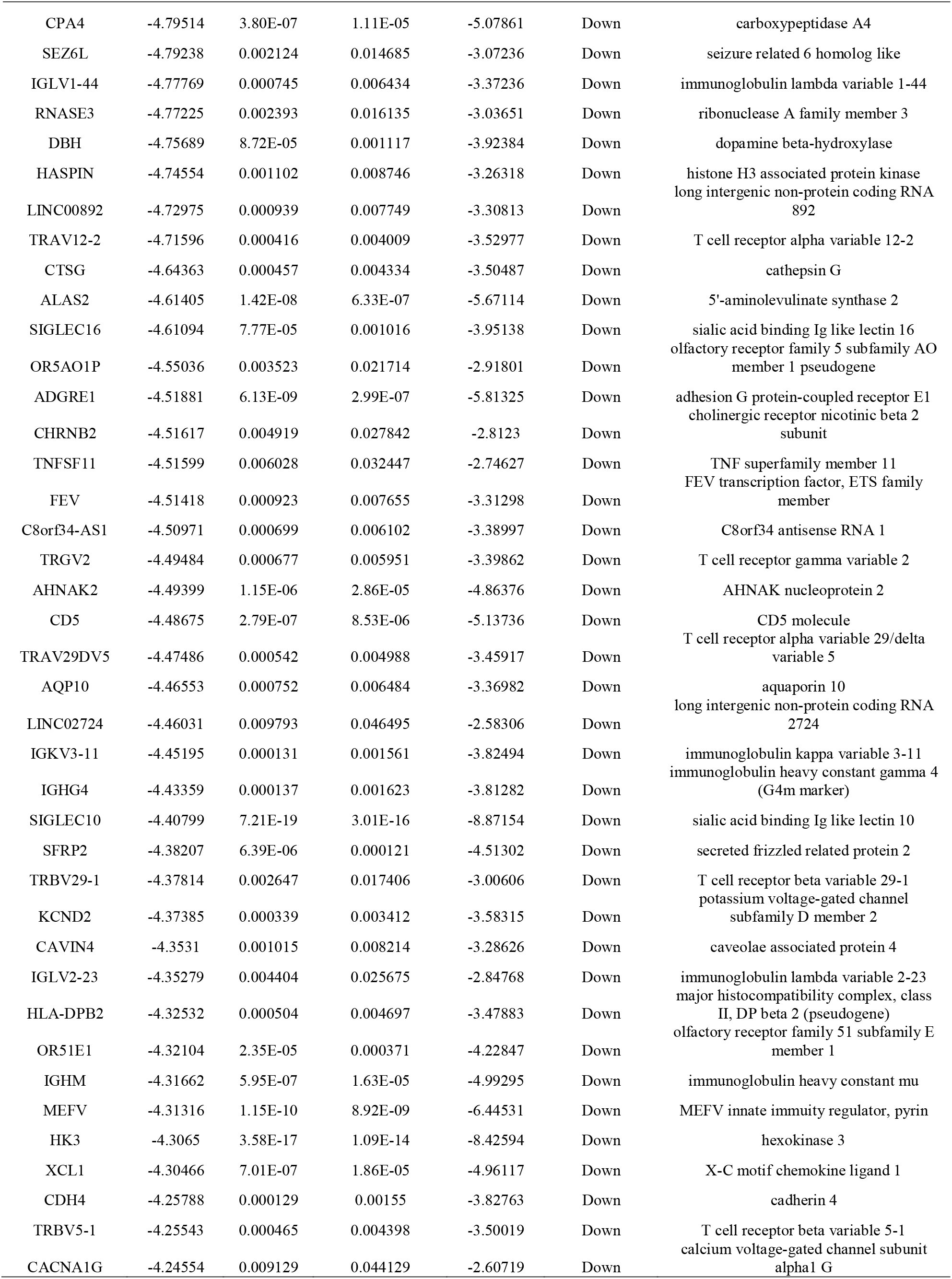
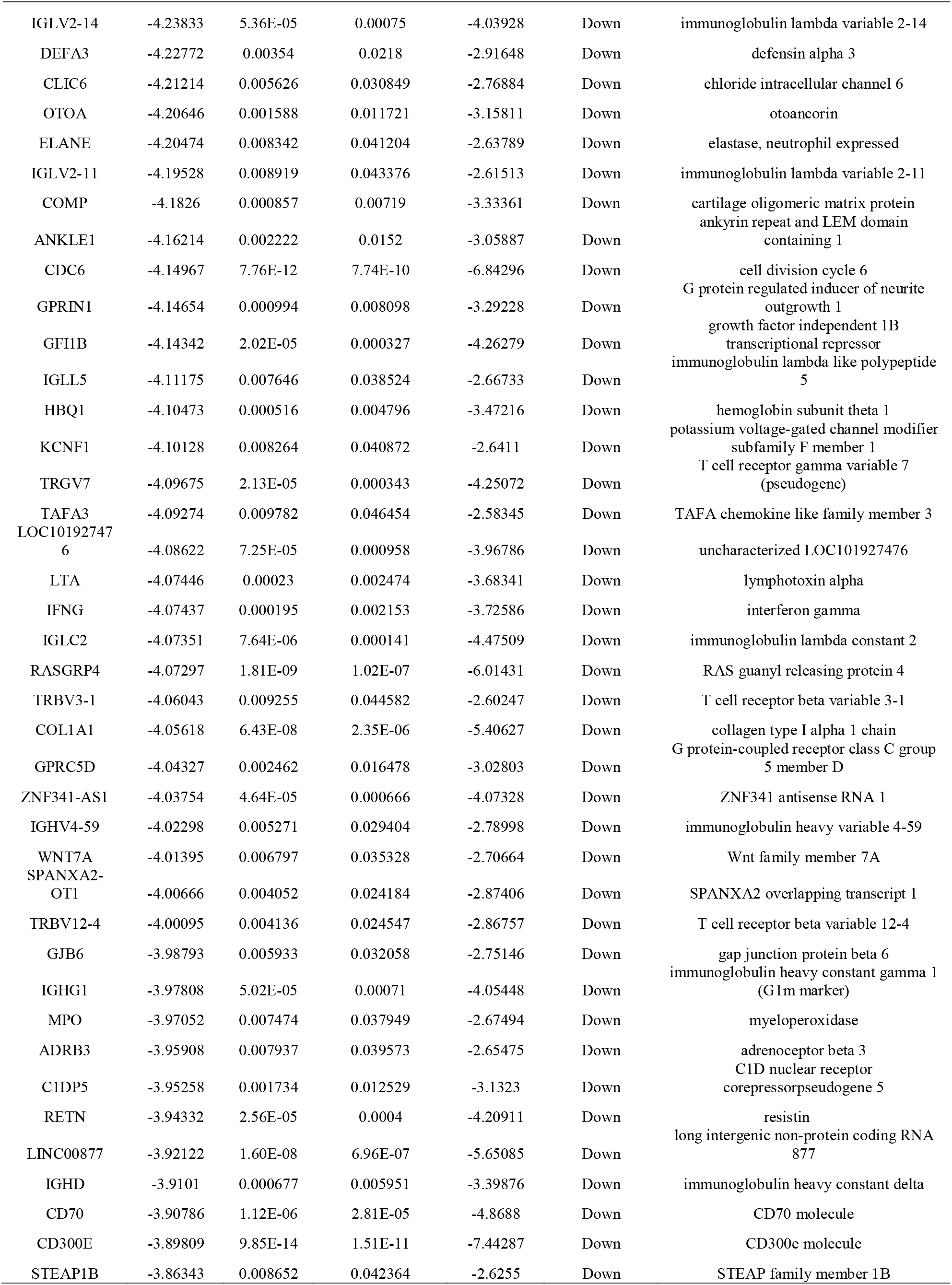
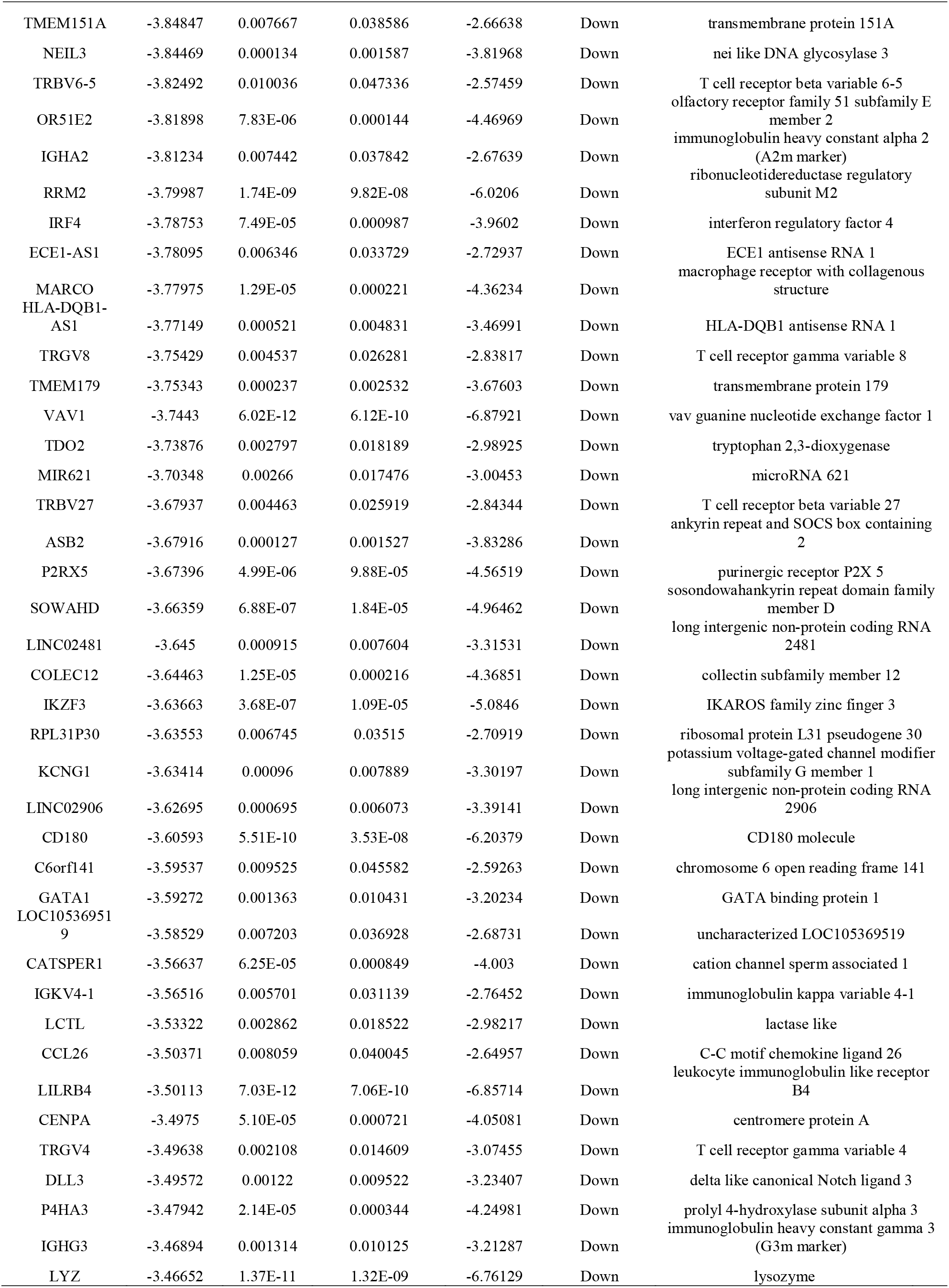
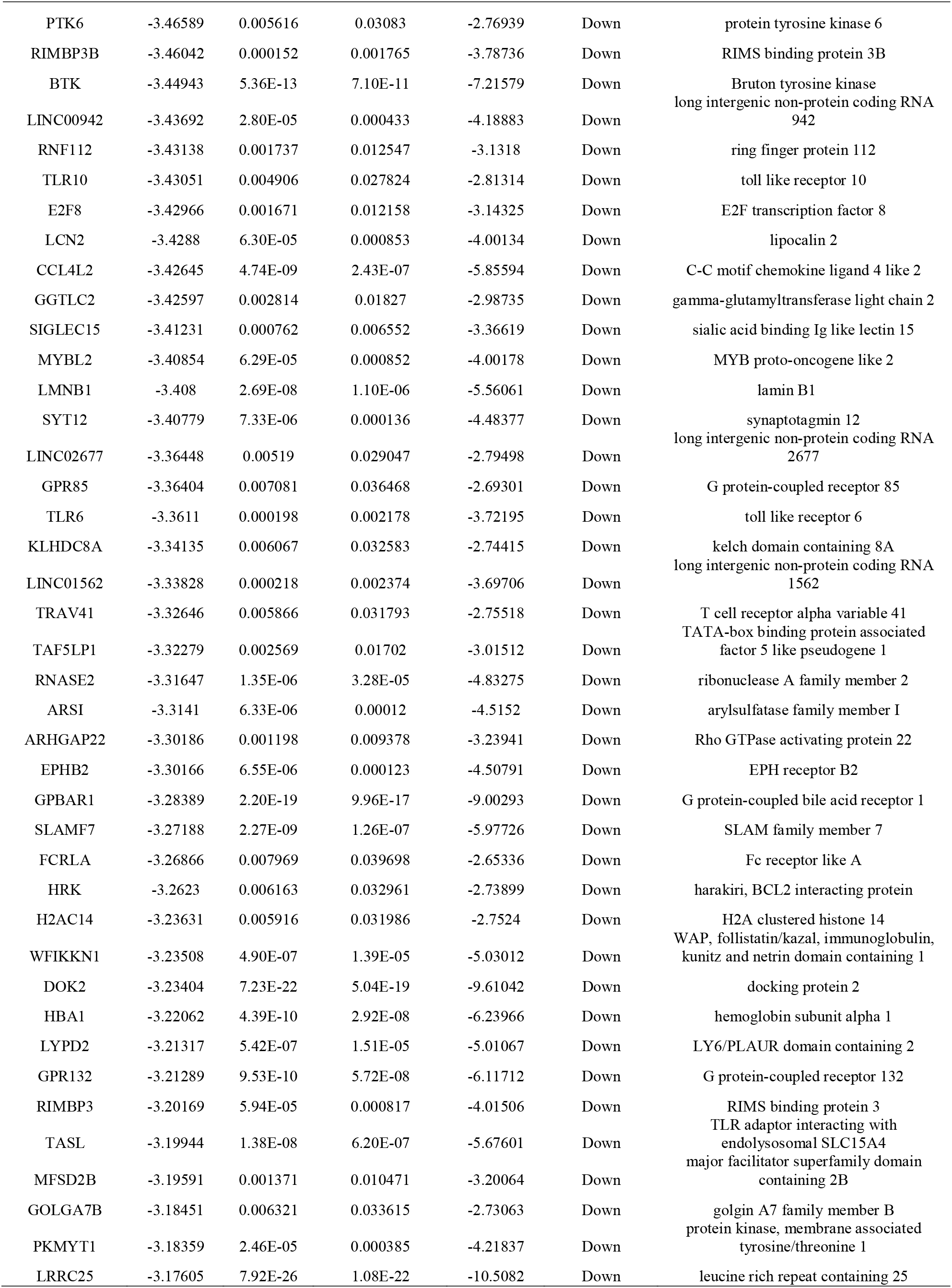
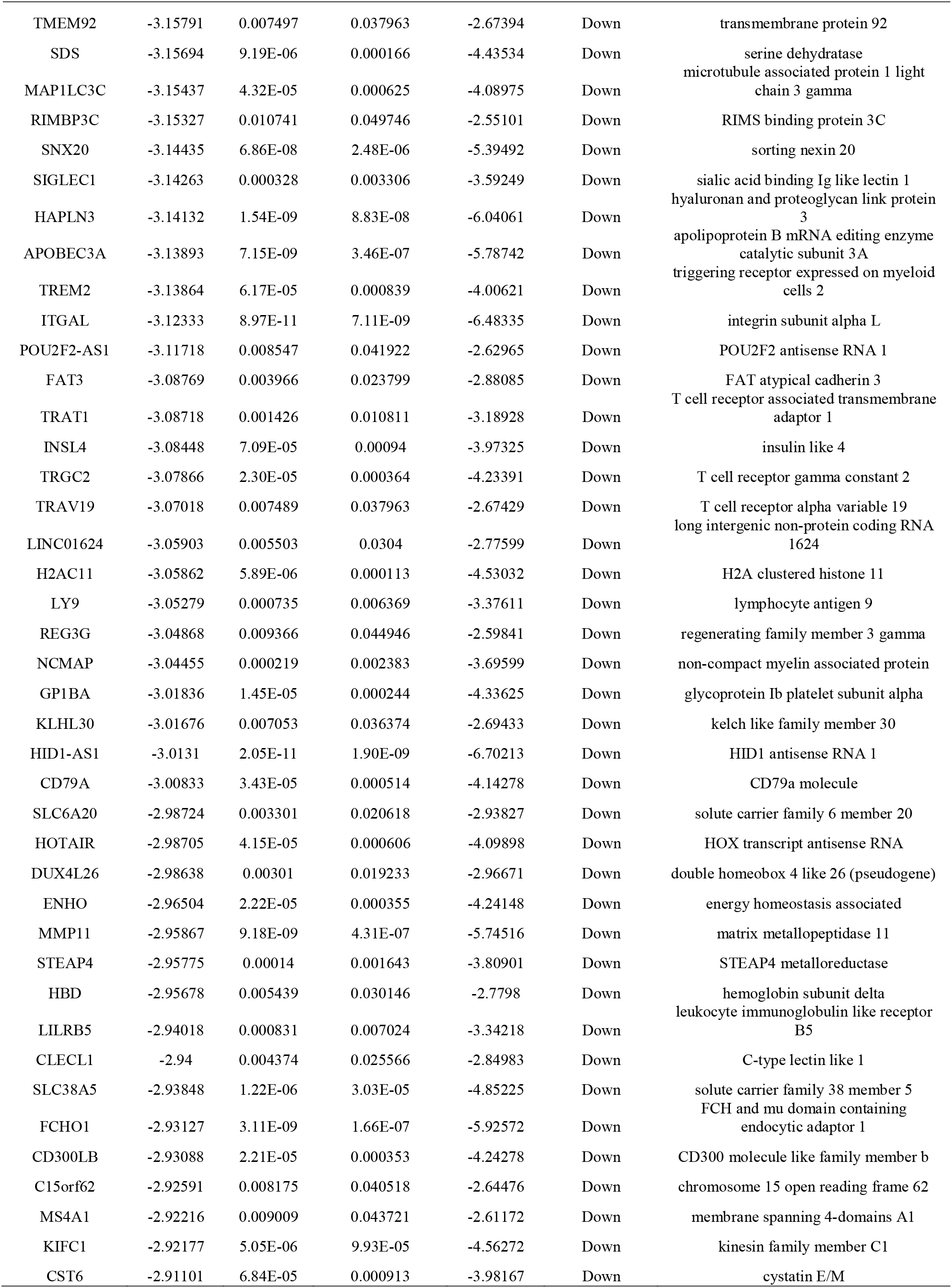
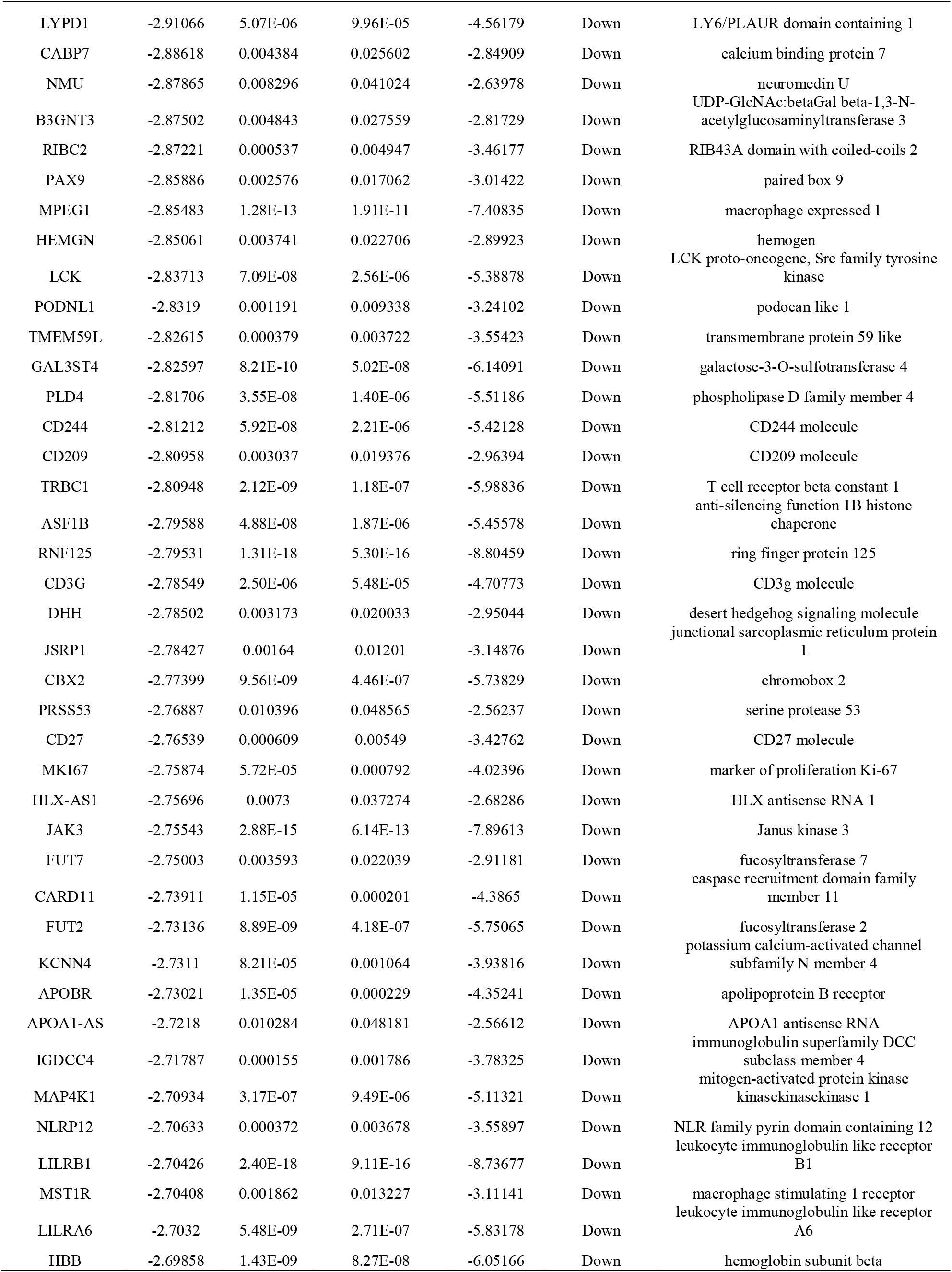
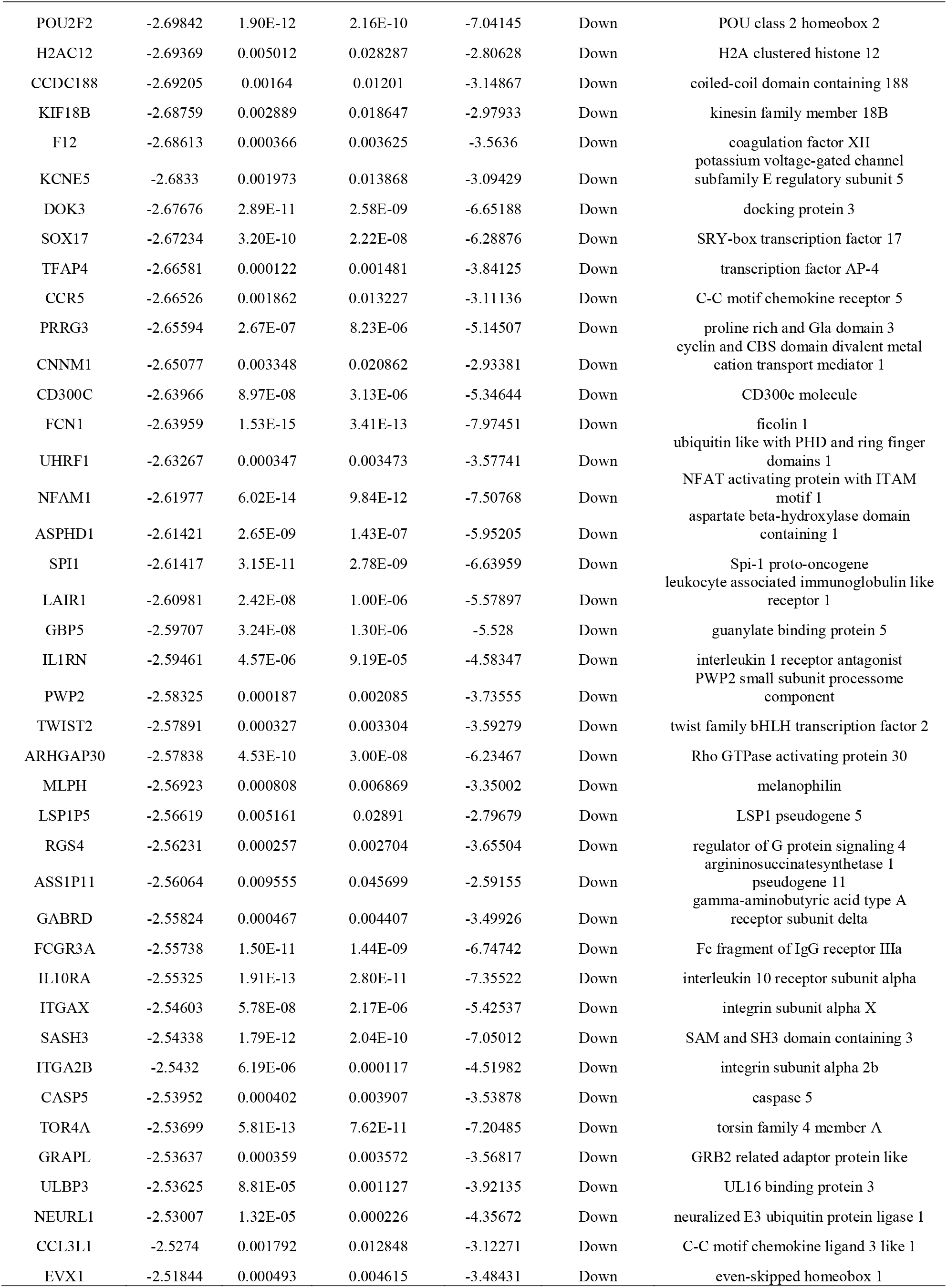
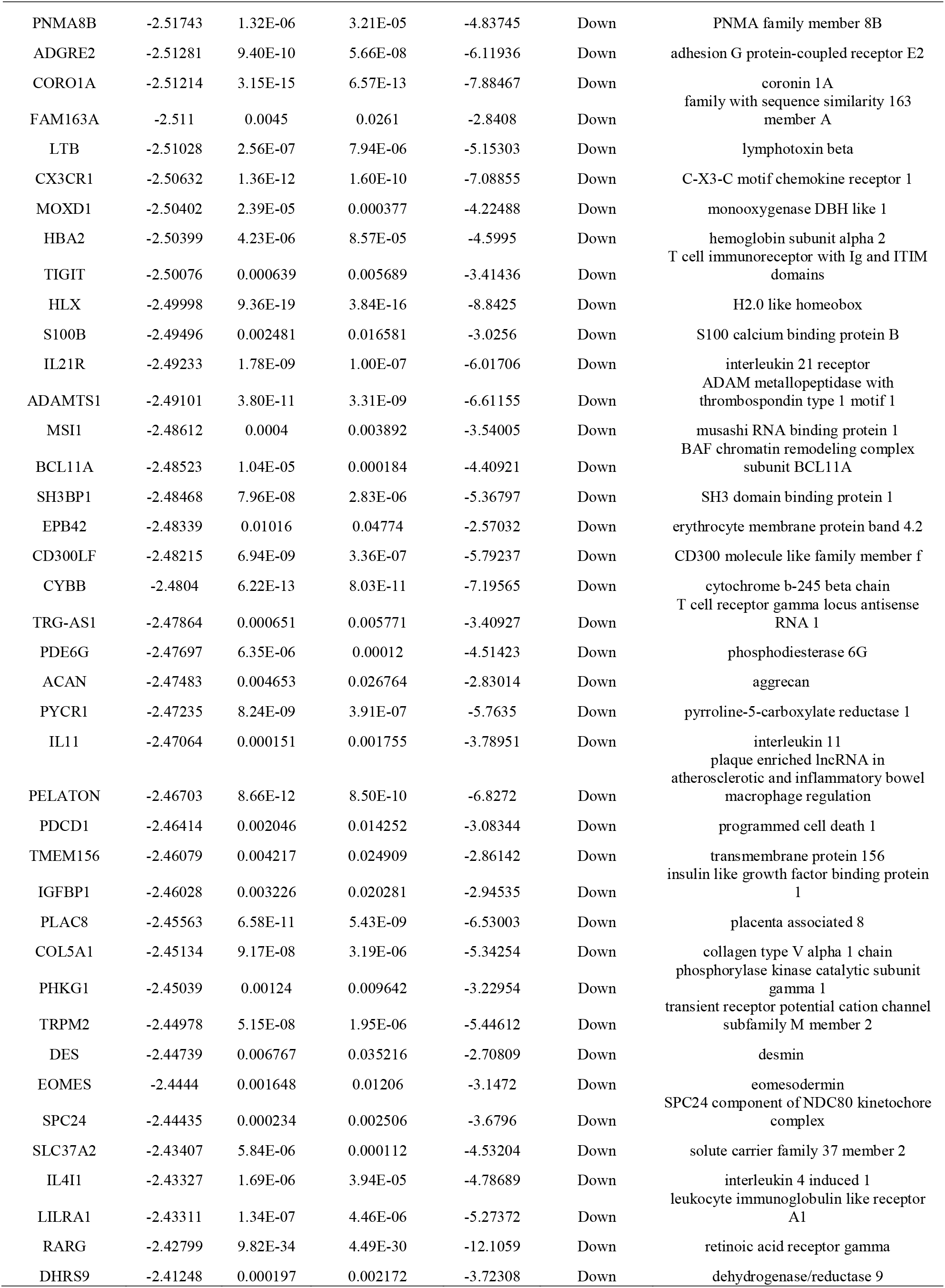
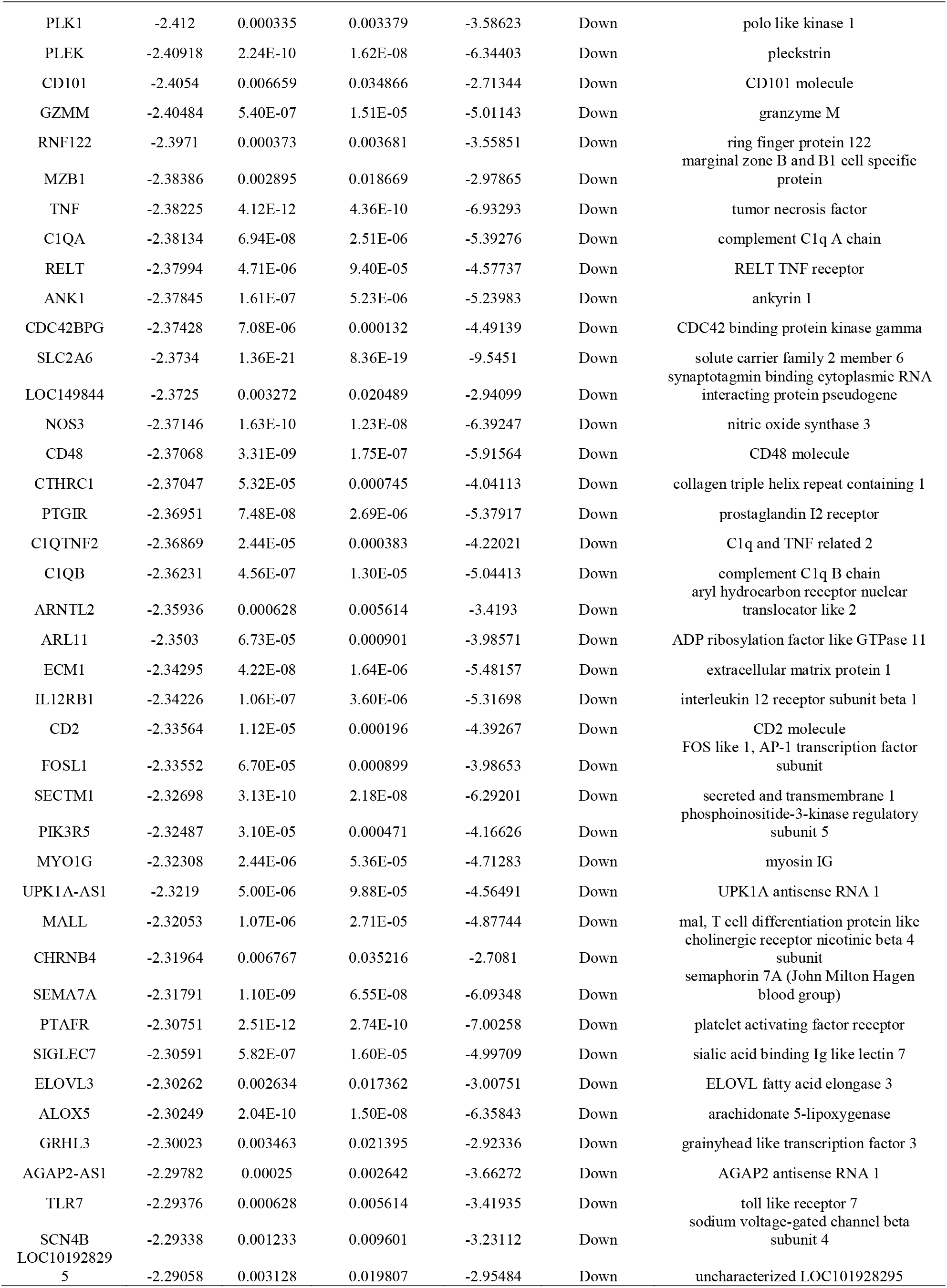
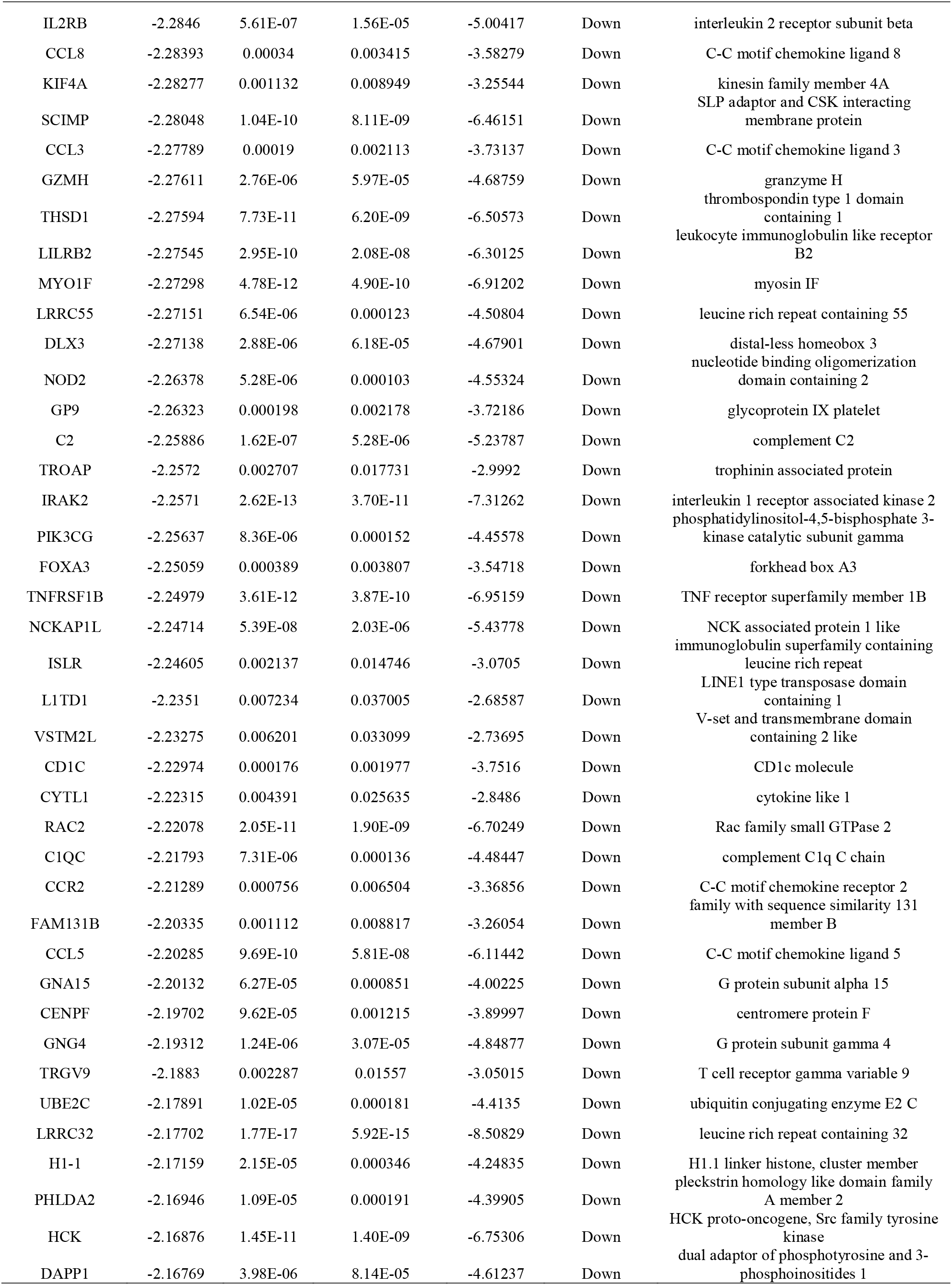
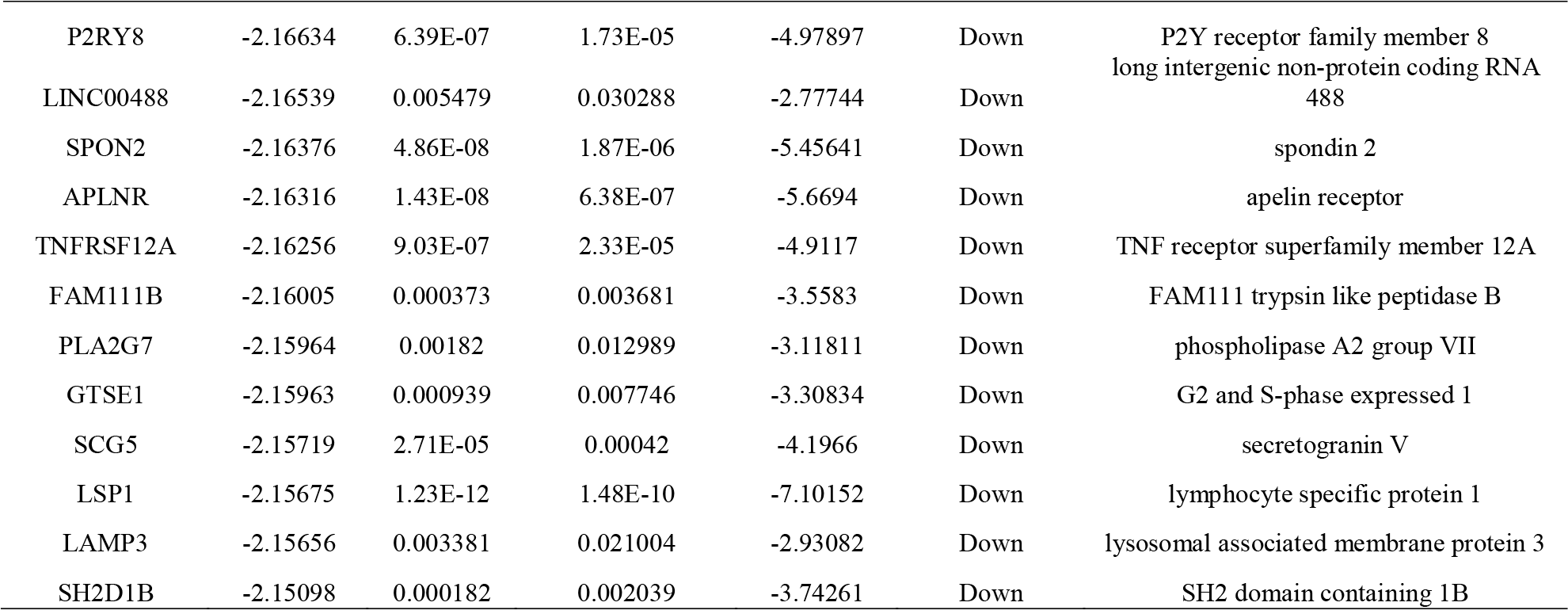
The statistical metrics for key differentially expressed genes (DEGs)

### GO and pathway enrichment analyses of DEGs

GO and REACTOME pathway enrichment analysis in the g:Profiler online software were applied for a deeper comprehension of the identified DEGs. The significant results are presented in Table 2. In the BP group, the up regulated genes were mainly clustered in multicellular organismal process and anatomical structure development, and the down regulated genes were mainly clustered inimmune system process and regulation of cellular process. For the CC group, the up regulated genes were primarily clustered in cell periphery and cell projection. The down regulated genes were primarily clustered in cell periphery and membrane. The up regulated genes in the MF group were mostly clustered in protein binding and ion binding, and the down regulated genes were mostly clustered in signaling receptor binding and molecular transducer activity. The top significantly enriched REACTOME pathways for the DEGs were also displayed by the g:Profiler online software and are presented in Table 3. The up regulated genes were associated with metabolism and transport of small molecules, while the down regulated genes were involved in immune system and signaling by interleukins.

**Table 2.**
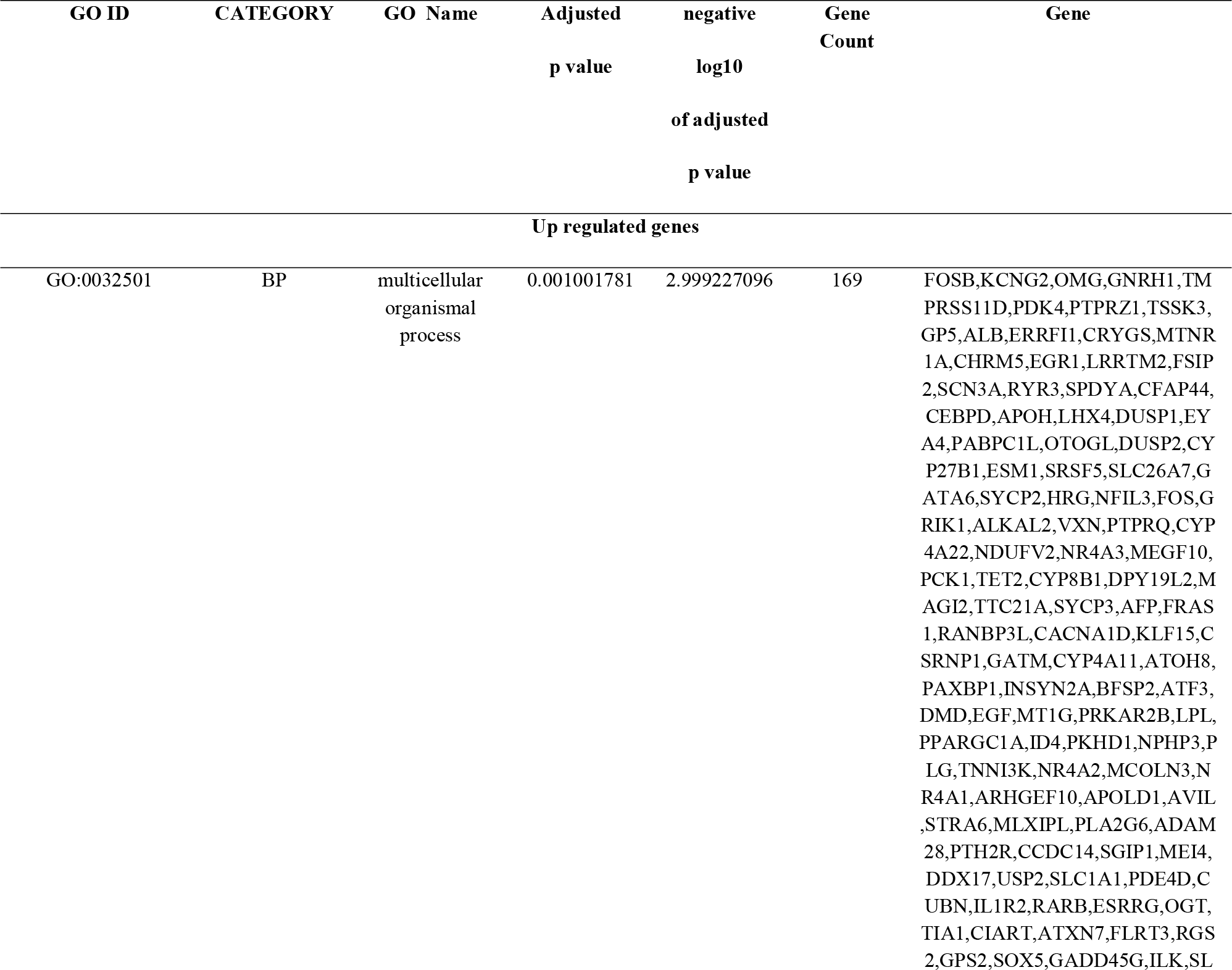
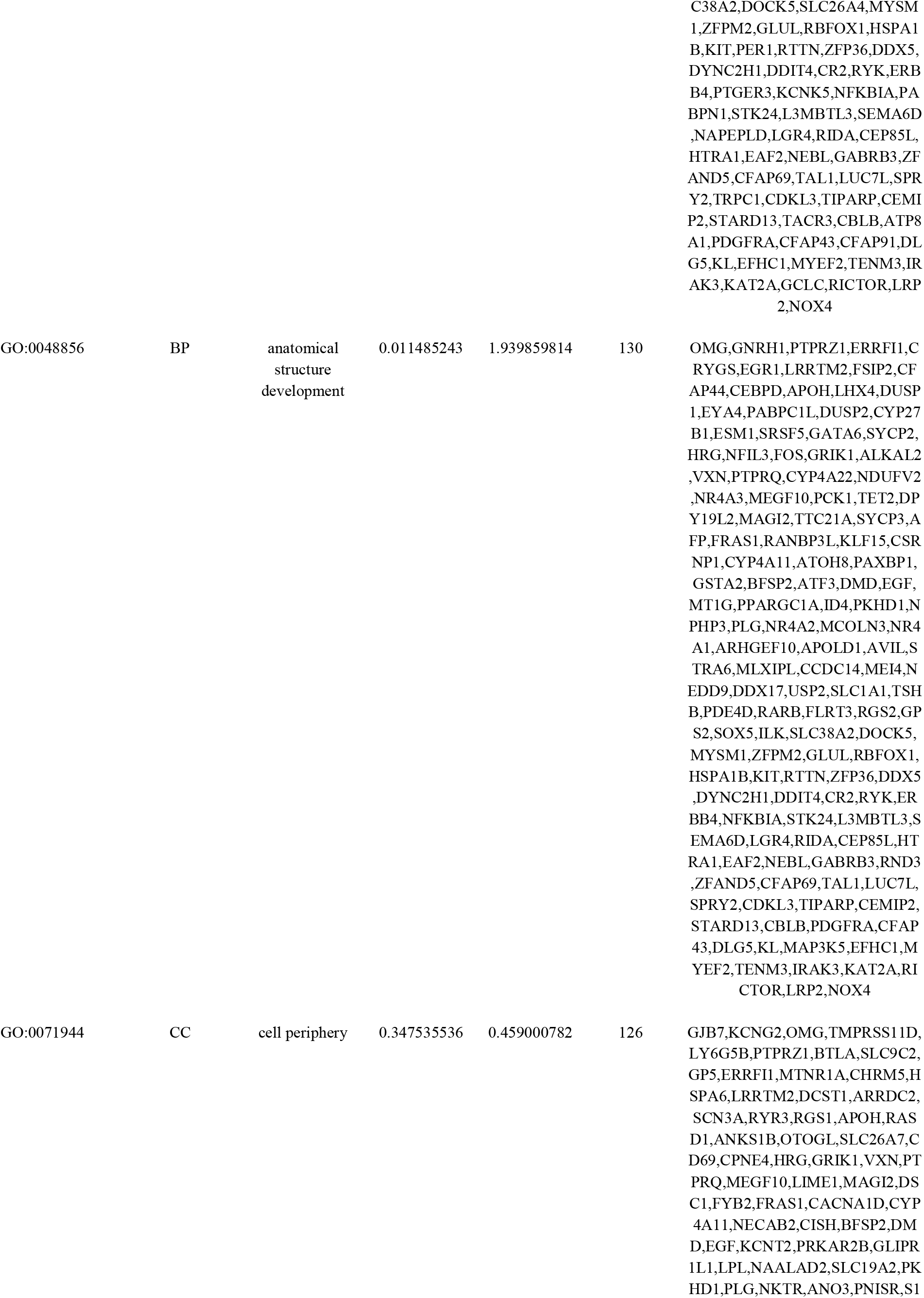
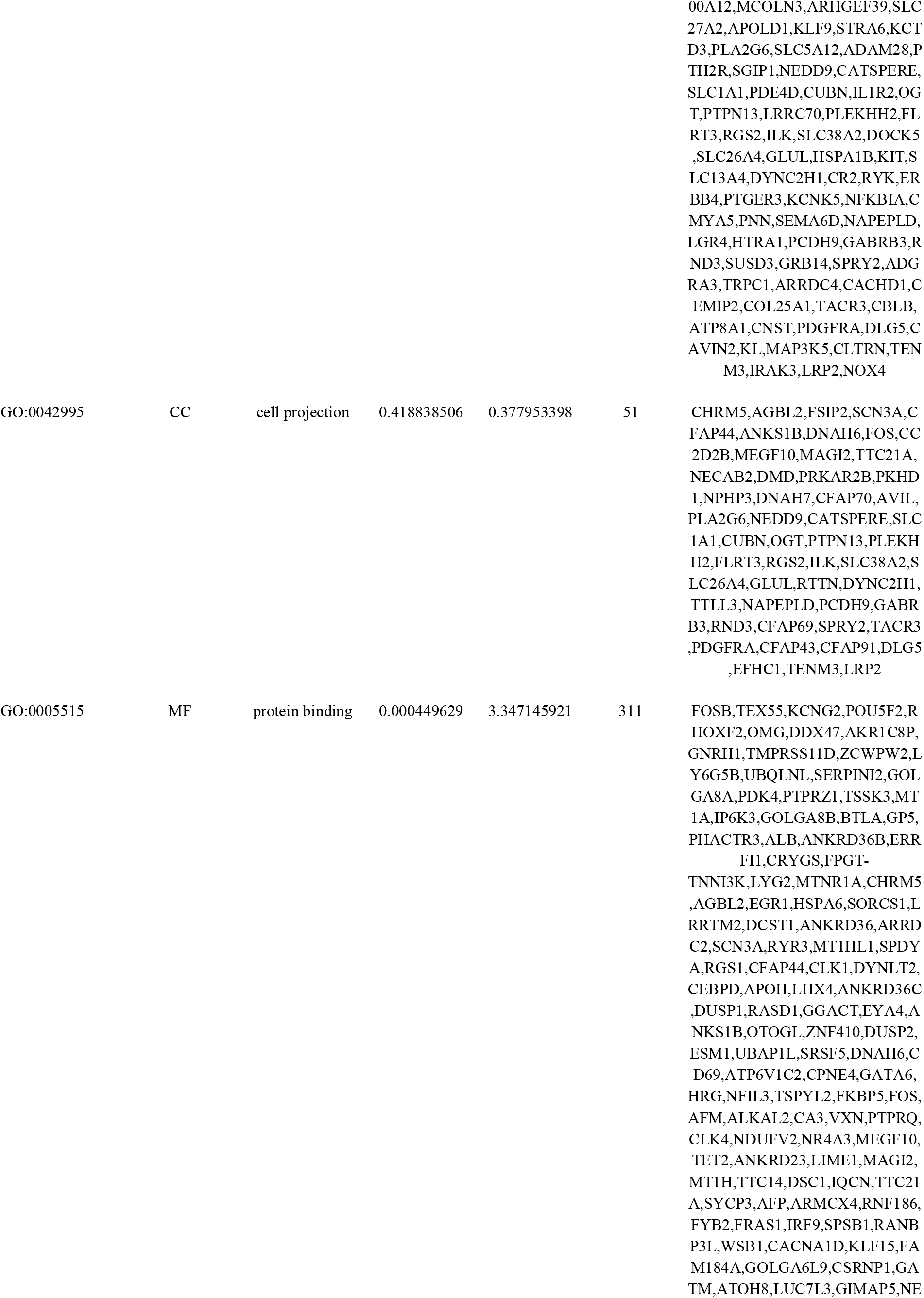
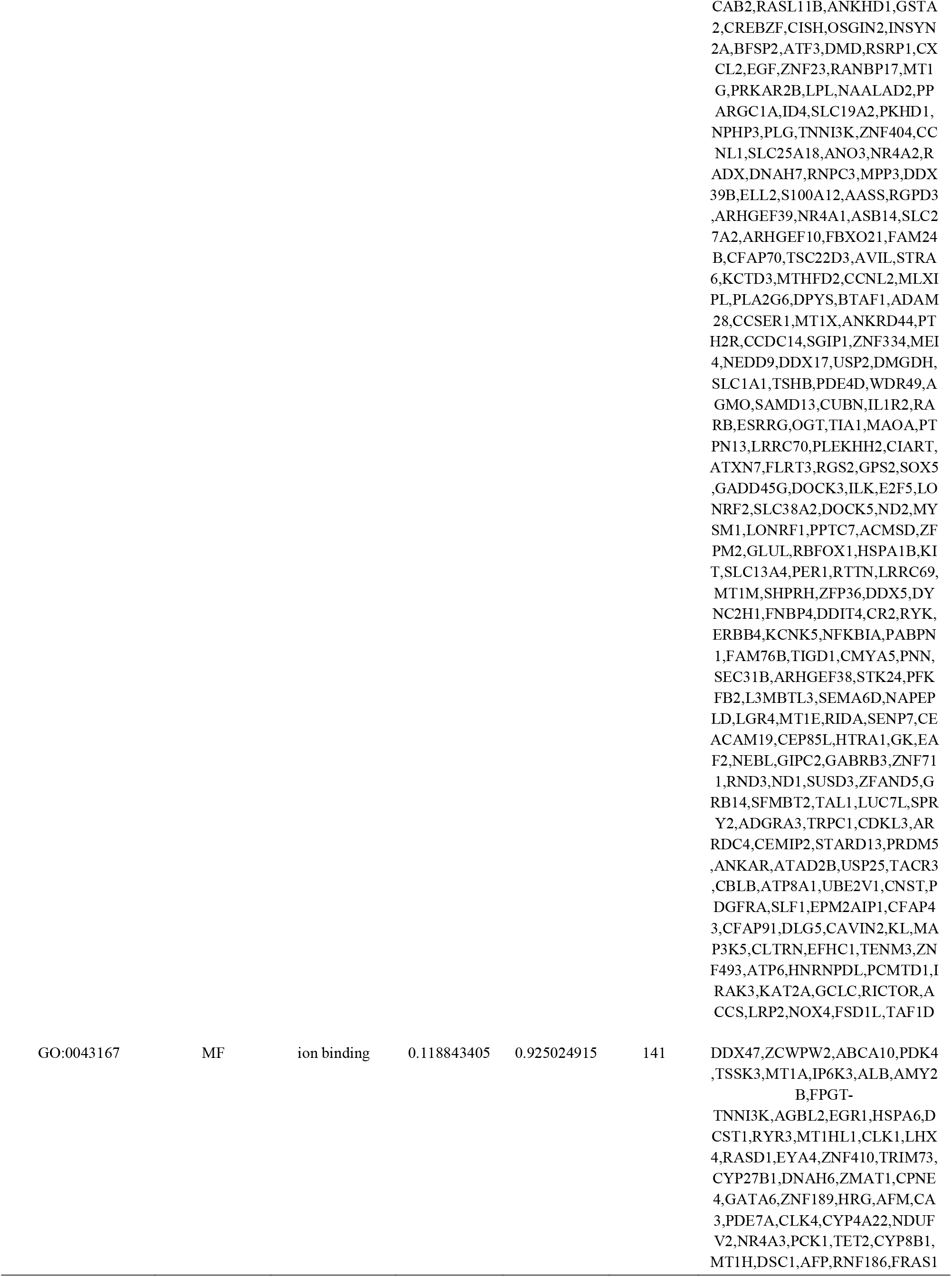
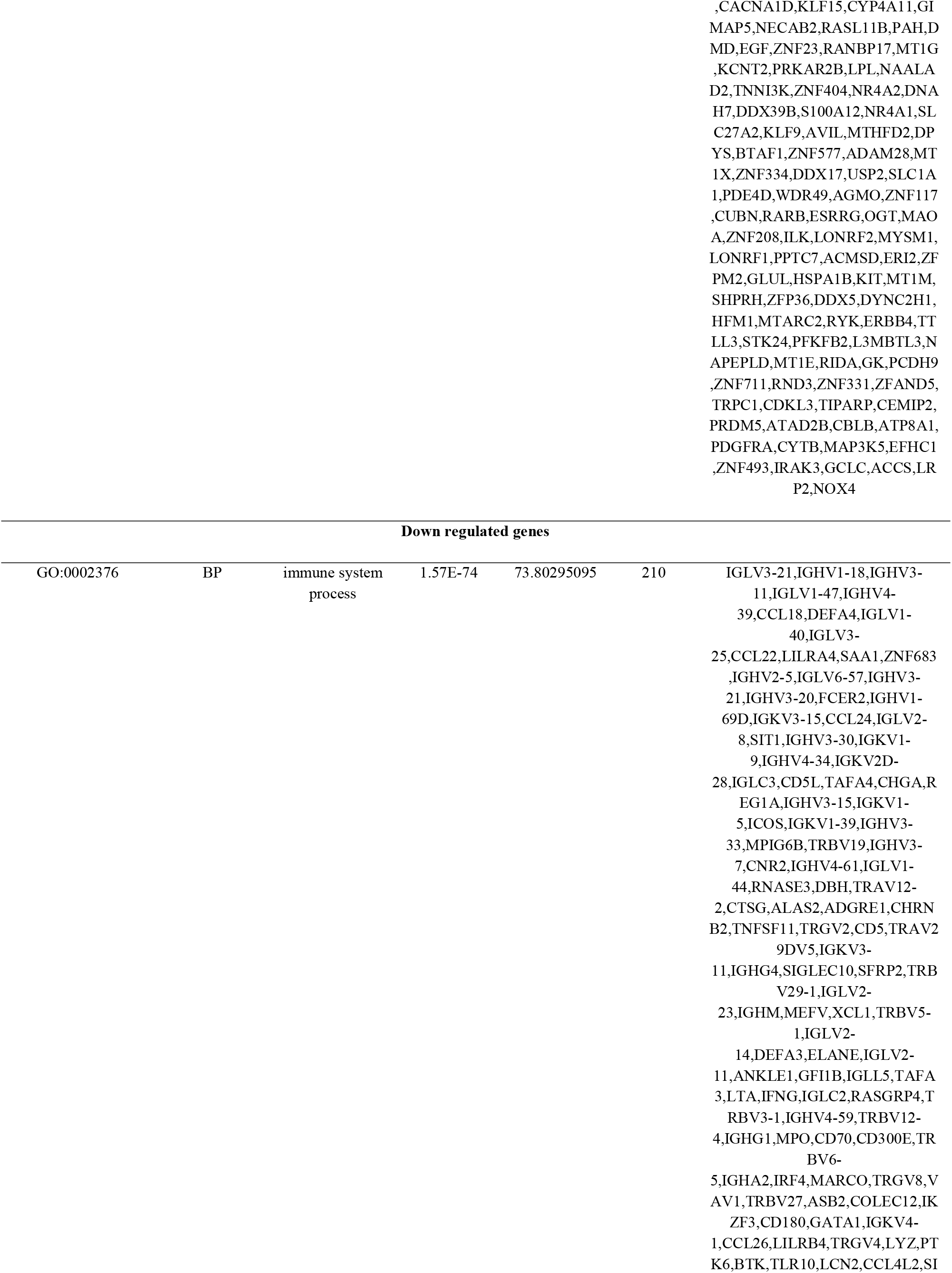
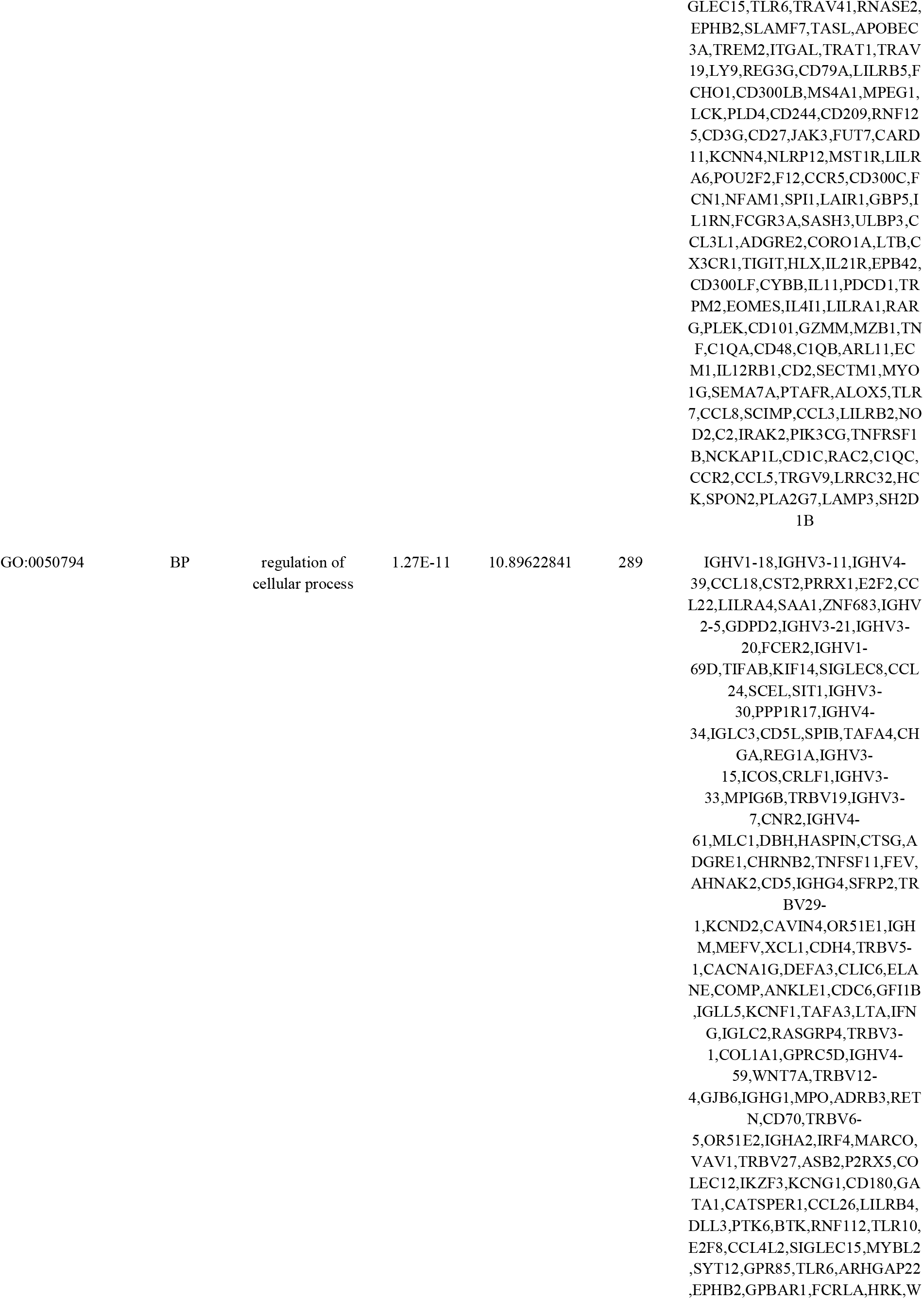
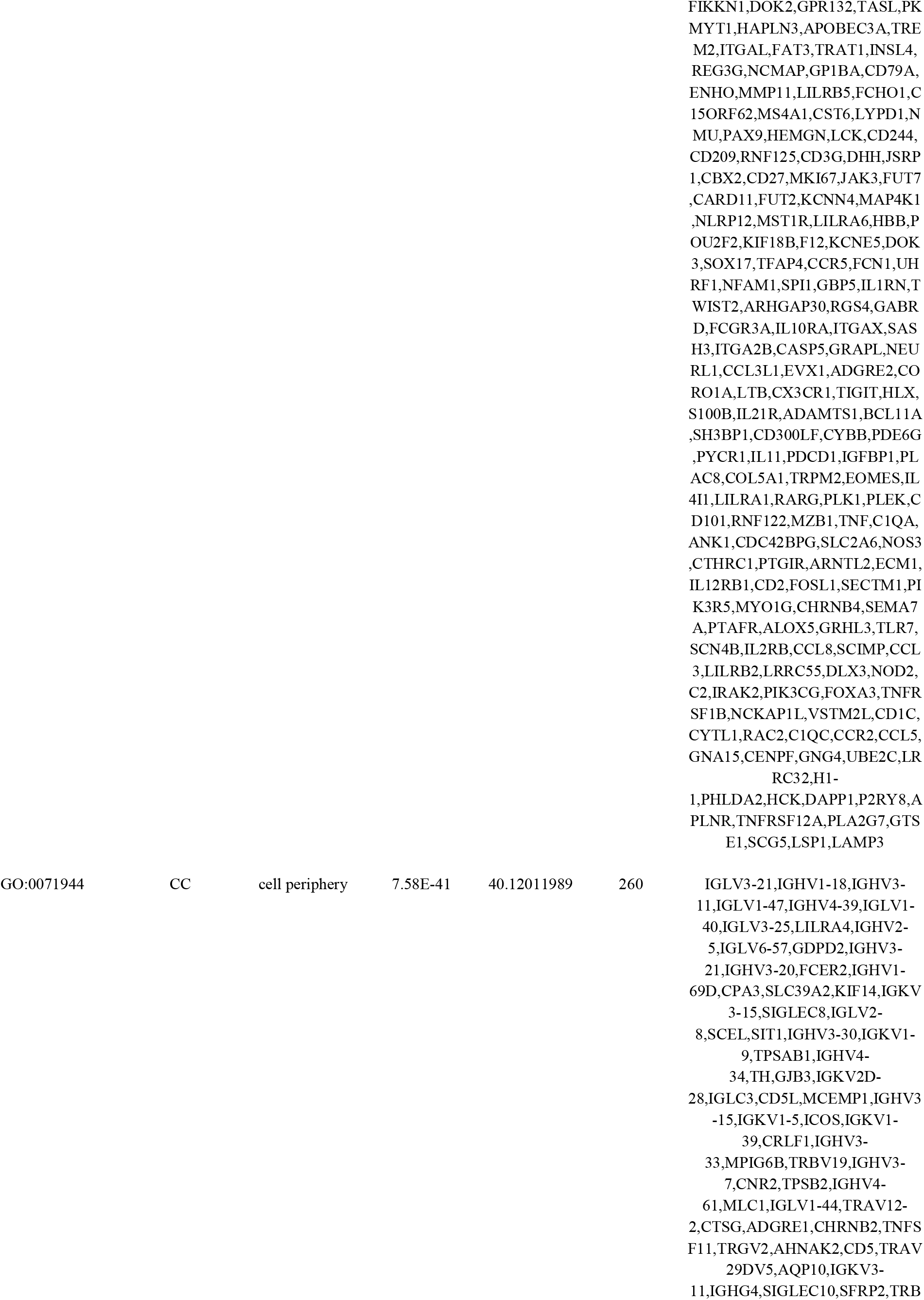
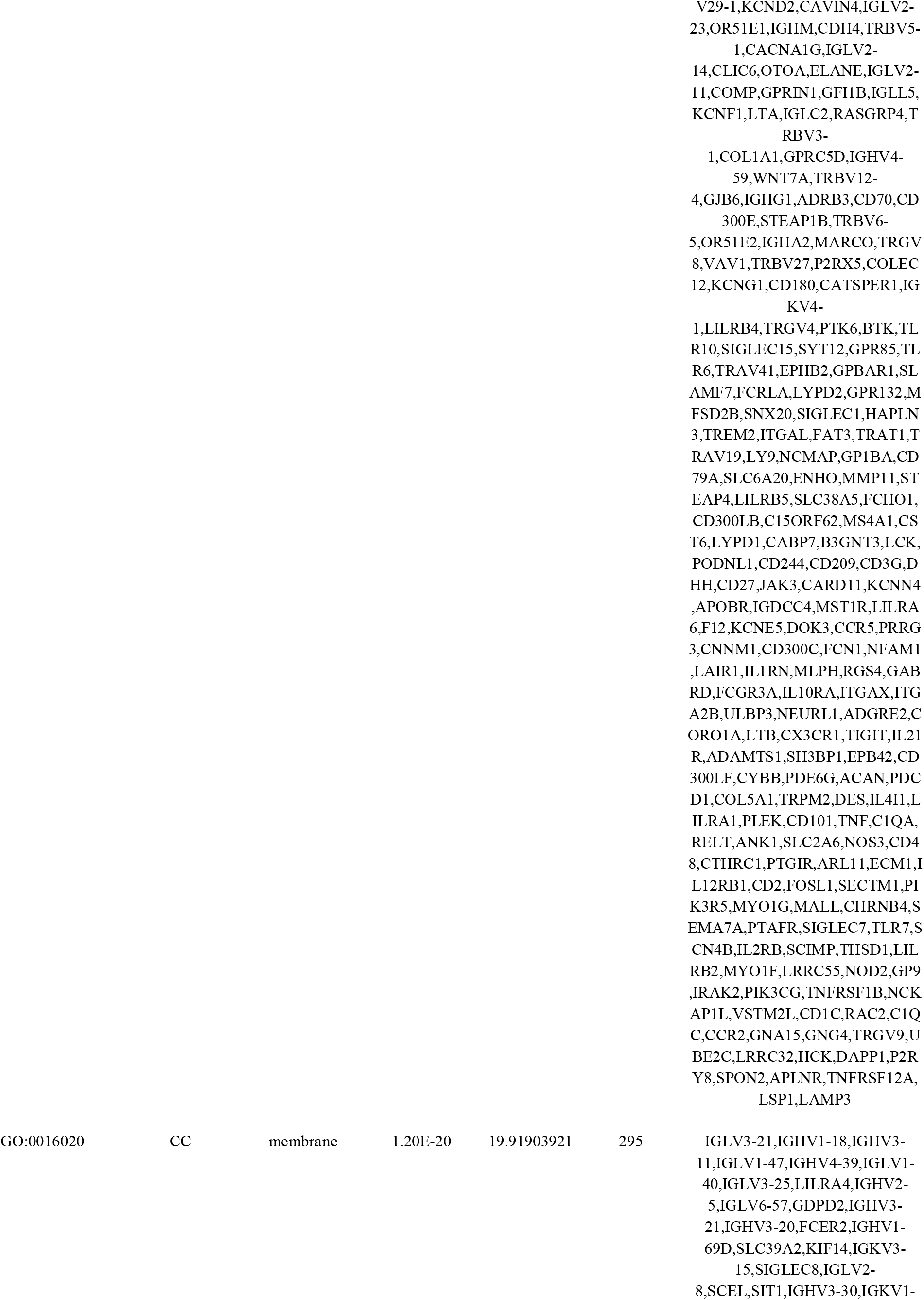
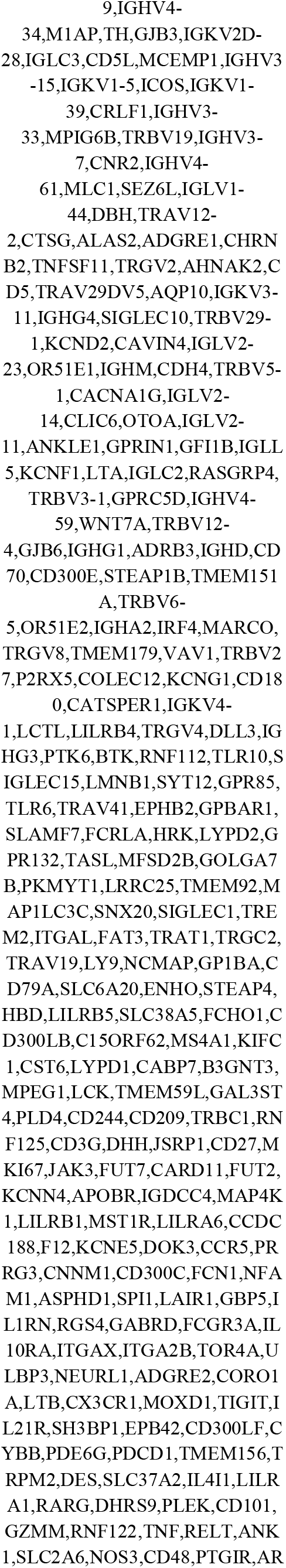
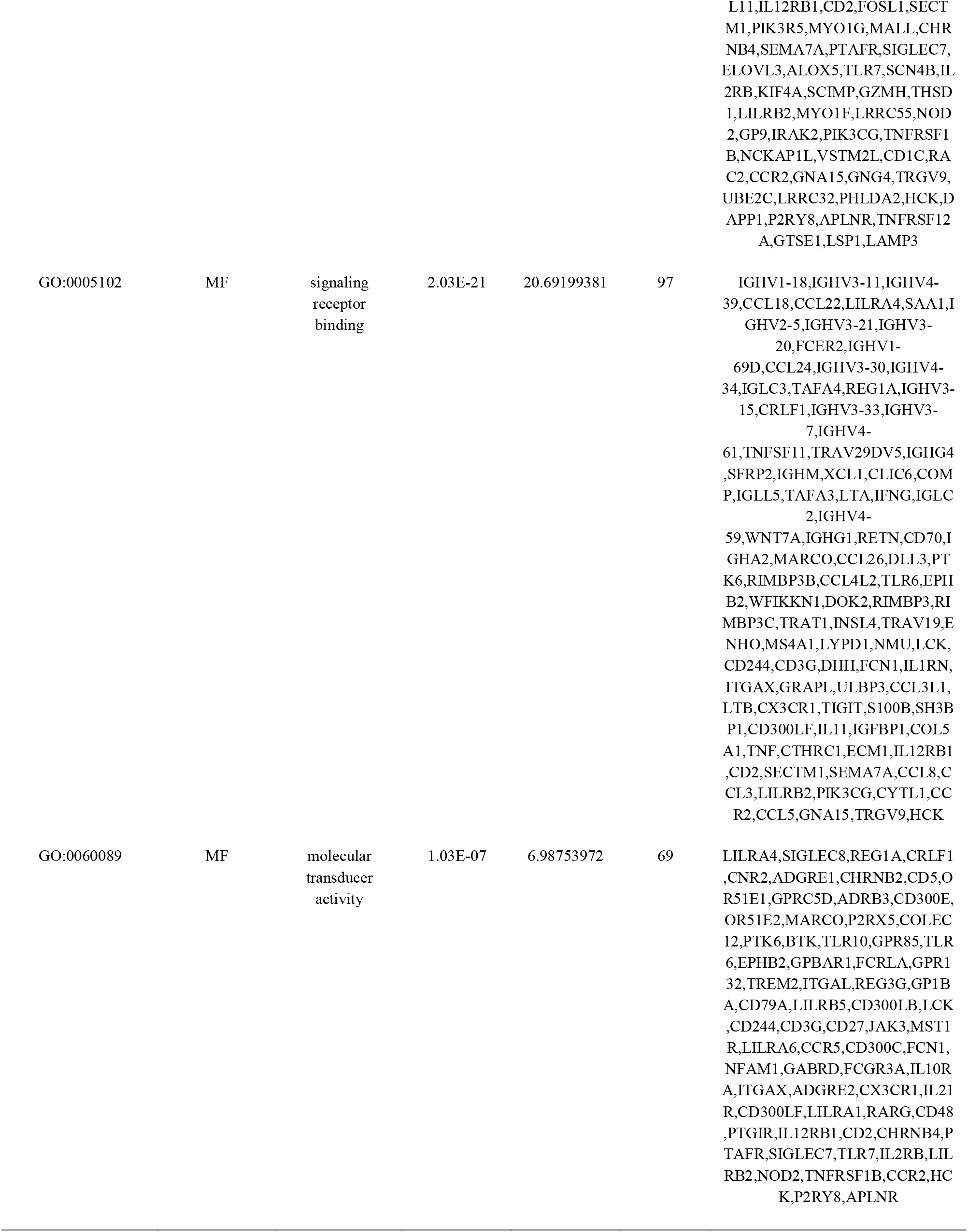
The enriched GO terms of the up and down regulated differentially expressed genes

**Table 3.**
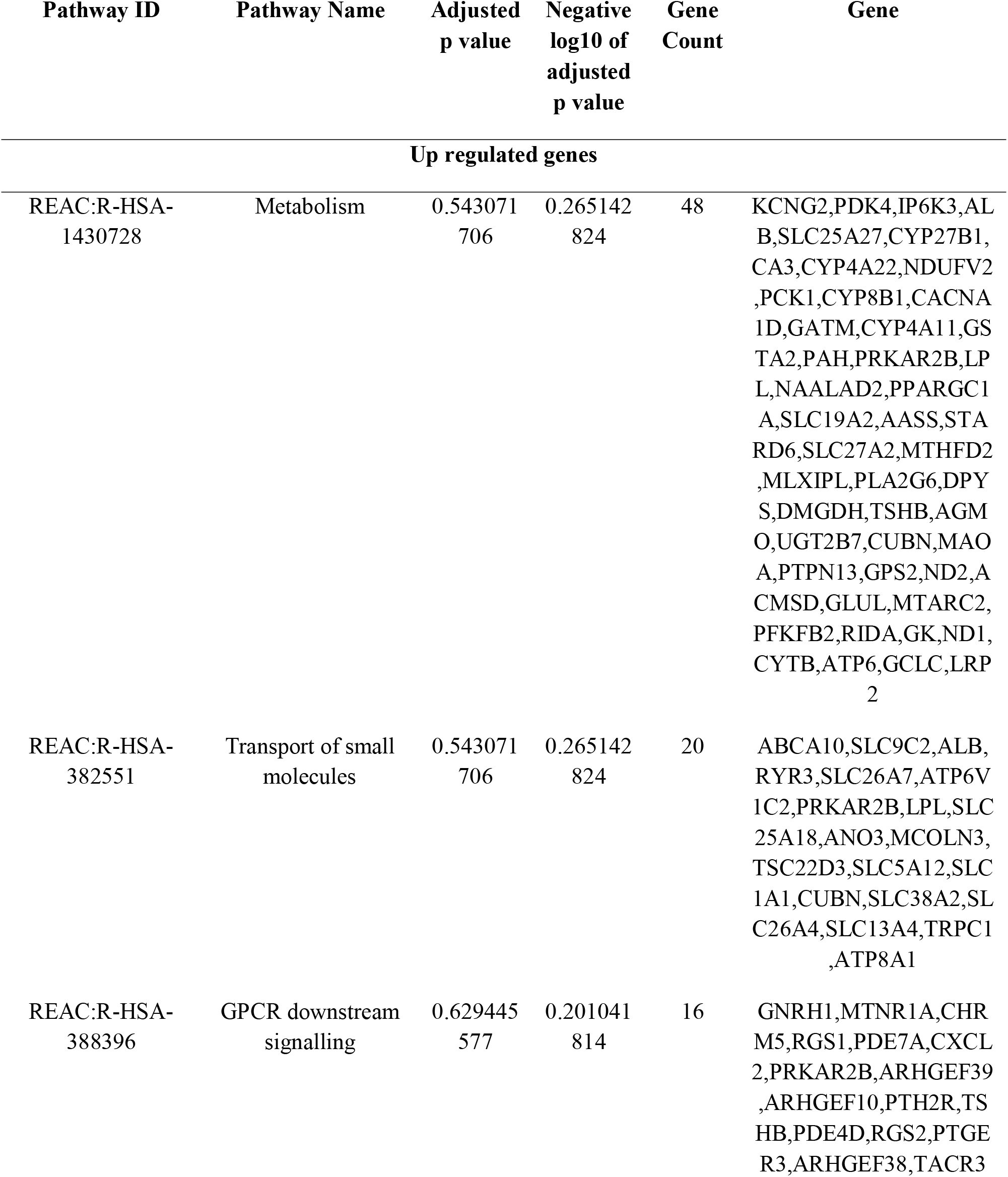
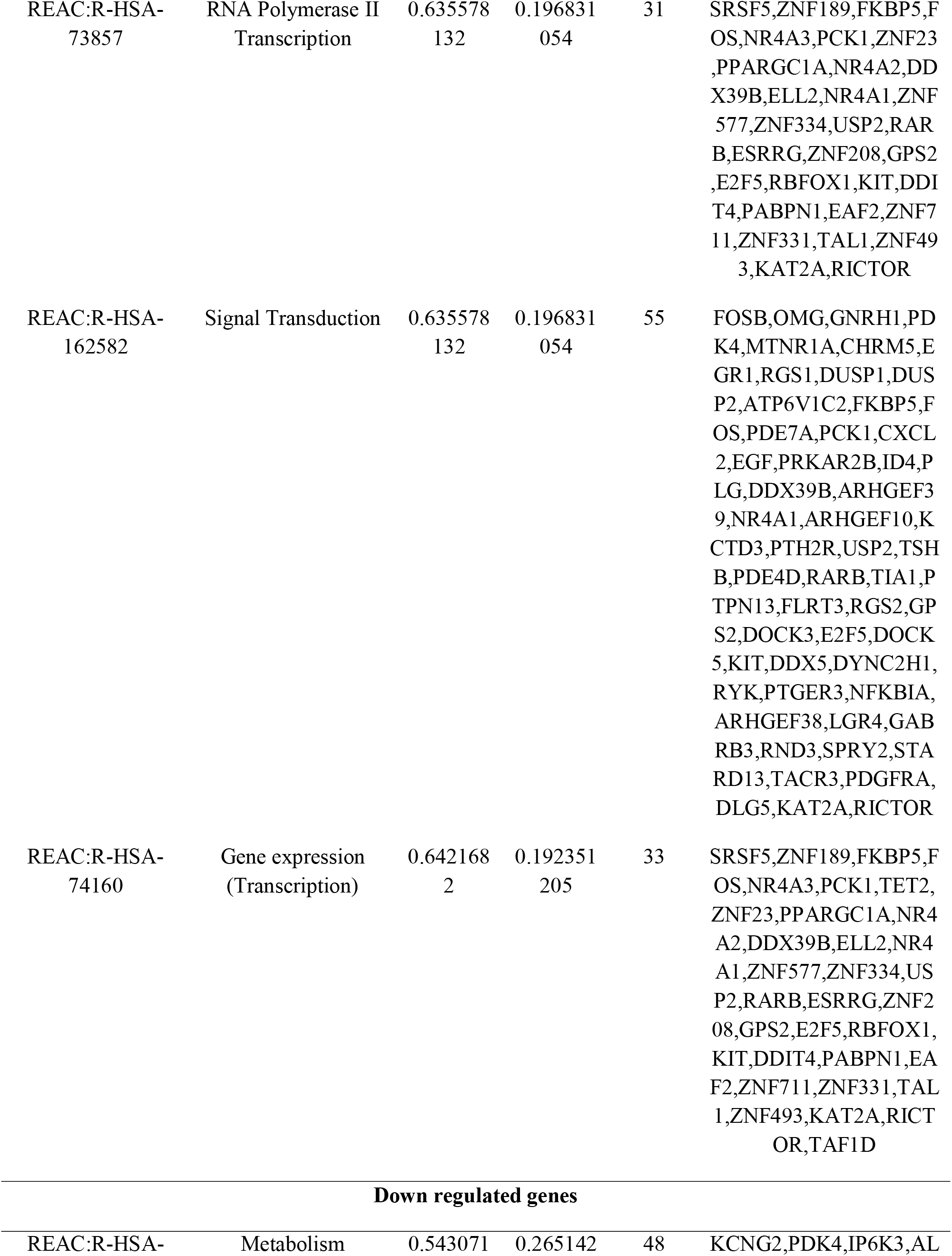
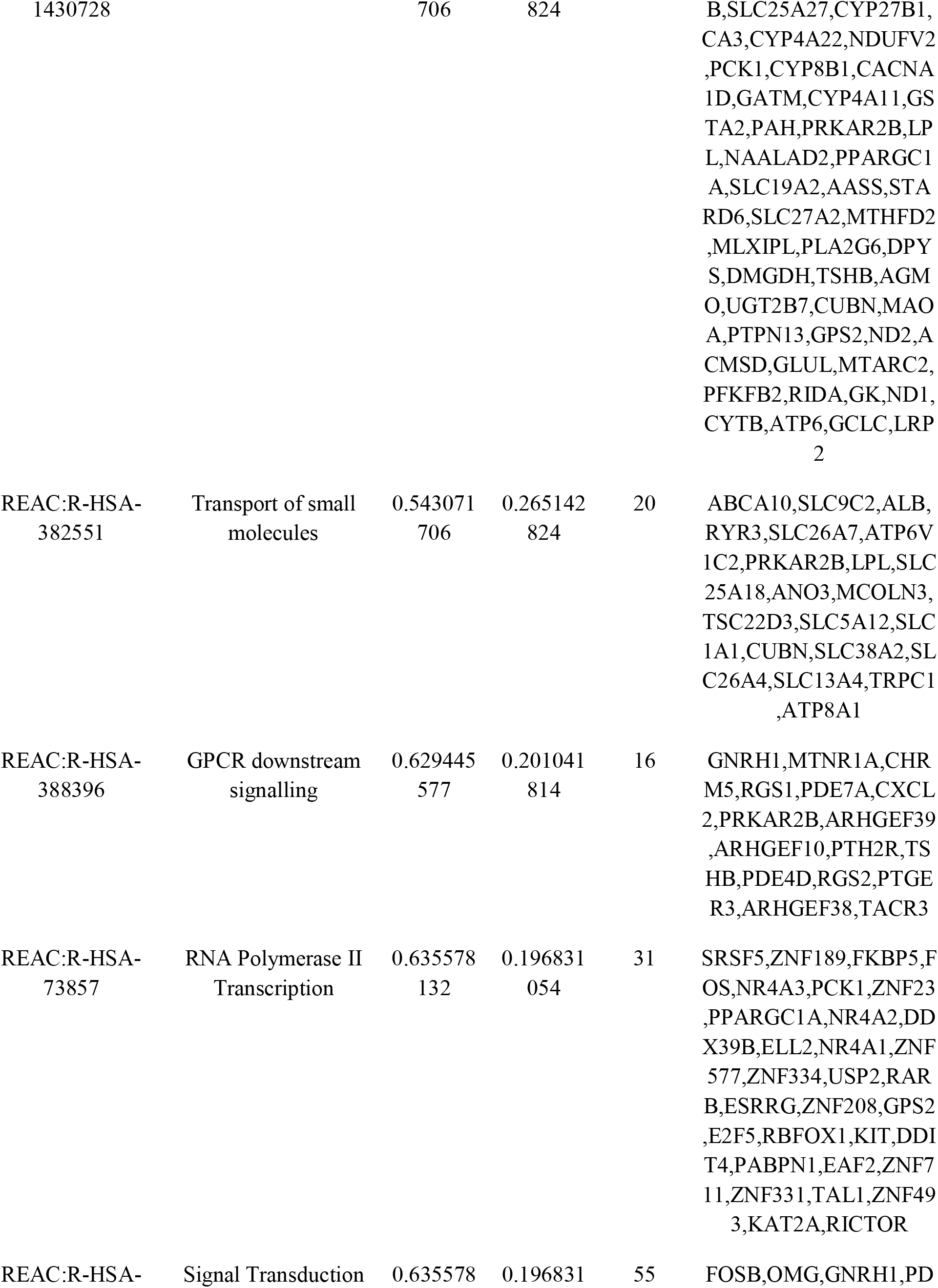
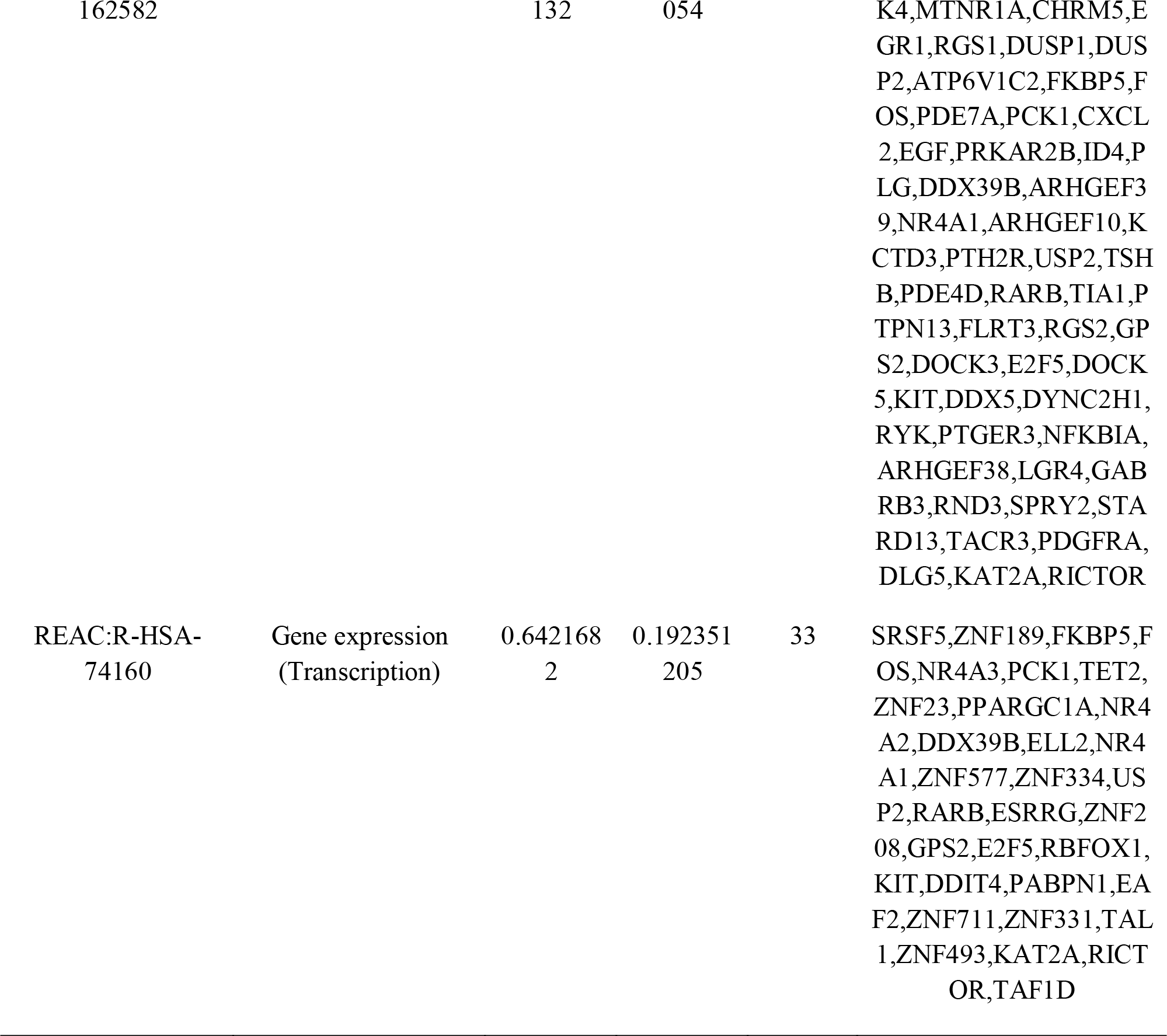
The enriched pathway terms of the up and down regulated differentially expressed genes

### Construction of the PPI network and module analysis

The DEG expression products in T1DM were constructed into PPI networks using the HIPPIE in Cytoscape software. The PPI network is shown in Fig. 4. There were 7117 nodes and 16762 edges in the network. Cytoscape software was applied to calculate the node degree, betweenness, stress and closeness of each node in the PPI network. The top genes were selected as hub genes, including ILK, DDX5, MATR3, ALB, FOS, MKI67, PLK1, TNF, LCK and GTSE1. The node degree, betweenness, stress and closeness of all nodes are shown in Table 4. PEWCC1 was applied to screen modules of the PPI network. The significant module 1 contained 26 nodes and 54 edges (Fig. 4A) and significant module 2 contained 14 nodes and 31 edges (Fig. 4B). Results showed that genes in these modules were mainly enriched in multicellular organismal process, protein binding, ion binding, regulation of cellular process, immune system and signaling by interleukins.

**Fig. 3.**
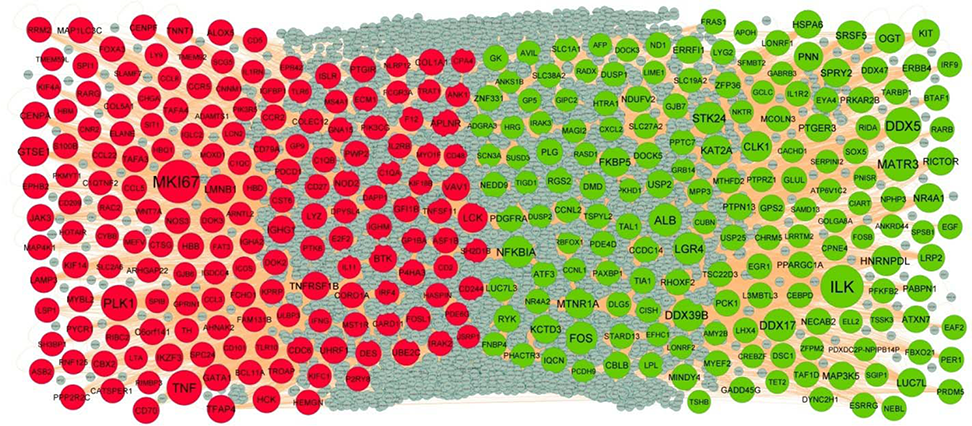
PPI network of DEGs. Up regulated genes are marked in green; down regulated genes are marked in red

**Fig. 4.**
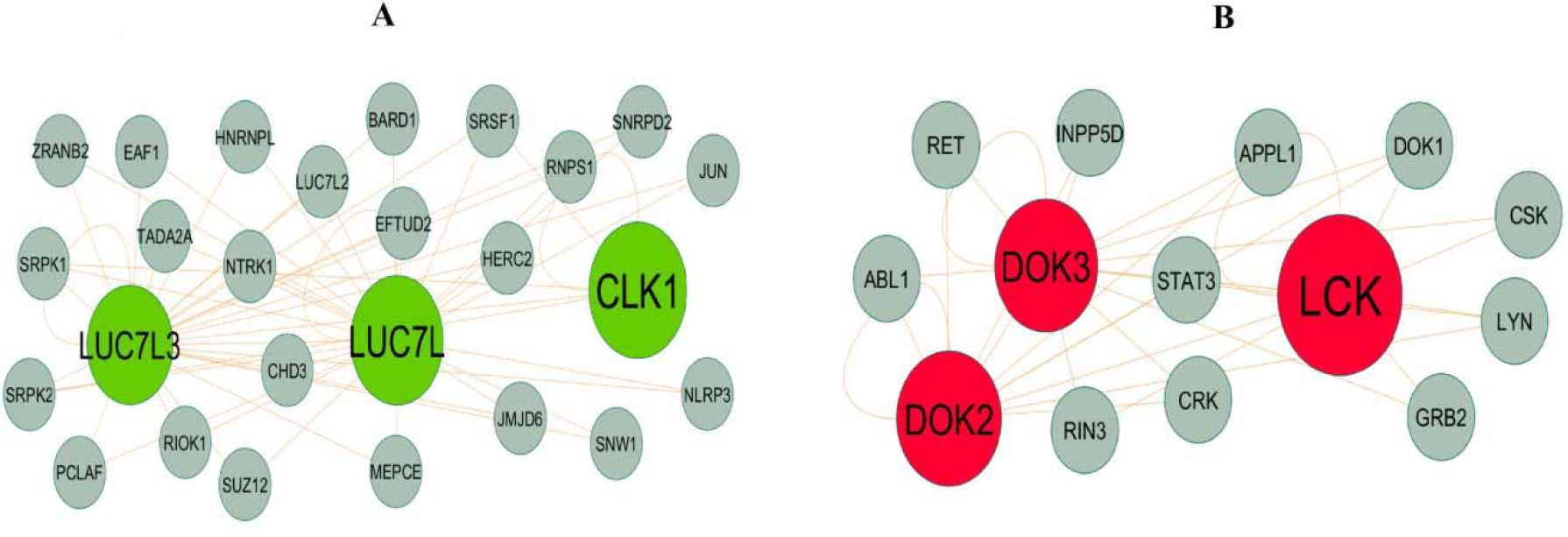
Modules selected from the DEG PPI between patients with FSGS and normal controls. (A) The most significant module was obtained from PPI network with 26 nodes and 54 edges for up regulated genes (B) The most significant module was obtained from PPI network with 14 nodes and 31 edges for down regulated genes. Up regulated genes are marked in green; down regulated genes are marked in red

**Table 4.**
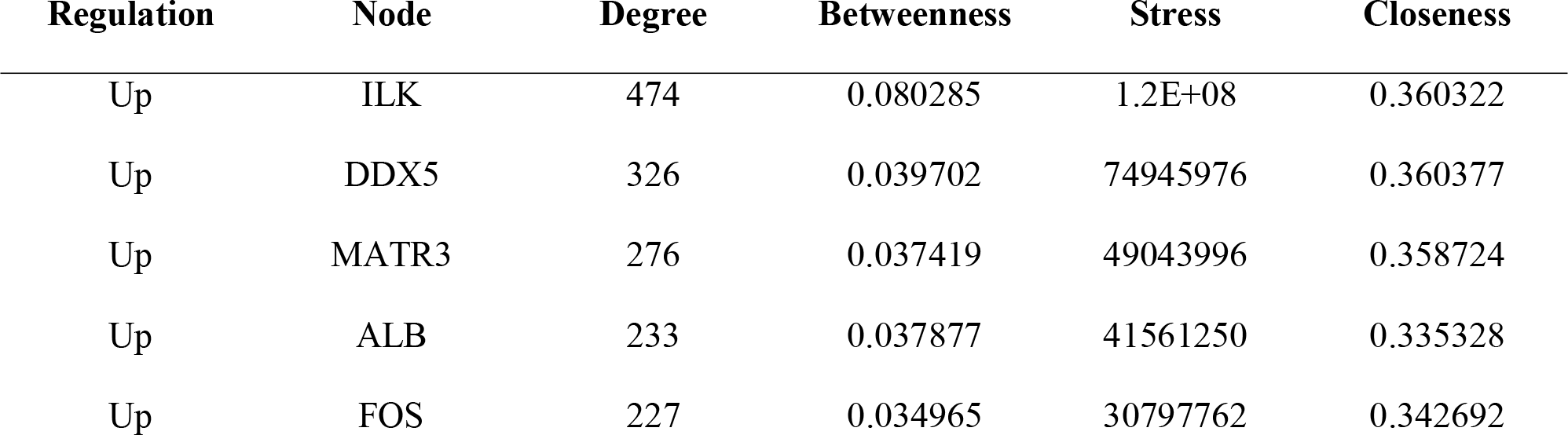
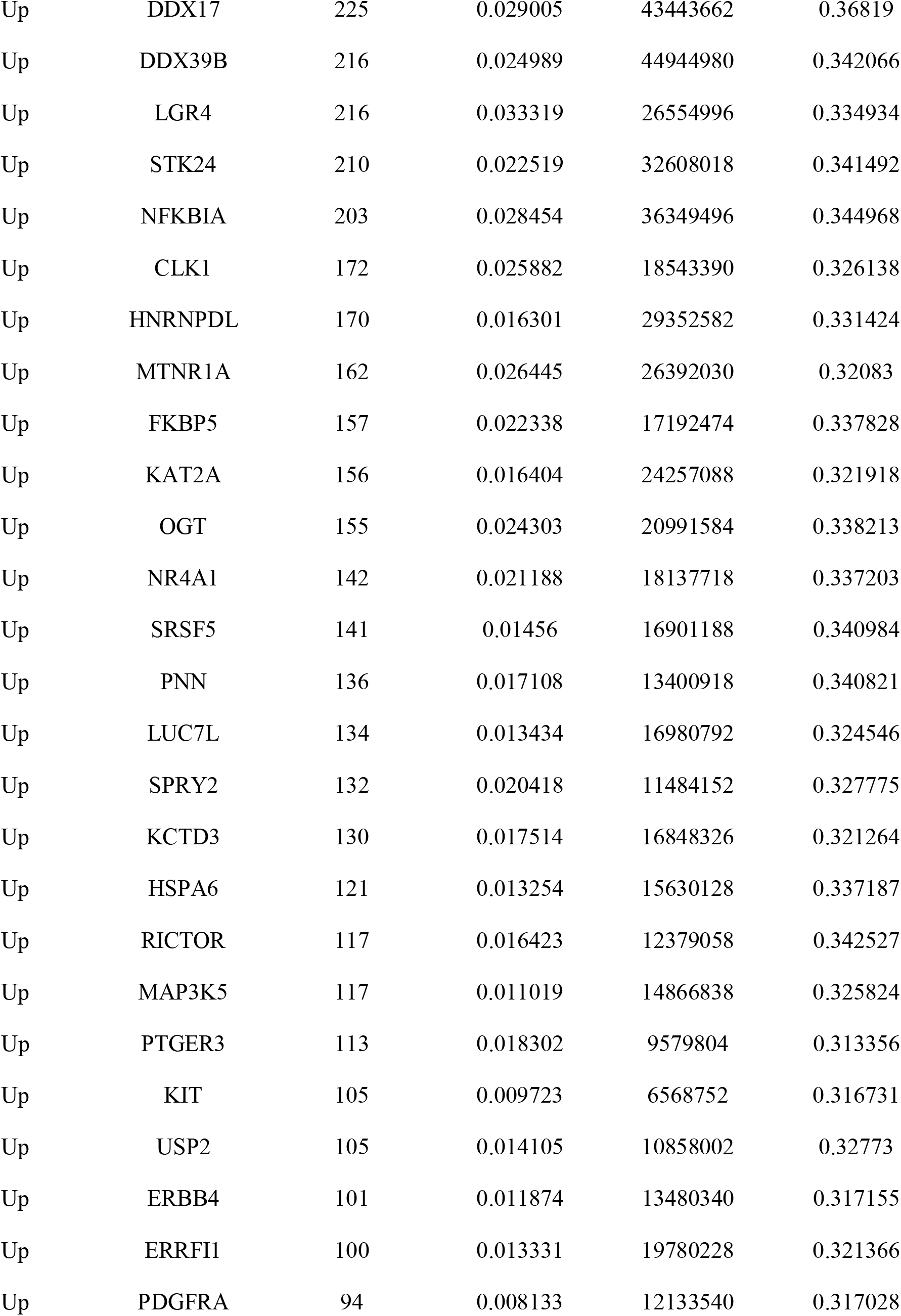
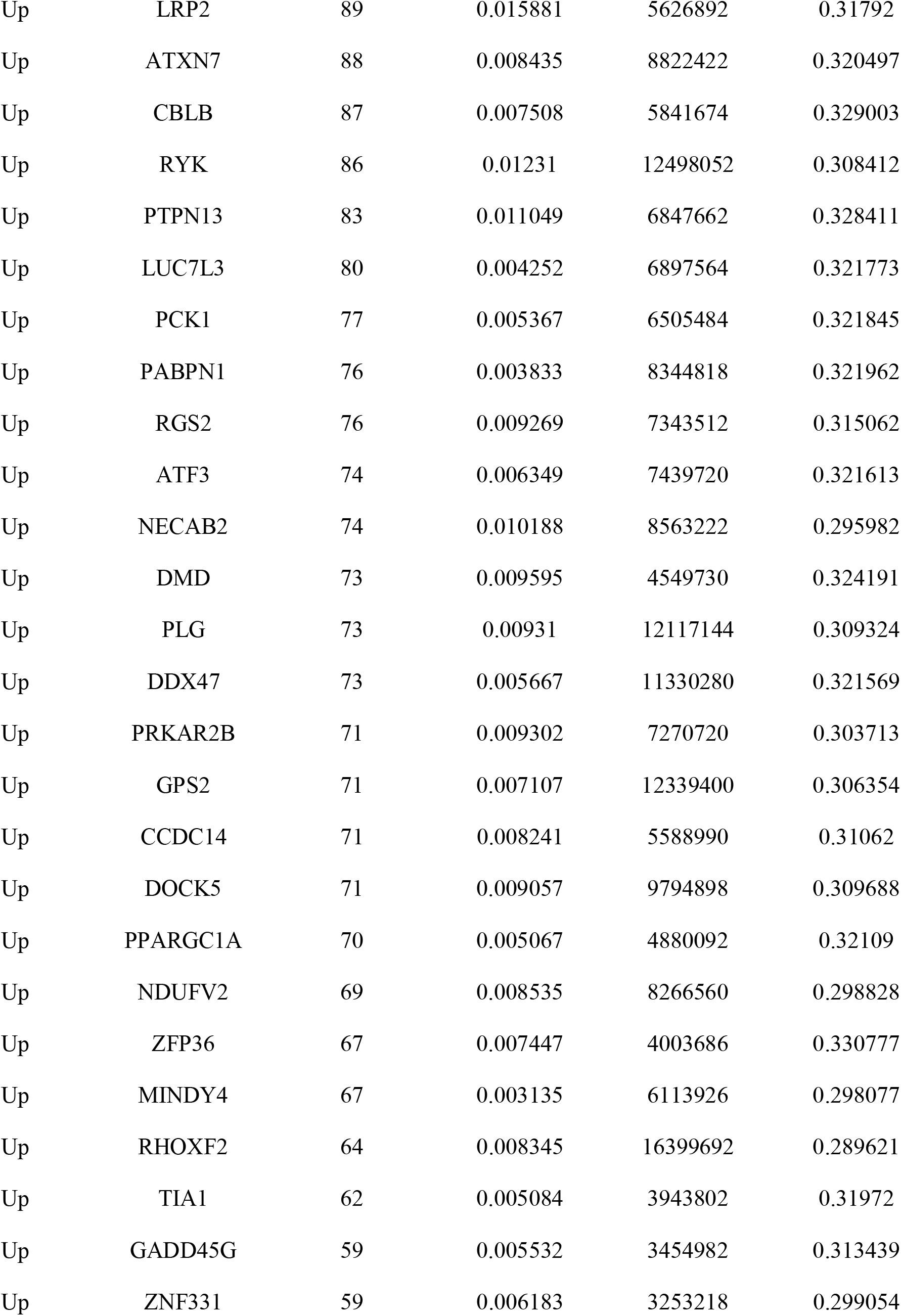
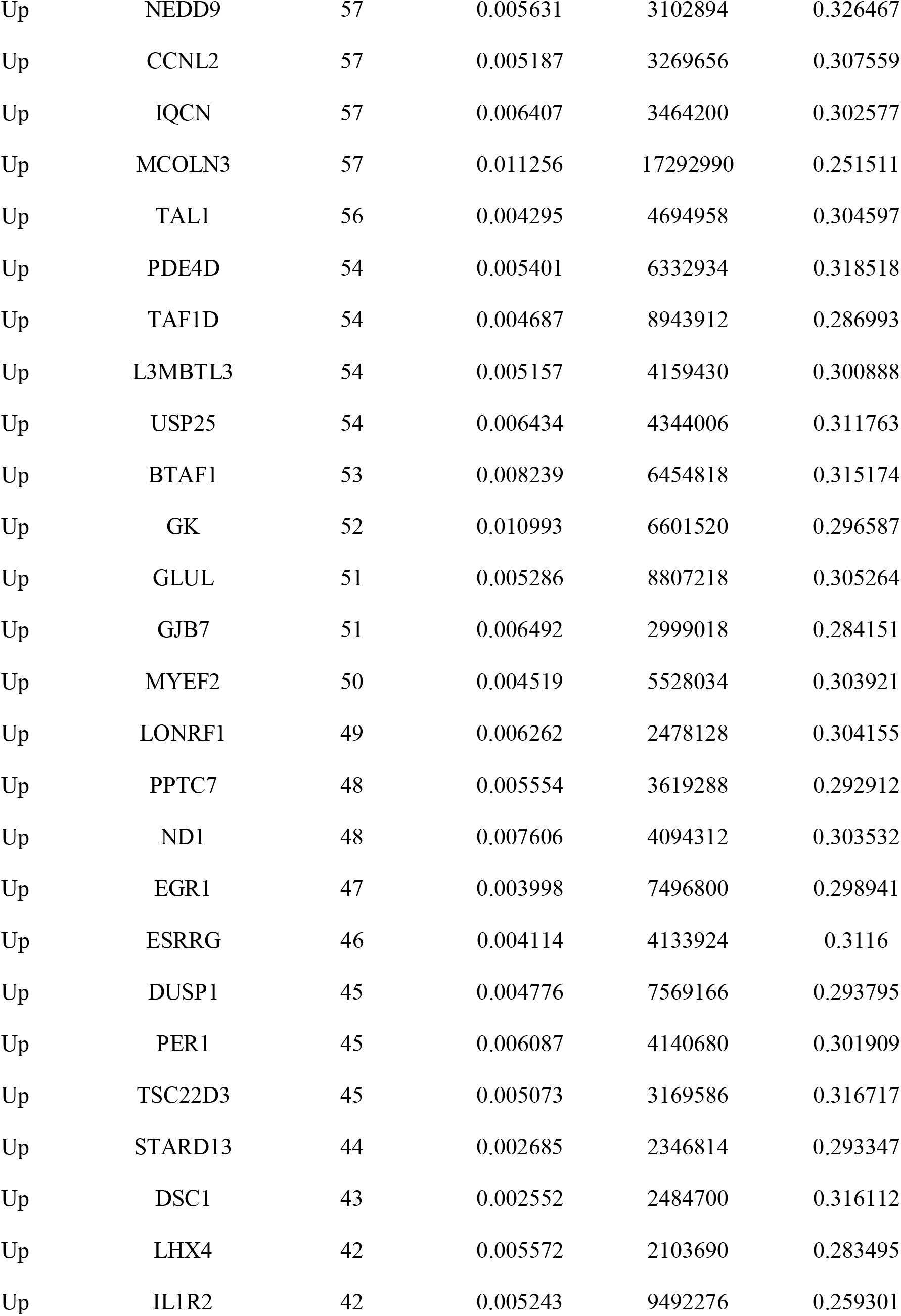
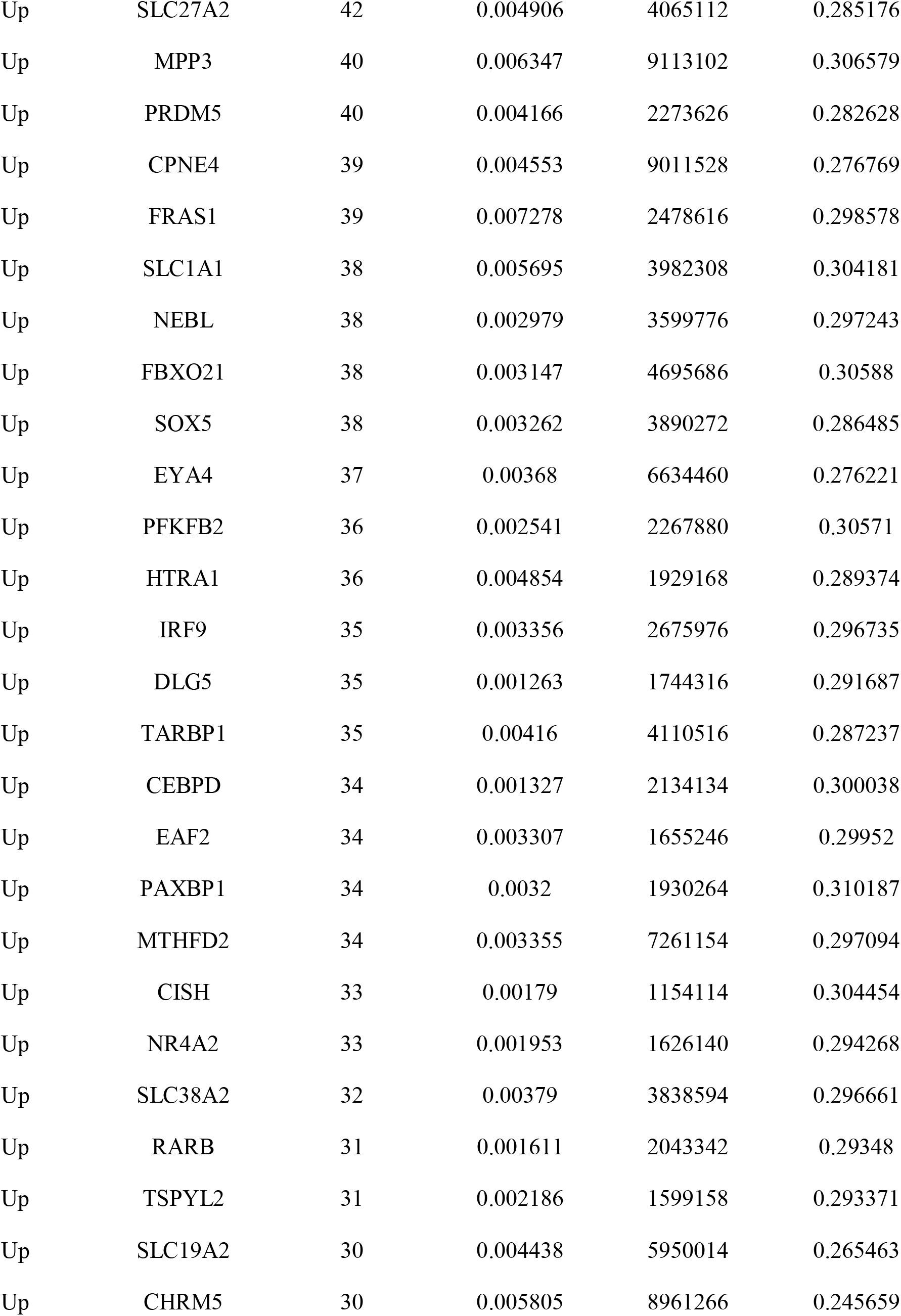
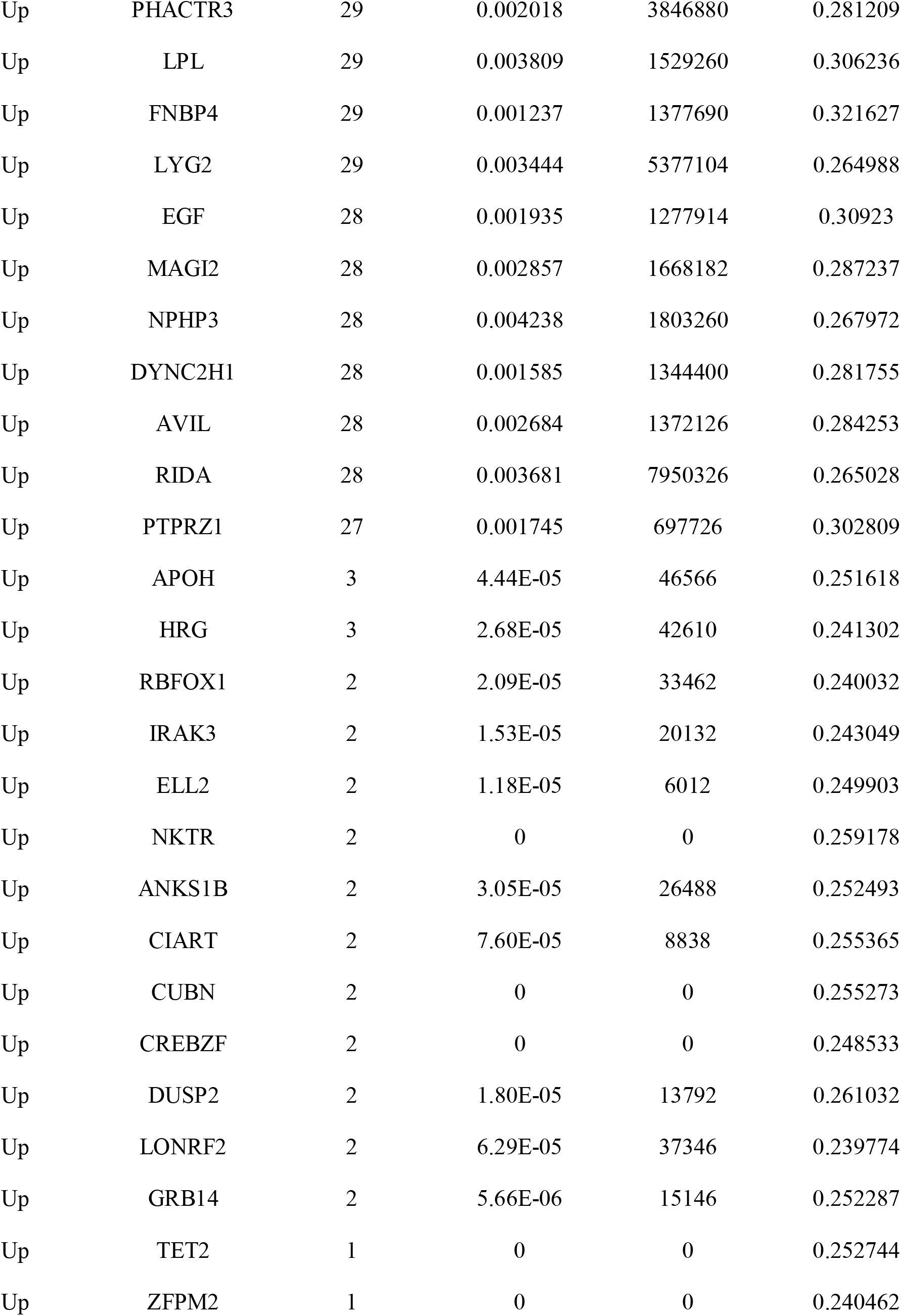
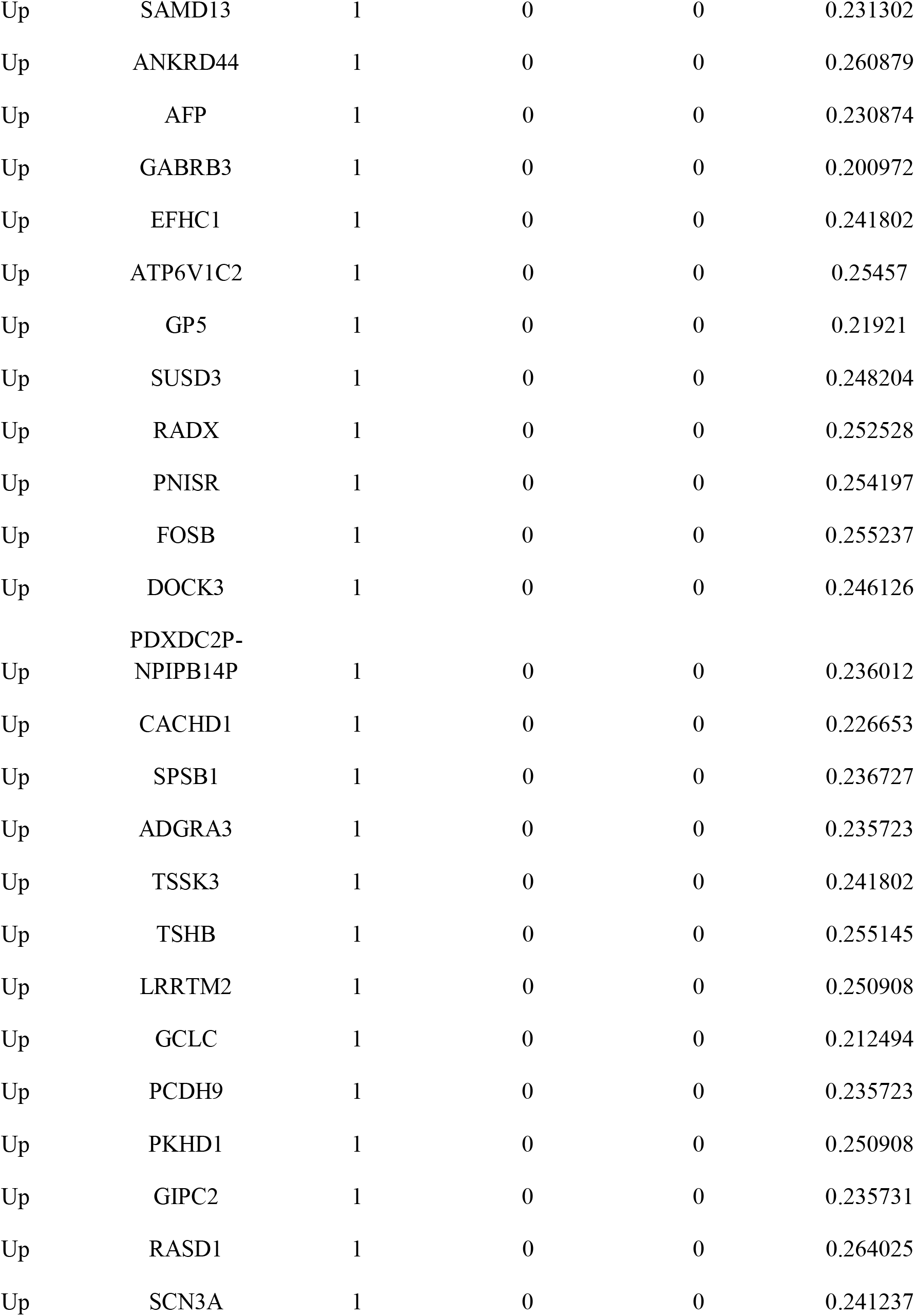
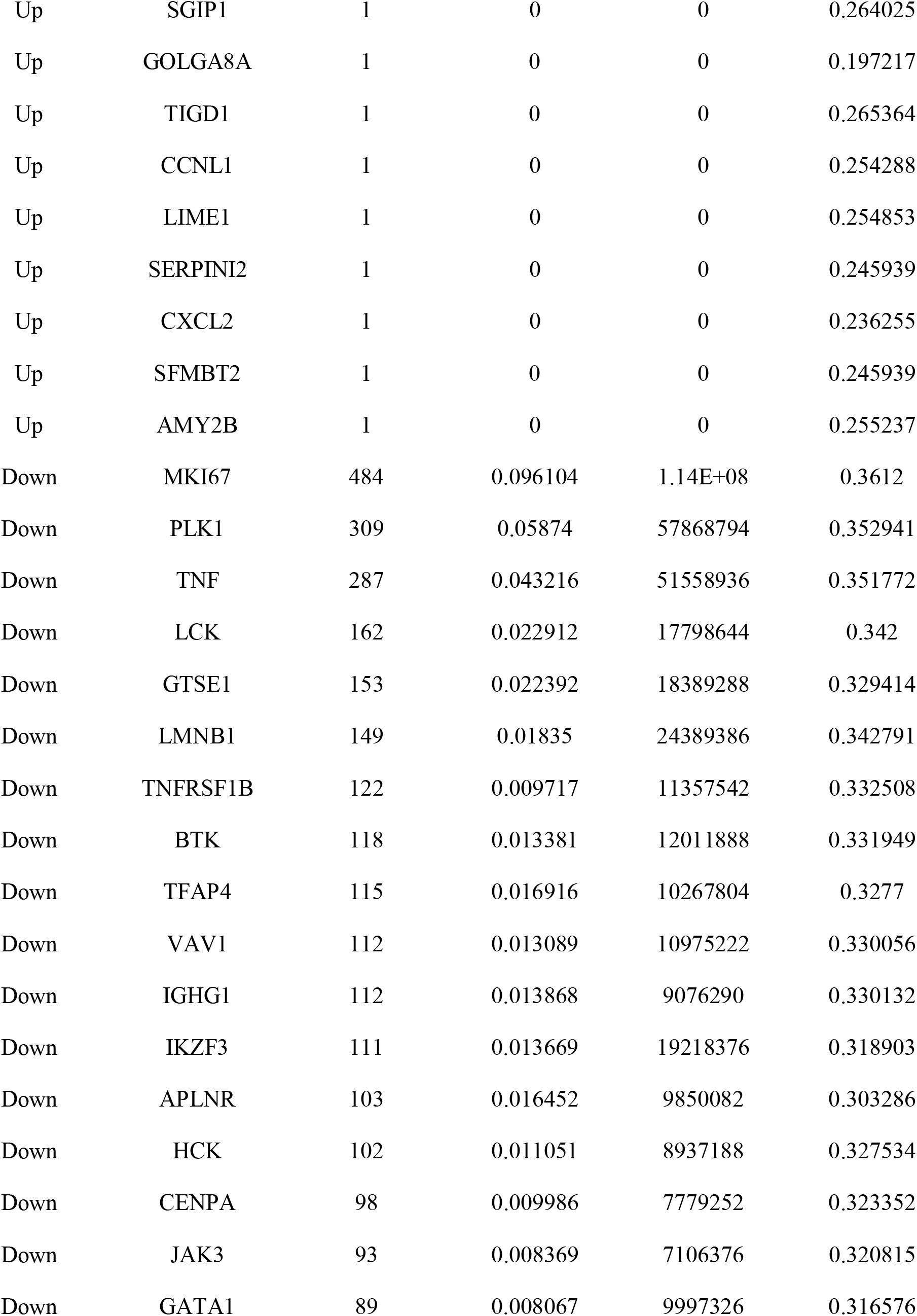
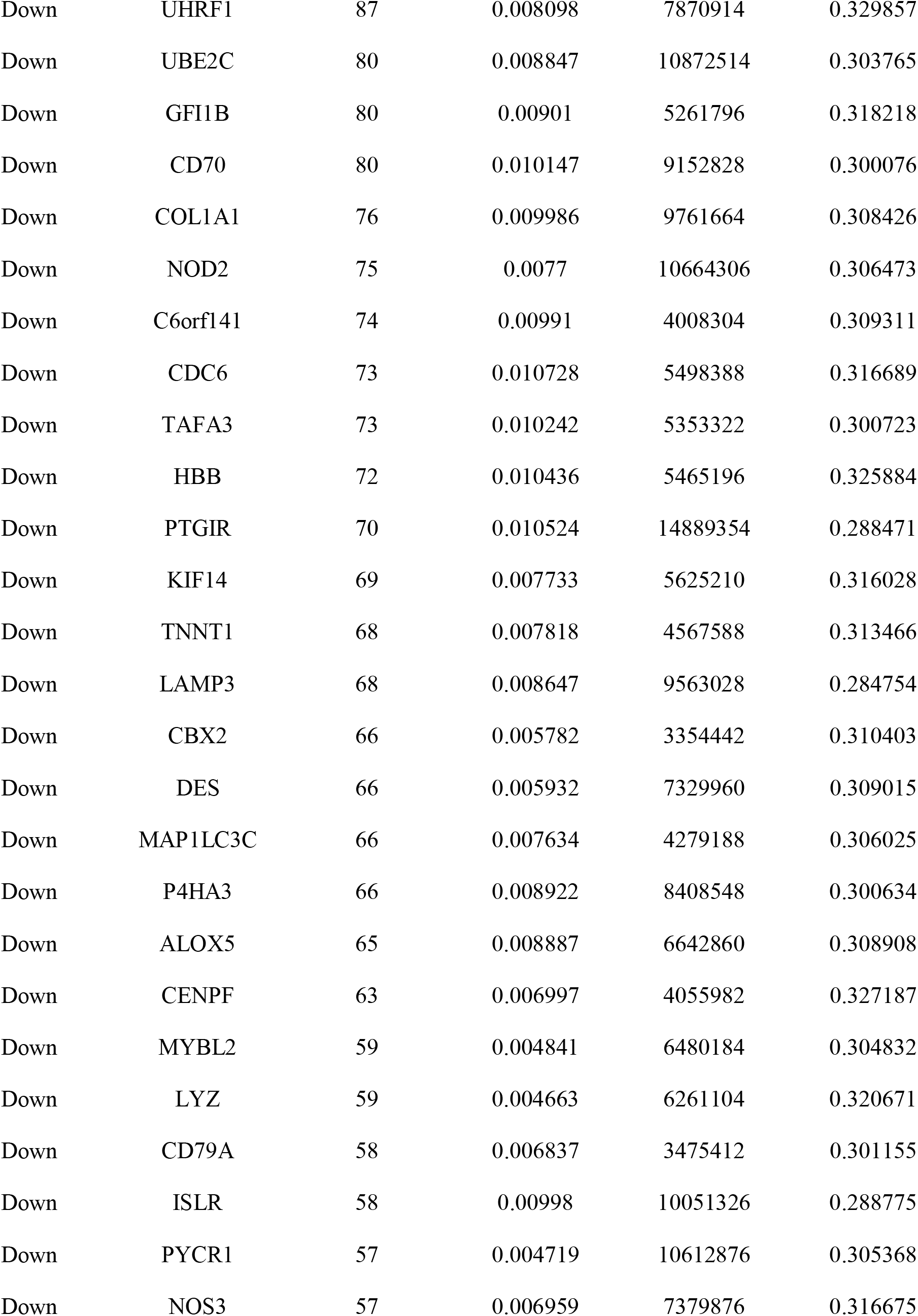
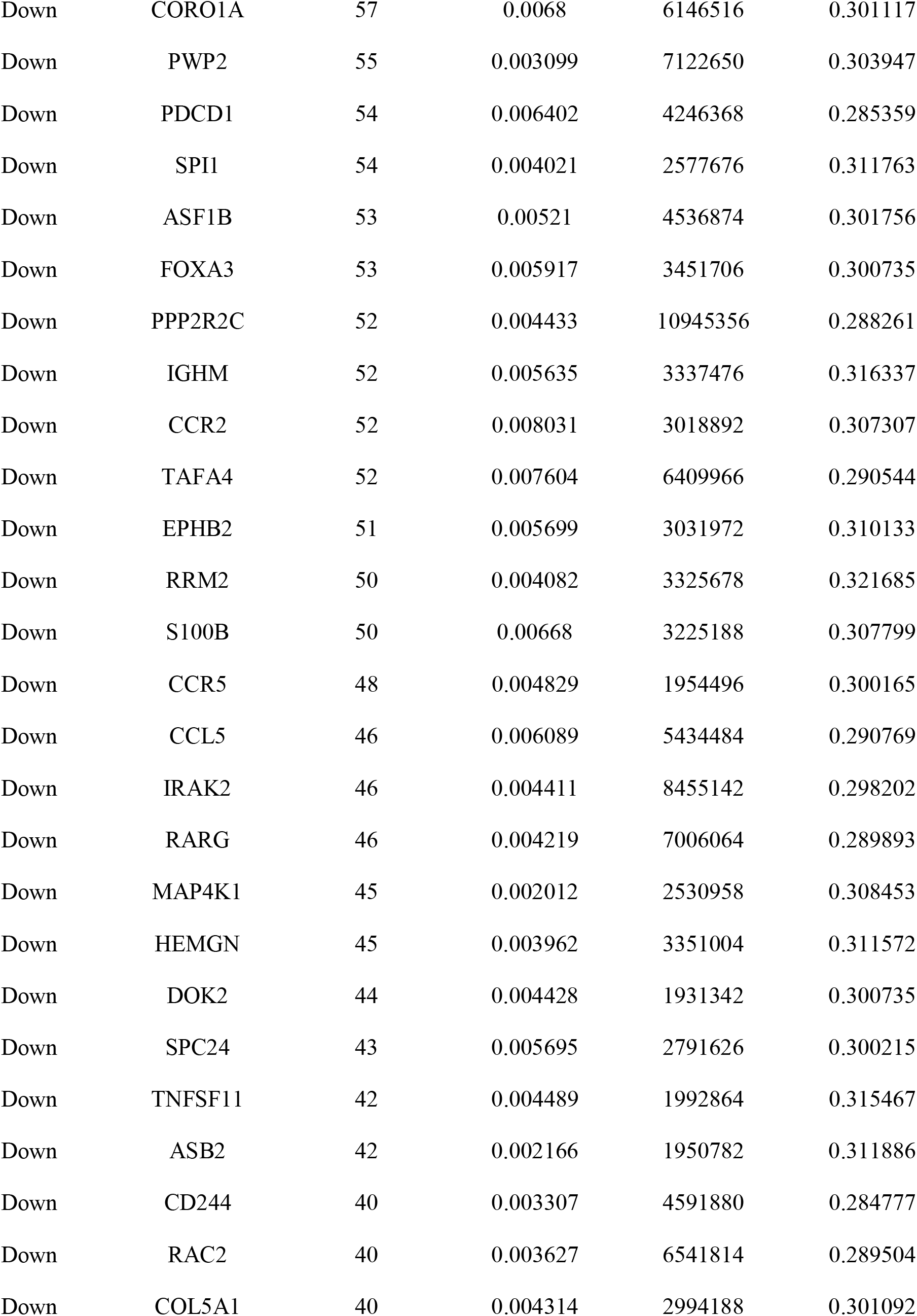
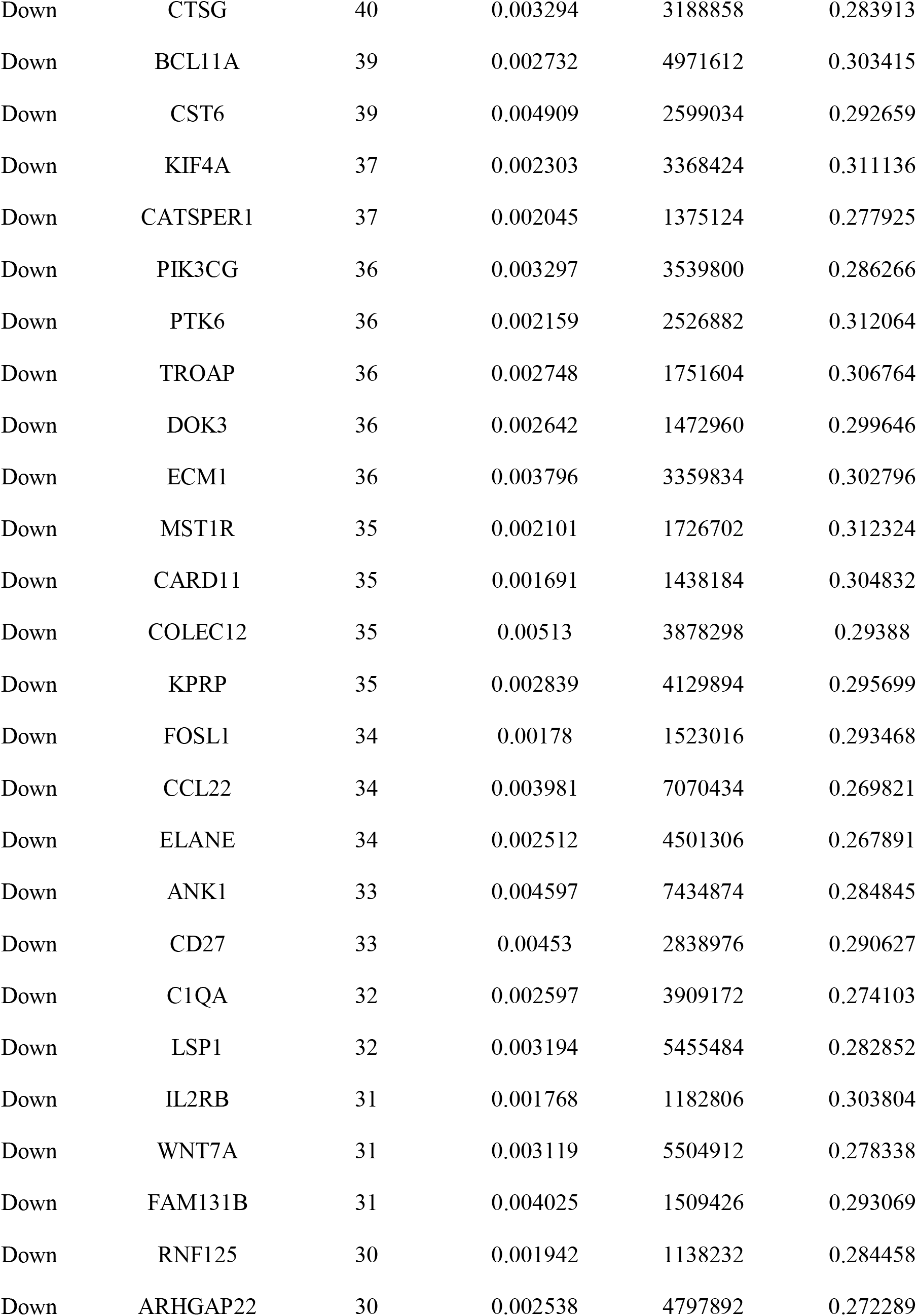
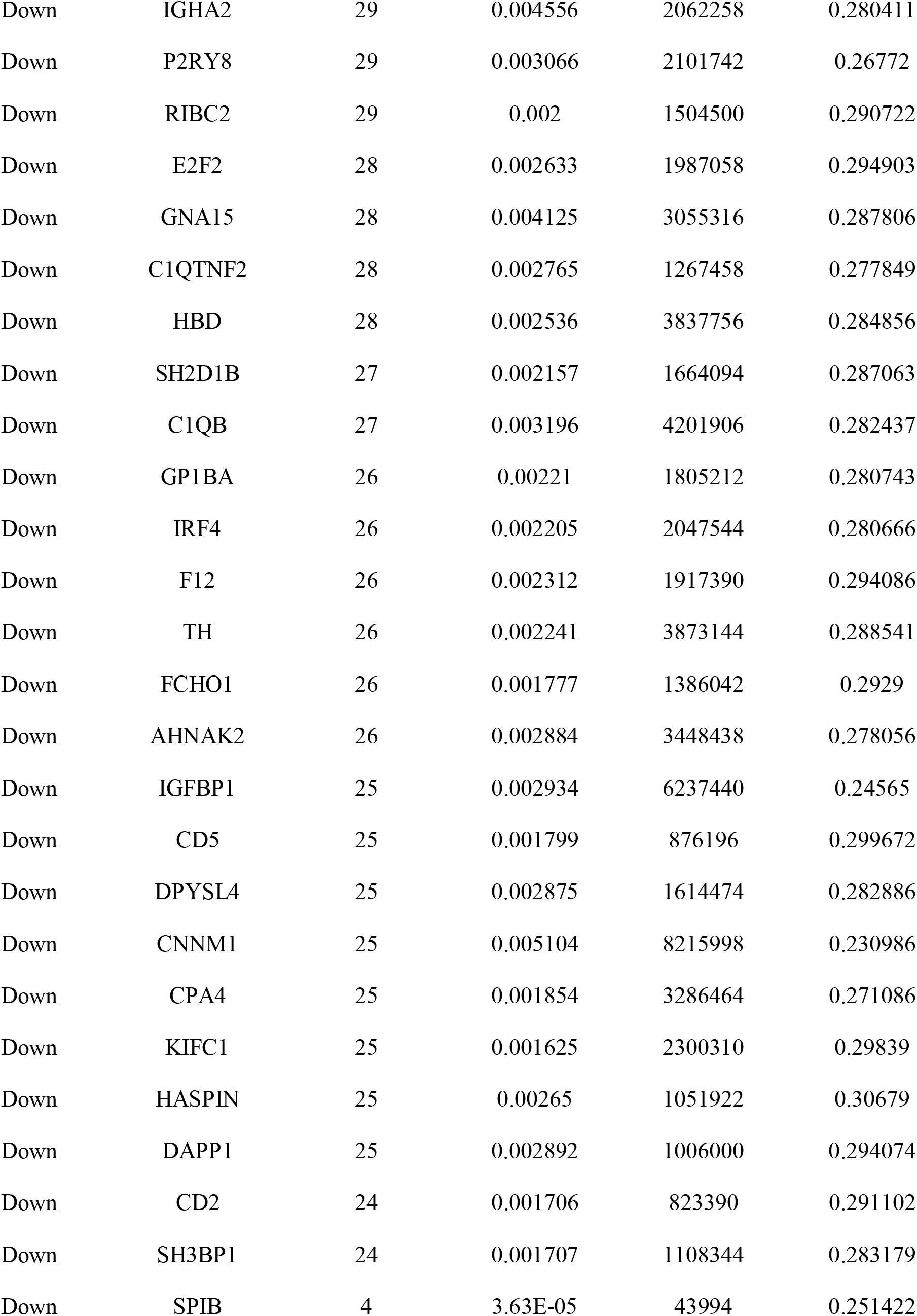
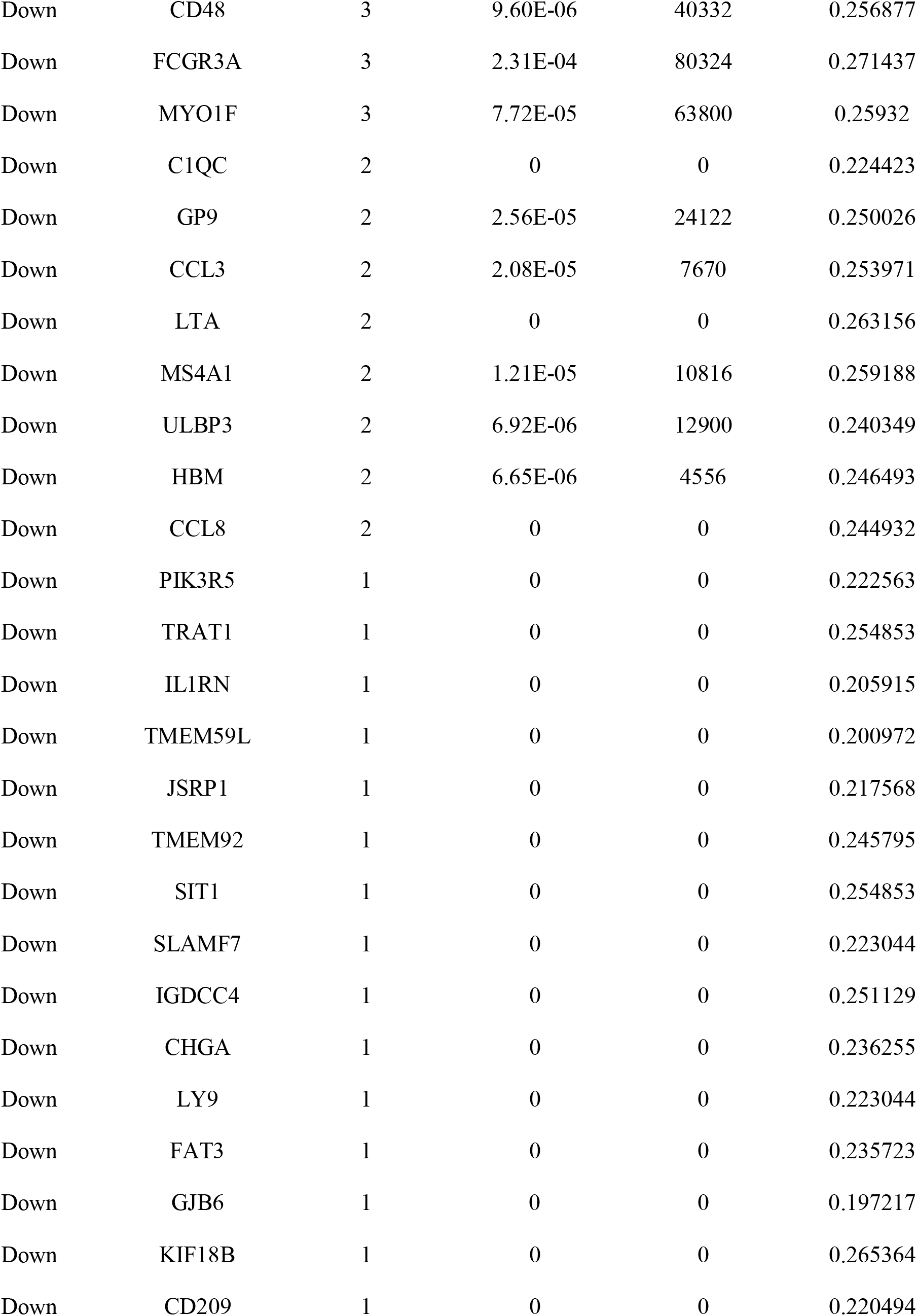
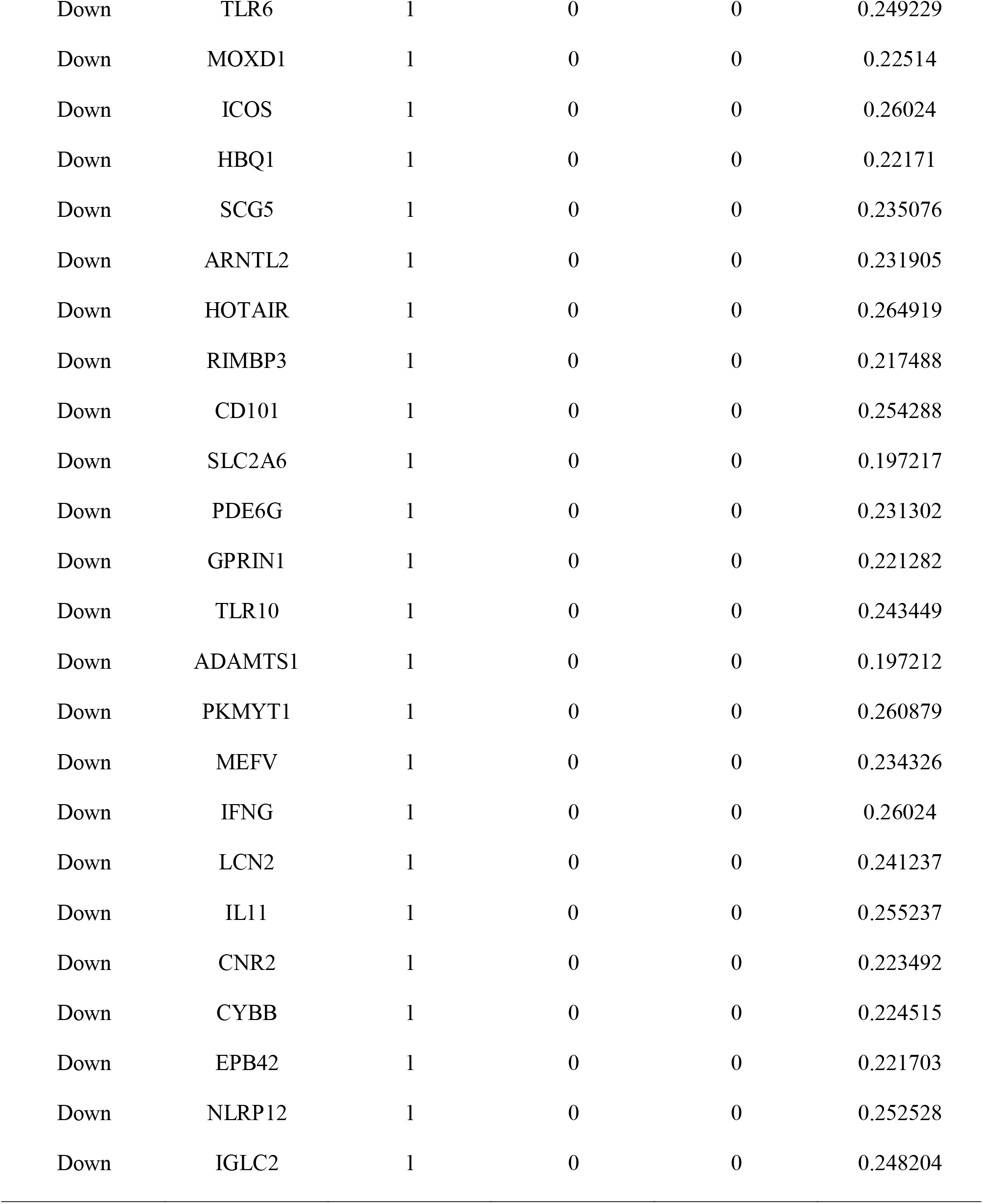
Topology table for up and down regulated genes

### miRNA-hub gene regulatory network construction

The hub gene expression products in T1DM were constructed into miRNA-hub gene regulatory network using the miRNet in Cytoscape software. miRNA-hub gene regulatory network was visualized, which involved 2482 nodes (328 genes and 2154 miRNAs) and 14789 edges (Fig. 5). Notably, a single genes might interact with different miRNAs. DDX17 was identified as targets of 192 miRNAs (ex; hsa-mir-503-5p); DDX5 was identified as targets of 155 miRNAs (ex; hsa-mir-500b-5p); MATR3 was identified as targets of 137 miRNAs (ex; hsa-mir-129-5p); DDX39B was identified as targets of 137 miRNAs (ex; hsa-mir-6079); FKBP5 was identified as targets of 116 miRNAs (ex; hsa-mir-4740-5p); MKI67 was identified as targets of 192 miRNAs (ex; hsa-mir-1231); LMNB1 was identified as targets of 157 miRNAs (ex; hsa-mir-190a-3p); IKZF3 was identified as targets of 147 miRNAs (ex; hsa-mir-1304-3p); TFAP4 was identified as targets of 95 miRNAs (ex; hsa-mir-520e); PLK1 was identified as targets of 94 miRNAs (ex; hsa-mir-21-5p) and are shown in Table 5. These findings revealed a complex regulatory network between miRNAs and hub genes in the process of FSGS.

**Fig. 5.**
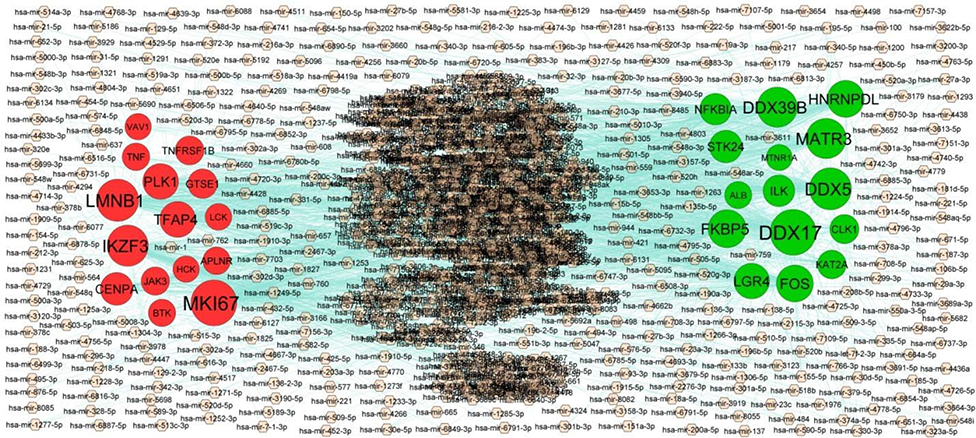
Target gene - miRNA regulatory network between target genes. The pinke color diamond nodes represent the key miRNAs; up regulated genes are marked in green; down regulated genes are marked in red.

**Table 5.** miRNA - target gene and TF - target gene interaction

| Regulation | Target Genes | Degree | MicroRNA | Regulation | Target Genes | Degree | TF |
| --- | --- | --- | --- | --- | --- | --- | --- |
| Up | DDX17 | 192 | hsa-mir-503-5p | Up | DDX39B | 56 | STAT3 |
| Up | DDX5 | 155 | hsa-mir-500b-5p | Up | FKBP5 | 17 | STAT1 |
| Up | MATR3 | 137 | hsa-mir-129-5p | Up | FOS | 15 | FOXD1 |
| Up | DDX39B | 137 | hsa-mir-6079 | Up | ILK | 12 | HOXA5 |
| Up | FKBP5 | 116 | hsa-mir-4740-5p | Up | HNRNPDL | 11 | GATA2 |
| Up | HNRNPDL | 105 | hsa-mir-550a-3-5p | Up | MATR3 | 10 | JUND |
| Up | FOS | 105 | hsa-mir-6077 | Up | CLK1 | 10 | FOXA1 |
| Up | LGR4 | 92 | hsa-mir-203a-3p | Up | STK24 | 8 | RUNX2 |
| Up | STK24 | 63 | hsa-mir-4511 | Up | DDX5 | 8 | BRCA1 |
| Up | NFKBIA | 54 | hsa-mir-548q | Up | NFKBIA | 8 | TFAP2C |
| Up | ILK | 54 | hsa-mir-1825 | Up | DDX17 | 7 | POU2F2 |
| Up | CLK1 | 30 | hsa-mir-218-2-3p | Up | KAT2A | 6 | FOXO3 |
| Up | KAT2A | 26 | hsa-mir-155-5p | Up | LGR4 | 5 | IRF2 |
| Up | ALB | 10 | hsa-mir-203a-3p | Up | MTNR1A | 4 | NKX3-2 |
| Up | MTNR1A | 1 | hsa-mir-31-5p | Up | ALB | 4 | YY1 |
| Down | MKI67 | 192 | hsa-mir-1231 | Down | TNF | 51 | GATA3 |
| Down | LMNB1 | 157 | hsa-mir-190a-3p | Down | LMNB1 | 12 | FOXF2 |
| Down | IKZF3 | 147 | hsa-mir-1304-3p | Down | VAV1 | 10 | JUN |
| Down | TFAP4 | 95 | hsa-mir-520e | Down | LCK | 10 | PAX2 |
| Down | PLK1 | 94 | hsa-mir-21-5p | Down | IKZF3 | 9 | REL |
| Down | CENPA | 66 | hsa-mir-6750-3p | Down | TNFRSF1B | 8 | USF2 |
| Down | GTSE1 | 33 | hsa-mir-320e | Down | PLK1 | 7 | SRY |
| Down | TNFRSF1B | 28 | hsa-mir-19a-3p | Down | CENPA | 7 | PRDM1 |
| Down | JAK3 | 22 | hsa-mir-6791-5p | Down | MKI67 | 7 | RELA |
| Down | APLNR | 15 | hsa-mir-146a-5p | Down | TFAP4 | 7 | ESR1 |
| Down | TNF | 15 | hsa-mir-187-3p | Down | HCK | 5 | TP53 |
| Down | BTK | 13 | hsa-mir-4269 | Down | GTSE1 | 4 | TFAP2A |
| Down | LCK | 6 | hsa-mir-210-3p | Down | APLNR | 4 | TP63 |
| Down | VAV1 | 4 | hsa-mir-27a-3p | Down | JAK3 | 2 | FEV |
| Down | HCK | 3 | hsa-mir-520f-3p | Down | BTK | 2 | SOX5 |

### TF-hub gene regulatory network construction

The hub gene expression products in T1DM were constructed into TF-hub gene regulatory network using the NetworkAnalyst in Cytoscape software. TF-hub gene regulatory network was visualized, which involved 421 nodes (325 genes and 96 TFs) and 2714 edges (Fig. 6). Notably, a single genes might interact with different TFs. DDX39B was identified as targets of 56 TFs (ex; STAT3); FKBP5 was identified as targets of 17 TFs (ex; STAT1); FOS was identified as targets of 15 TFs (ex; FOXD1); ILK was identified as targets of 12 TFs (ex; HOXA5); HNRNPDL was identified as targets of 11 TFs (ex; GATA2); TNF was identified as targets of 51 TFs (ex; GATA3); LMNB1 was identified as targets of 12 TFs (ex; FOXF2); VAV1 was identified as targets of 10 TFs (ex; JUN); LCK was identified as targets of 10 TFs (ex; PAX2); IKZF3 was identified as targets of 9 TFs (ex; REL) and are shown in Table 5. These findings revealed a complex regulatory network between TFs and hub genes in the process of FSGS.

**Fig. 6.**
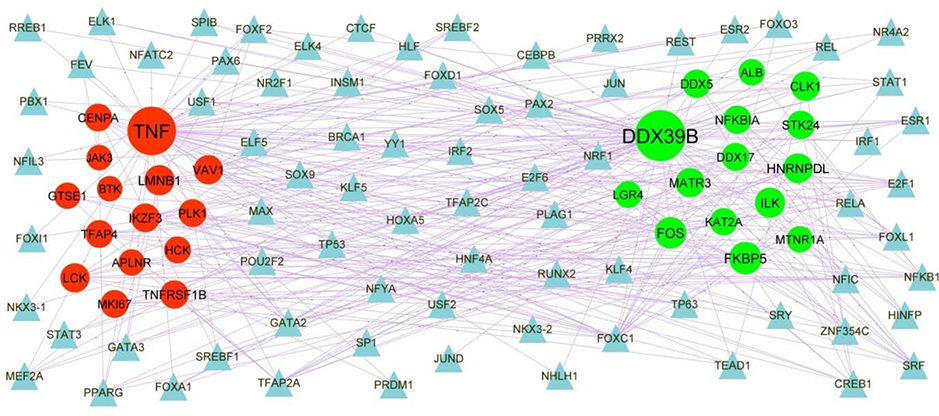
Target gene - TF regulatory network between target genes. The blue color triangle nodes represent the key TFs; up regulated genes are marked in green; down regulated genes are marked in red.

### Receiver operating characteristic curve (ROC) analysis

The ROC curve was used to assess the predictive accuracy of hub genes. ROC curves and AUC values are shown in Fig. 7. All AUC values exceeded 0.9, while the up regulated hub genes ILK, DDX5, MATR3, ALB and FOS, and down regulated hub genes MKI67, PLK1, TNF, LCK and GTSE1 had AUC values >0.9. The ROC analysis revealed that the expression levels of these genes had excellent predictive performance and were able to discriminate FSGS from normal control.

**Fig. 7.**
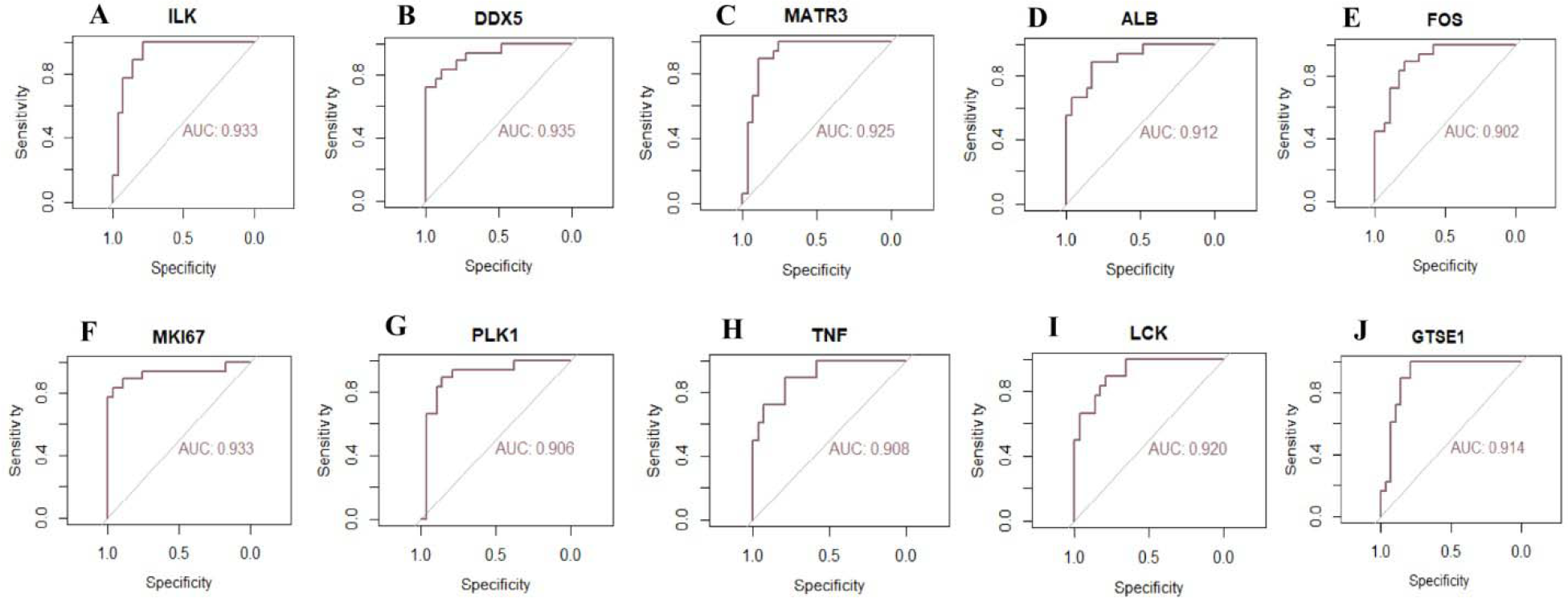
ROC curve analyses of hub genes. A) ILK B) DDX5 C) MATR3 D) ALB E) FOS F) MKI67 G) PLK1 H) TNF I) LCK J) GTSE1

## Discussion

FSGS is a glomerular injury specific to kidney, and deficiency in our knowledge of the exact etiology and pathogenesis of FSGS restricts the ability to treat this disease. Therefore, understanding the molecular mechanism associated in FSGS is extremely key to promote more effective diagnostic, prognostics and therapeutic strategies. A NGS study is an ideal way to comprehensively investigate FSGS. In this investigation, NGS data were comprehensively analyzed using bioinformatics analysis to identify FSGS related genes and disease mechanisms. A total of 976 DEGs (488 up regulated and 488 down regulated) were identified. Roth and Berntorp [36] found that the GNRH1 gene is involved in the pathogenesis of diabetes mellitus. Investigation have shown that elevated OMG (oligodendrocyte myelin glycoprotein) [37], IGHV4-39 [38] and CCL18 [39] levels are associated with an increased risk of multiple sclerosis. Altered expression of CCL18 [40] is associated with progression of cardiovascular diseases. CCL18 is reported to play a central role in polycystic ovarian syndrome [41].

The GO and REACTOME pathway database contains information on systematic analysis of gene functions, linking genomics with functional information. Pathways include metabolism [42], transport of small molecules [43] and immune system [44] are linked with progression of FSGS. PDK4 [45], ALB (albumin) [46], EGR1 [47], RYR3 [48], APOH (apolipoprotein H) [49], CYP27B1 [50], ESM1 [51], GATA6 [52], PCK1 [53], TET2 [54], AFP (alpha fetoprotein) [55], CACNA1D [56], ATF3 [57], EGF (epidermal growth factor) [58], LPL (lipoprotein lipase) [59], PPARGC1A [60], PLG (plasminogen) [61], NR4A1 [62], STRA6 [63], MLXIPL (MLX interacting protein like) [64], PLA2G6 [65], RGS2 [66], GPS2 [67], SOX5 [68], GLUL (glutamate-ammonia ligase) [69], RYK (receptor like tyrosine kinase) [70], NFKBIA (NFKB inhibitor alpha) [71], LGR4 [72], SPRY2 [73], TRPC1 [74], KL (klotho) [75], GCLC (glutamate-cysteine ligase catalytic subunit) [76], NOX4 [77], CD69 [78], SLC19A2 [79], S100A12 [80], MT1A [81], SORCS1 [82], FKBP5 [83], AFM (afamin) [84], CA3 [85], MAOA (monoamine oxidase A) [86], ND1 [87], ATP6 [88], CHGA (chromogranin A) [89], ICOS (inducible T cell costimulator) [90], DBH (dopamine beta-hydroxylase) [91], CD5 [92], LTA (lymphotoxin alpha) [93], IFNG (interferon gamma) [94], MPO (myeloperoxidase) [95], CD70 [96], CD300E [97], COLEC12 [98], TLR10 [99], LCN2 [100], SLAMF7 [101], TREM2 [102], ITGAL (integrin subunit alpha L) [103], CD27 [104], JAK3 [105], CCR5 [106], FCN1 [107], IL1RN [108], CX3CR1 [109], PDCD1 [110], TRPM2 [111], PLEK (pleckstrin) [112], CD101 [113], TNF (tumor necrosis factor) [114], CD48 [115], ALOX5 [116], TLR7 [117], CCL3 [118], C2 [119], TNFRSF1B [120], CCR2 [121], PLA2G7 [122], TH (tyrosine hydroxylase) [123], WNT7A [124], ADRB3 [125], GPBAR1 [126], SLC6A20 [127], FUT2 [128], ANK1 [129], NOS3 [130], APLNR (apelin receptor) [131], COMP (cartilage oligomeric matrix protein) [132], RETN (resistin) [133], NMU (neuromedin U) [134], S100B [135], IGFBP1 [136], COL1A1 [137], HBB (hemoglobin subunit beta) [138] and PLAC8 [139] genes plays important regulatory roles in diabetes mellitus. Various genes such as PDK4 [140], ALB (albumin) [141], EGR1 [142], RYR3 [48], CYP27B1 [143], GATA6 [144], NR4A3 [145], TET2 [146], CACNA1D [147], KLF15 [148], CYP4A11 [149], ATOH8 [150], ATF3 [151], EGF (epidermal growth factor) [152], LPL (lipoprotein lipase) [153], PPARGC1A [154], PLG (plasminogen) [155], USP2 [156], PDE4D [157], RGS2 [158], ILK (integrin linked kinase) [159], SLC26A4 [160], PER1 [161], LGR4 [72], GABRB3 [162], TRPC1 [163], KL (klotho) [164], NOX4 [165], S100A12 [166], NEDD9 [167], ANKRD36 [168], IRF9 [169], GIMAP5 [170], ND2 [171], CHGA (chromogranin A) [172], DBH (dopamine beta-hydroxylase) [173], LTA (lymphotoxin alpha) [174], MPO (myeloperoxidase) [175], CD70 [176], LCN2 [177], TLR6 [178], TREM2 [179], NLRP12 [180], CCR5 [181], IL1RN [182], CX3CR1 [183], TRPM2 [184], TNF (tumor necrosis factor) [185], TLR7 [186], NOD2 [187], TNFRSF1B [188], CCR2 [181], CCL5 [189], PLA2G7 [190], TH (tyrosine hydroxylase) [191], WNT7A [192], ADRB3 [193], IL10RA [194], NOS3 [195], PIK3R5 [196], APLNR (apelin receptor) [197], COMP (cartilage oligomeric matrix protein) [198], RETN (resistin) [199], S100B [200], IGFBP1 [201] and COL1A1 [202] are crucial in the progression of hypertension. Evidence suggests that PDK4 [203], ALB (albumin) [204], EGR1 [205], RYR3 [206], DUSP1 [207], EYA4 [208], CYP27B1 [143], ESM1 [51], GATA6 [209], NR4A3 [210], TET2 [211], KLF15 [212], CYP4A11 [213], ATF3 [214], DMD (dystrophin) [215], LPL (lipoprotein lipase) [216], PPARGC1A [154], NPHP3 [217], TNNI3K [218], PLG (plasminogen) [219], NR4A2 [220], NR4A1 [221], ARHGEF10 [222], STRA6 [223], MLXIPL (MLX interacting protein like) [224], PDE4D [225], CUBN (cubilin) [226], OGT (O-linked N-acetylglucosamine (GlcNAc) transferase) [227], RGS2 [228], SOX5 [229], ILK (integrin linked kinase) [230], SLC26A4 [231], GLUL (glutamate-ammonia ligase) [69], RBFOX1 [232], HSPA1B [233], ZFP36 [234], DDIT4 [235], NFKBIA (NFKB inhibitor alpha) [71], TRPC1 [236], KL (klotho) [237], GCLC (glutamate-cysteine ligase catalytic subunit) [238], NOX4 [239], CD69 [240], S100A12 [241], KLF9 [242], MT1A [81], FKBP5 [243], CA3 [85], IRF9 [244], LUC7L3 [245], CXCL2 [246], PFKFB2 [247], ND1 [87], ATP6 [248], CCL22 [249], SAA1 [250], CD5L [251], CHGA (chromogranin A) [252], CNR2 [253], DBH (dopamine beta-hydroxylase) [254], CTSG (cathepsin G) [255], SFRP2 [256], MEFV (MEFV innate immuity regulator, pyrin) [257], DEFA3 [258], ELANE (elastase, neutrophil expressed) [259], LTA (lymphotoxin alpha) [260], IFNG (interferon gamma) [261], MPO (myeloperoxidase) [262], CD70 [263], IRF4 [264], LCN2 [265], TLR6 [266], SLAMF7 [101], TREM2 [267], JAK3 [268], NLRP12 [269], CCR5 [270], FCN1 [271], NFAM1 [272], SPI1 [273], IL1RN [274], CX3CR1 [109], CYBB (cytochrome b-245 beta chain) [275], IL11 [276], TRPM2 [277], TNF (tumor necrosis factor) [278], ECM1 [279], SEMA7A [280], ALOX5 [281], TLR7 [282], CCL3 [118], NOD2 [283], PIK3CG [284], RAC2 [285], CCR2 [286], CCL5 [287], PLA2G7 [288], LAMP3 [289], MLC1 [290], KCND2 [291], CAVIN4 [292], ADRB3 [293], SIGLEC1 [294], STEAP4 [295], KCNE5 [296], RGS4 [297], NOS3 [298], IL2RB [299], MYO1F [300], APLNR (apelin receptor) [197], TNFRSF12A [301], COMP (cartilage oligomeric matrix protein) [302], RETN (resistin) [303], S100B [304], IGFBP1 [305], CTHRC1 [306], CYTL1 [307] and COL1A1 [202] have a strong role in progression of cardiovascular diseases. PDK4 [308], EGR1 [309], CEBPD (CCAAT enhancer binding protein delta) [310], DUSP1 [311], DUSP2 [312], CYP27B1 [313], NR4A3 [314], PCK1 [53], TET2 [315], CYP8B1 [316], KLF15 [317], ATF3 [318], EGF (epidermal growth factor) [319], LPL (lipoprotein lipase) [320], PPARGC1A [154], NR4A1 [321], STRA6 [322], USP2 [323], ESRRG (estrogen related receptor gamma) [324], GPS2 [67], ILK (integrin linked kinase) [325], DOCK5 [326], ZFPM2 [327], ZFP36 [328], STK24 [329], LGR4 [330], TRPC1 [331], IRAK3 [332], LRP2 [333], NOX4 [334], CD69 [335], MAP3K5 [336], FKBP5 [337], CA3 [338], MAOA (monoamine oxidase A) [339], ND2 [340], GK (glycerol kinase) [341], UGT2B7 [342], CYTB (cytochrome b) [343], SAA1 [344], CD5L [345], CHGA (chromogranin A) [346], CNR2 [347], ELANE (elastase, neutrophil expressed) [348], LTA (lymphotoxin alpha) [93], MPO (myeloperoxidase) [95], IRF4 [349], LYZ (lysozyme) [350], TLR10 [99], LCN2 [100], TLR6 [351], TREM2 [352], CD27 [104], NLRP12 [353], CCR5 [354], IL1RN [355], CX3CR1 [356], TNF (tumor necrosis factor) [357], PTAFR (platelet activating factor receptor) [358], TLR7 [359], NOD2 [360], CCR2 [361], CCL5 [354], TH (tyrosine hydroxylase) [123], WNT7A [124], ADRB3 [362], GPBAR1 [363], STEAP4 [364], SLC38A5 [365], RGS4 [366], SLC37A2 [367], NOS3 [368], SIGLEC7 [369], ELOVL3 [370], APLNR (apelin receptor) [131], COMP (cartilage oligomeric matrix protein) [371], RETN (resistin) [133], NMU (neuromedin U) [134], S100B [372], IGFBP1 [373] and PLAC8 [139] have been shown to play a role in obesity development. A recent investigation identified PDK4 [374], ALB (albumin) [375], CYP27B1 [376], PCK1 [377], TET2 [378], EGF (epidermal growth factor) [379], LPL (lipoprotein lipase) [380], PLG (plasminogen) [381], NR4A2 [382], SLC1A1 [383], OGT (O-linked N-acetylglucosamine (GlcNAc) transferase) [384], ERBB4 [385], L3MBTL3 [386], NAPEPLD (N-acyl phosphatidylethanolamine phospholipase D) [387], HTRA1 [388], CBLB (Cbl proto-oncogene B) [389], PDGFRA (platelet derived growth factor receptor alpha) [390], KL (klotho) [391], LRP2 [392], BTLA (B and T lymphocyte associated) [393], RGS1 [394], S100A12 [395], ND2 [396], CCL22 [397], ICOS (inducible T cell costimulator) [398], CNR2 [399], DBH (dopamine beta-hydroxylase) [400], CD5 [401], LTA (lymphotoxin alpha) [402], IFNG (interferon gamma) [403], MPO (myeloperoxidase) [404], CD70 [405], IRF4 [406], VAV1 [407], IKZF3 [408], BTK (Bruton tyrosine kinase) [409], LCN2 [410], TREM2 [411], CD27 [412], F12 [413], CCR5 [414], CX3CR1 [415], IL21R [416], CYBB (cytochrome b-245 beta chain) [417], IL11 [418], TRPM2 [419], EOMES (eomesodermin) [420], TNF (tumor necrosis factor) [421], CD2 [422], SEMA7A [423], TLR7 [424], CCL3 [425], NOD2 [426], CCR2 [427], CCL5 [428], TH (tyrosine hydroxylase) [429], SIGLEC1 [430], HBD (hemoglobin subunit delta) [431], NOS3 [432], SIGLEC7 [433], IL2RB [434], RETN (resistin) [435] and S100B [436] as major genes contributing to multiple sclerosis. Musante et al. [437], Han et al. [438], Zhang et al. [439], Sato et al. [440], Yang et al. [441], El-Aouni et al. [442], Perisic et al. [443], Chung et al. [444], Wilkening et al. [445], Kahvecioglu et al. [446], Hu et al. [447] and Worthmann et al. [448] revealed that ALB (albumin), KLF15, ATF3, LPL (lipoprotein lipase), CUBN (cubilin), ILK (integrin linked kinase), PLEKHH2, TNF (tumor necrosis factor), CCR2, SPON2, LRRC55 and IGFBP1 might be the potential targets for FSGS diagnosis and treatment. Alves et al [449], Ma et al [450], Esposito et al [451], Ai et al [452], Lu et al [453], Bui et al [454], Bao et al [455], Zuo et al [456], Su et al [457], Wang et al [458], Šalamon et al [459], Harskamp et al [460], Ćwiklińska et al [461], Hishida et al [462], Gunay-Aygun et al [463], Simpson et al [464], Svenningsen et al [465], Wang et al [466], Rao et al [467], Chen et al [468], Bedin et al [469], Cao et al [470], Jang et al [471], de Frutos et al [472], Lin et al [473], Wang et al [474], Liu and Zhuang [475], Feng et al [476], Yu et al [477], Reed et al [478], Milillo et al [479], Chen et al [74], Chen et al [480], Vieira et al [76], Charlton et al [481], Shi et al [482], Zhou et al [483], Nazari et al [241], Greene et al [484], Lazar et al [485], Jobst-Schwan et al [486], Du et al [487], Liu et al [85], Liu et al [488], Liu et al [489], Lee et al [490], Wen et al [491], Sakai et al [492], Zhang et al [493], Wang et al [494], CD5L Castelblanco et al [251], Chen et al [495], Swanson et al [496], Ksiazek, [497], Li et al [498], Zhang et al [499], Fernandes et al [500], Bajwa et al [501], Kisic et al [502], Itani et al [176], Liu et al [503], Chan et al [504], Harrison et al [505], Zhao et al [506], Courbon et al [507], Lin et al [508], Carney, [509], Yan et al [510], Mann et al [511], Lefebvre et al [512], von Vietinghoff and Kurts [513], Menendez-Castro et al [514], Kurata et al [515], Yeo et al [114], Lee et al [516], Huang et al [117], Lee et al [517], Stroo et al [518], Lee et al [519], Feng [520], Qin et al [521], Oliveira et al [522], Yang et al [523], Yahya et al [130], Allegretti et al [524], Díez et al [303] and Park et al [525] found that ALB (albumin), ERRFI1, MTNR1A, EGR1, DUSP1, GATA6, TET2, MAGI2, AFP (alpha fetoprotein), KLF15, GATM (glycine amidinotransferase), EGF (epidermal growth factor), LPL (lipoprotein lipase), PPARGC1A, PKHD1, NPHP3, PLG (plasminogen), NR4A1, AVIL (advillin), STRA6, CUBN (cubilin), OGT (O-linked N-acetylglucosamine (GlcNAc) transferase), RGS2, ILK (integrin linked kinase), RBFOX1, DDIT4, RYK (receptor like tyrosine kinase), ERBB4, PTGER3, KCNK5, SPRY2, TRPC1, KL (klotho), GCLC (glutamate-cysteine ligase catalytic subunit), LRP2, NOX4, RGS1, S100A12, PLEKHH2, SORCS1, ATP6V1C2, FKBP5, CA3, IRF9, ELL2, PFKFB2, ATP6, CCL22, SAA1, CCL24, CD5L, CHGA (chromogranin A), CNR2, DBH (dopamine beta-hydroxylase), TNFSF11, XCL1, ELANE (elastase, neutrophil expressed), IFNG (interferon gamma), MPO (myeloperoxidase), CD70, IRF4, VAV1, LYZ (lysozyme), BTK (Bruton tyrosine kinase), LCN2, EPHB2, TREM2, PLD4, JAK3, KCNN4, CCR5, CX3CR1, IL11, TRPM2, TNF (tumor necrosis factor), SEMA7A, TLR7, CCL8, NOD2, CCR2, CCL5, WNT7A, SIGLEC1, DOK3, NOS3, SIGLEC7, RETN (resistin) and S100B are closely related to the development of other kidney diseases. Özcan et al. [526], Emokpae et al. [527], Batista et al. [528], Darbari et al. [529], ElAlfy et al. [530], Hounkpe et al. [531], Vogel et al. [532], Afifi et al. [533], Shmukler et al. [534], El Sissy et al. [535], Cavalcante et al. [536], Sadler et al. [537], Kalai et al. [538], Silva et al. [539], Jhun et al. [540] and Cai et al. [541] that ALB (albumin), LPL (lipoprotein lipase), KL (klotho), UGT2B7, IFNG (interferon gamma), MPO (myeloperoxidase), BTK (Bruton tyrosine kinase), CD209, KCNN4, CCR5, TNF (tumor necrosis factor), CCR2, CCL5, NOS3, S100B and HBB (hemoglobin subunit beta) are associated with progression of sickle cell disease. Studies showed that altered expression of MTNR1A [542], GATA6 [543], EGF (epidermal growth factor) [544], LPL (lipoprotein lipase) [545], PPARGC1A [546], ERBB4 [547], KL (klotho) [548], GCLC (glutamate-cysteine ligase catalytic subunit) [549], NOX4 [550], SORCS1 [551], FKBP5 [552], CCNL1 [553], USP25 [554], SAA1 [555], MPO (myeloperoxidase) [556], GATA1 [557], LCN2 [558], IL1RN [559], IL11 [560], PDCD1 [561], TNF (tumor necrosis factor) [562], TNFRSF1B [563], APLNR (apelin receptor) [564], COMP (cartilage oligomeric matrix protein) [565], RETN (resistin) [566] and IGFBP1 [567] were associated with the progression of polycystic ovarian syndrome. Combined with the results of GO and REACTOME pathway enrichment analysis, these results imply these genes might participate in FSGS.

PPI network was constructed via the HIPPIE online database and Cytoscape software. Then, vital up and down regulated genes were screened from the modules by PEWCC1 analysis. Hao et al. [568] demonstrated that altered expression of PLK1 was found to be substantially related to diabetes mellitus. Pal-Ghosh et al [569] reported that altered expression of PLK1 could be an index for hypertension. Altered expression of PLK1 [570] is associated with progression of cardiovascular diseases. Considering that the conversion of the biological functions of normal cells is fundamental to the pathology of FSGS, we infer that DDX5, MATR3, FOS, LUC7L, CLK1, MKI67, LCK, GTSE1 and DOK2 might take part in the progression of FSGS. Therefore, these molecular markers might be used as potential effective candidates for early diagnosis or prognosis of FSGS.

MiRNA-hub gene regulatory network and TF-hub gene regulatory network analyses were performed to better understand the interactions of the hub genes with miRNA and TF. Previous studies have reported that hsa-mir-1231 [571], STAT3 [572], STAT1 [573], GATA2 [574] and GATA3 [575] are related to cardiovascular diseases. Hsa-mir-21-5p [576] and PAX2 [577] are believed to be associated with FSGS. Liu et al [578], Park et al [579], Yang et al [580], Yu et al [581] and Forero-Delgadillo et al [582] mentioned that hsa-mir-21-5p, STAT3, STAT1, GATA2 and PAX2 were related to other kidney diseases. Studies have indicated that hsa-mir-21-5p [583], STAT3 [584], STAT1 [585] and GATA2 [586] plays a substantial role in hypertension. Ortiz-Dosal et al [587], Su et al [588] and Cox et al [589] demonstrated that the altered expression of hsa-mir-21-5p, STAT3 and STAT1 are associated with progression of obesity. Altered expression and activity of STAT3 [590], STAT1 [591] and GATA3 [592] have been demonstrated in diabetes mellitus. As previously reported, EZH2 [593] is altered expressed in polycystic ovarian syndrome. STAT3 [594], STAT1 [594] and GATA3 [595] were a diagnostic markers of multiple sclerosis and could be used as therapeutic targets. The findings from the present study indicated that the DDX17, DDX39B, HNRNPDL, LMNB1, TFAP4, hsa-mir-503-5p, hsa-mir-500b-5p, hsa-mir-129-5p, hsa-mir-6079, hsa-mir-4740-5p, hsa-mir-190a-3p, hsa-mir-1304-3p, hsa-mir-520e, FOXD1, HOXA5, FOXF2, JUN and REL might be a new biomarkers for FSGS.

In conclusion, we identified several altered expressed genes in FSGS that might involve in the pathogenesis of FSGS. Our investigation provides a comprehensive bioinformatics analysis of DEGs in FSGS, helps to understand the underlying molecular mechanisms of FSGS, and might provide key molecular markers and therapeutic targets for FSGS. Further experiments are required to confirm the expression and key functions of the identified genes in FSGS.

## Acknowledgement

I thank Sean Eddy, University of Michigan, Ann Arbor, MI, USA, very much, the author who deposited their NGS dataset GSE197307, into the public GEO database.

## Ethical approval

This article does not contain any studies with human participants or animals performed by any of the authors.

## Informed consent

No informed consent because this study does not contain human or animals participants.

## Availability of data and materials

The datasets supporting the conclusions of this article are available in the GEO (Gene Expression Omnibus) (https://www.ncbi.nlm.nih.gov/geo/) repository. [(GSE197307) https://www.ncbi.nlm.nih.gov/geo/query/acc.cgi?acc=GSE197307)]

## Consent for publication

Not applicable.

## Author Contributions

B. V. - Writing original draft, and review and editing

C. V. - Software and investigation

